# “Multiplex” rheostat positions cluster around allosterically critical regions of the lactose repressor protein

**DOI:** 10.1101/2020.11.17.386979

**Authors:** Leonidas E. Bantis, Daniel J. Parente, Aron W. Fenton, Liskin Swint-Kruse

## Abstract

Amino acid variation at “rheostat” positions provides opportunity to modulate various aspects of protein function – such as binding affinity or allosteric coupling – across a wide range. Previously a subclass of “multiplex” rheostat positions was identified at which substitutions simultaneously modulated more than one functional parameter. Using the Miller laboratory’s dataset of ∼4000 variants of lactose repressor protein (LacI), we compared the structural properties of multiplex rheostat positions with (i) “single” rheostat positions that modulate only one functional parameter, (ii) “toggle” positions that follow textbook substitution rules, and (iii) “neutral” positions that tolerate any substitution without changing function. The combined rheostat classes comprised >40% of LacI positions, more than either toggle or neutral positions. Single rheostat positions were broadly distributed over the structure. Multiplex rheostat positions structurally overlapped with positions involved in allosteric regulation. When their phenotypic outcomes were interpreted within a thermodynamic framework, functional changes at multiplex positions were uncorrelated. This suggests that substitutions lead to complex changes in the underlying molecular biophysics. Bivariable and multivariable analyses of evolutionary signals within multiple sequence alignments could not differentiate single and multiplex rheostat positions. Phylogenetic analyses – such as ConSurf – could distinguish rheostats from toggle and neutral positions. Multivariable analyses could also identify a subset of neutral positions with high probability. Taken together, these results suggest that detailed understanding of the underlying molecular biophysics, likely including protein dynamics, will be required to discriminate single and multiplex rheostat positions from each other and to predict substitution outcomes at these sites.

---

The marriage of protein biochemistry and protein evolution to study the contributions of nonconserved amino acid positions to functional variation (*e.g*. (1)) is a relatively young field. Many of these studies have relied upon exchanging amino acids between homologs or between actively evolving proteins (*e.g*. (2-6)) and upon ancestral reconstruction from sequence alignments (*e.g*. (7-13)). As an alternative approach, we agnostically, but systematically, substituted nonconserved positions with a wide variety of amino acids. This strategy revealed a class of functionally-important positions with substitution outcomes that differed strikingly from textbook expectations (Supplementary Figure 1; (14-20)). For this new class of “rheostat” positions, a key feature is their ability to modulate function (*e.g*. binding affinity, allosteric coupling, or ligand transport) when substituted. That is, when different amino acids are substituted at individual rheostat positions, their resulting functional values lie on a continuum: ranging from better than wild-type, to intermediate values, to completely nonfunctional (Supplementary Figure 1C). As such, changes at rheostat positions provide opportunity to “dial” function up or down by large or small amounts, depending on which amino acid is substituted.

To be classified as a rheostat position, amino acid substitution need only modulate a single functional parameter. However, multiple biochemical parameters are necessary to fully describe the functions of many proteins. For example, in human liver pyruvate kinase, substrate binding is regulated by both an allosteric activator and an allosteric inhibitor, and five biochemical parameters are frequently monitored to assess binding and allosteric regulation (15, 20-23). This allowed us to detect a subclass of “multiplex” rheostat positions, for which substitution simultaneously modulated multiple functional parameters in pyruvate kinase (15, 20). Notably, changes in the affected parameters did not correlate as would occur if, for example, the binding affinity for allosteric effector correlated with the magnitude of allosteric response. Furthermore, the pyruvate kinase multiplex rheostat positions were located around its two allosteric binding sites.

Thus, we wondered whether the underlying molecular mechanisms of rheostatic modulation of function were related to the mechanisms underlying allosteric regulation of function. Determining this is a daunting task, since it remains very challenging to dissect molecular mechanisms of allosteric regulation. As a starting point, we reasoned that a whole-protein assessment of rheostat positions could reveal whether regions involved in allosteric regulation are enriched for either multiplex or “single” rheostat positions (those affecting only one functional parameter). Alternatively, either multiplex or single rheostat positions could be evenly spaced throughout a protein. Experimentally, it is an equally daunting task to measure multiple functional parameters in a whole-protein, comprehensive substitution study. Fortunately, a unique, whole-protein substitution dataset is available for the allosterically-regulated *E. coli* lactose repressor protein (LacI) (24, 25).

LacI is a member of the LacI/GalR transcription repressor family, which comprises orthologs and paralogs that regulate many aspects of bacterial metabolism. To regulate transcription, LacI binds both to DNA “operator” sequences and to small molecule allosteric sugars (reviewed in (26); Figure 1A). *In vivo*, LacI-DNA binding inhibits RNA polymerase transcription of downstream genes (“repression”). When sugars bind at the LacI allosteric site (Figure 1A, gray spheres), the affinity of LacI for DNA operator is modified. The well-known effect of the allosteric “inducers” allolactose and isopropyl β-D-1-thiogalactopyranoside (IPTG) is to weaken affinity for the DNA operator and thereby alleviate repression; this process is called “induction” (27, 28).

**Figure 1.**
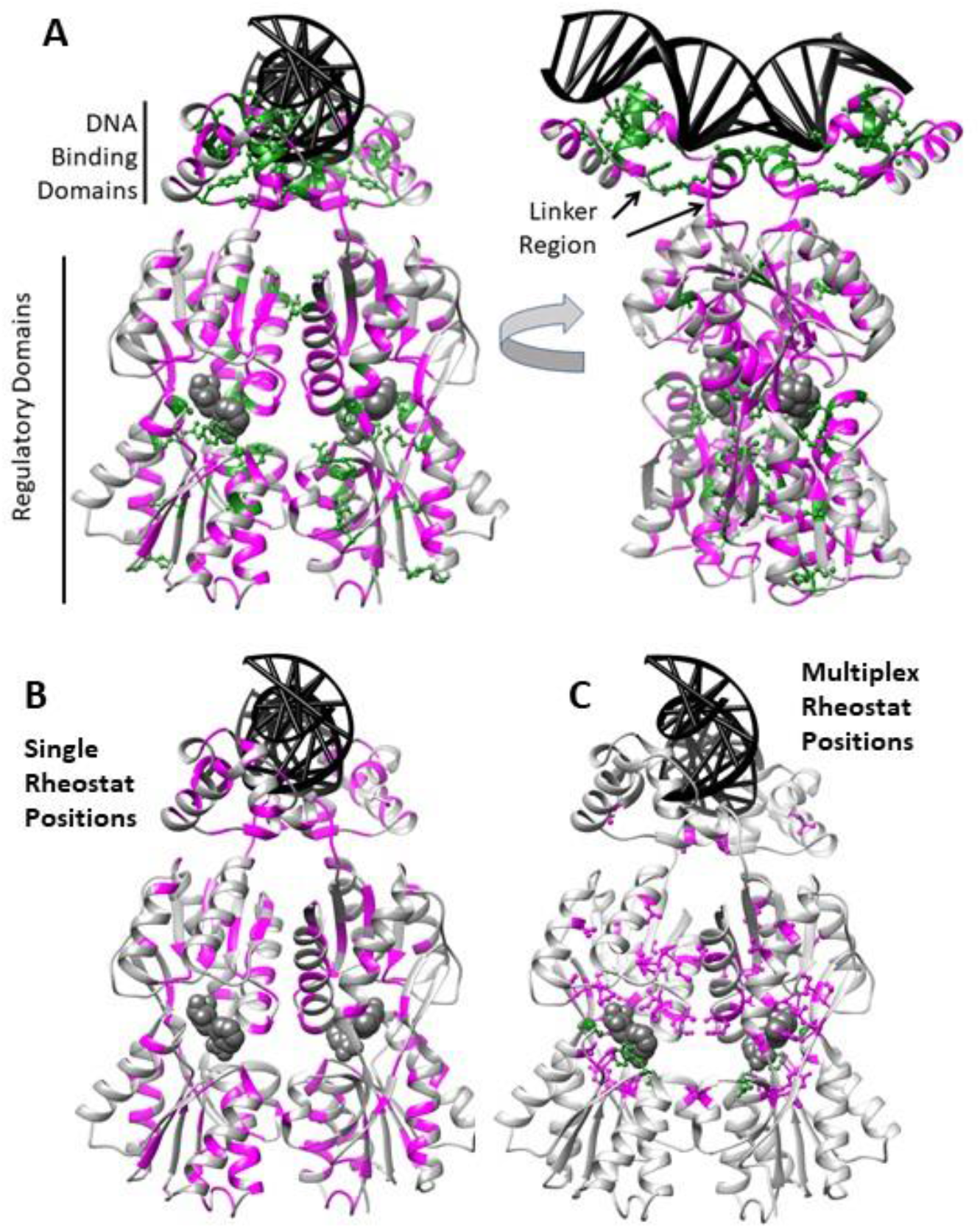
Rheostat (magenta ribbon) and toggle (green ball and stick) positions mapped onto the structure of the LacI homodimer bound to one DNA operator (black ladder) and two sugars in the allosteric binding site (gray spacefilling) (pdb 1EFA; (104)). (Note that the sugars in this structure are “anti-inducers” that enhance DNA binding affinity but bind to the same site as IPTG (27, 28).) The LacI homodimer is the minimal functional unit for both repression and induction; wild-type LacI also has a C-terminal helix at the bottom of the regulatory domain that facilitates formation of a dimer-of-dimers (reviewed in (34)). The Miller data used in the current study were determined using tetrameric LacI, but the tetramerization domains were not substituted. (A) All rheostat and toggle positions are mapped to show the extensive coverage of the protein structure as well as the locations of the DNA binding and regulatory domains. The beginning and end of the linker region are shown with black arrows. Note that toggle positions (green ball-and-stick) are enriched near the two binding sites. (B) The locations of single rheostat positions that modulate either repression or induction are distributed across the protein. (C) The locations of multiplex rheostat positions co-localized with allosteric regions. In this panel, magenta ball-and-stick shows the locations for the subset of double rheostat positions, which modulate both repression and induction, and green ball-and stick shows positions that are both rheostat for repression and toggle for induction.

The regions important to LacI allosteric regulation and communication have been identified by a broad range of structural, functional, and computational studies (*e.g*. (29-44)). These allosteric regions include many positions that are nonconserved in the broader LacI/GalR family (45-47). Indeed, allosteric effector specificity and regulation, along with operator DNA specificity, have evolved functional variation among the LacI/GalR paralogs (34, 48, 49), which could involve amino acid changes at rheostat positions. Indeed, one allosteric region (the “linker”, Figure 1) was the region where rheostat positions were first identified (18).

To identify the presence and prevalence of multiplex rheostat positions in other LacI regions, we turned to the seminal study carried out by the laboratory of Jeffery Miller. In this study, repression and ITPG induction phenotypes were measured for 12-13 substitutions at nearly every LacI position (summarized in (24, 25); hereafter referred to as the “Miller data”). These phenotypes have been related to biochemical parameters by numerous studies with purified proteins (summarized in (50)). In addition to comparing single and multiplex rheostat positions in the current study, we expanded our view to compare and contrast features of rheostat positions with (i) LacI positions that follow textbook substitution rules (*i.e*., toggle positions, defined in Supplementary Figure 1A-B) and (ii) LacI positions that tolerate any amino acid substitution without changing any of the functional parameters assessed (*i.e*. neutral positions, Supplementary Figure 1D).

We first investigated whether structural regions involved in LacI allosteric regulation were enriched for single or multiplex rheostat positions. In the Miller data, multiplex rheostat positions could manifest in either of two ways: (i) as “double” rheostat positions, for which substitutions rheostatically modulate both repression and induction, or (ii) as “rheostat/toggle” positions, for which substitutions rheostatically modulate repression and abolish induction. (“Toggle/rheostat” positions – that toggle repression and rheostatically modify induction – may also exist but cannot be detected in the Miller data; for induction to be detected, repression must be measurable.) For the multiplex positions, we further interpreted phenotypic data within a thermodynamic framework to reason that the various functional parameters changed by substitutions were uncorrelated. These functional outcomes must therefore arise from complex changes in the molecular biophysics underlying various aspects of LacI function.

We next compared the bioinformatic characteristics of the single and multiplex positions, to determine whether they could be distinguished using evolutionary information encoded in multiple sequence alignments. These analyses also provided opportunity to address rigorously several general questions that frequently arise about rheostat positions, including: “How nonconserved is a rheostat position?” and “Do rheostat positions have strong co-evolutionary signals?”. Finally, although not the original focus of this work, we present encouraging results about ways to further improve predictions of the locations of neutral positions.

## Results

The Miller data, with 12-13 substitutions for nearly every position in LacI, meets the criterion of having at least 10-12 substitutions per position that are required to reliably identify rheostatic, toggle, or neutral substitution character (20). These data are reported as repression and induction phenotypes, each of which is the composite of multiple functional and structural equilibria (described further in Methods and Discussion).

As shown in Figure 2, the repression phenotype is altered by amino acid variants that alter the amounts of either L-O complex (LO_2_ or LI_4_O_2_), whereas the induction phenotype is altered by variants that alter the amounts of any of the L-I complexes (LI_4_O_2_, LI_4_, or LI_4_N_2_). In the current study, the repression and induction phenotype data were separately analyzed with the RheoScale calculator to determine the overall substitution behavior for each position in LacI (Supplementary Table 1). Example calculations are shown in Supplementary Figure 2, individual position assignments are in Supplementary List 1, and results are summarized in Table 1 and Table 2. Substitution outcomes for each position are mapped on the LacI structure in Figure 1 and Supplementary Figures 3-6. Forty percent of the LacI positions rheostatically altered either repression and/or induction (Table 2), which exceeds the numbers of either toggle or neutral positions. This is likely a low estimate, since many unclassified positions may also be rheostat positions (Supplementary Figure 6). Further, neither the low-resolution repression nor induction assays measured instances of “enhanced” function, which has been observed in other high-resolution data sets (*e.g*. (18, 51)).

**Figure 2.**
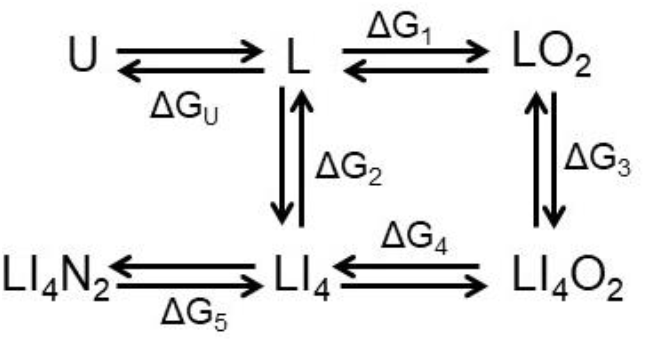
Thermodynamic processes contributing to LacI repression and induction. “L” is folded tetrameric LacI, “O” is operator DNA, “I” is inducer. Each tetramer binds two operators and four inducers. Each of the free energies shown are composite binding events for several binary associations (L binding to two O and/or four I). ΔG_1_ includes the additional binding energy of looping that occurs when a LacI dimer-of-dimers binds two operators on a contiguous piece of DNA. ΔG for dissociation reactions were used to derive Equations 1-4; on this figure, these reactions are shown by the placements of “ΔG” above/below or right/left of the arrows. “U” represents a composite of folded monomer or unfolded LacI, neither of which can bind operator DNA with high affinity. This population would include variants like Y282D, which cause monomeric protein to predominate. Note that, *in vivo*, unfolded protein would likely also be degraded, which would add an irreversible step to this scheme and was omitted for simplicity. Likewise, the current scheme omits folded dimer, which could still bind a single DNA operator; in practice this dimer is unlikely to exist among the variants studied because tetramerization is stronger than dimerization (128). “N” is nonspecific genomic DNA. LacI-inducer can bind to both operator and nonspecific DNA (99, 100, 129). The vast excess of non-specific DNA competes LacI off of the operator DNA. In the absence of inducer, LacI affinity for operator is ∼1000-fold tighter than for non-specific DNA (100). Thus, the complex of un-induced LacI and nonspecific DNA is not shown.

**Table 1.** Numbers of LacI positions assigned to each substitution category for repression and induction phenotypes.

| Category | Repression<sup>a</sup> | Induction<sup>b</sup> |
| --- | --- | --- |
| Neutral | 108 | 171 |
| Rheostat | 109 | 49 |
| Toggle | 30 | 12 |
| Unclassified | 81 | 60 |
| Positions assessed | 328<sup>c</sup> | 292<sup>d</sup> |

<sup>a</sup>Repression rheostat and toggle positions are mapped on the LacI structure in Supplementary Figure 3.
<sup>b</sup>Induction rheostat and toggle positions are mapped on the LacI structure in Supplementary Figure 4.
<sup>c</sup>Miller *et al* did not substitute the C-terminal tetramerization domain or position 1 (24, 25).
<sup>d</sup>Induction cannot be measured for many of the variants in repression category 4 (“I-“ in the original Miller reports). If too few substitutions were available for a position, RheoScale does not calculate scores. For that reason, many repression toggle positions were excluded from analyses of induction phenotypes.

**Table 2.** Numbers of LacI positions with each aggregate substitution behavior.

| Category | Affected Phenotype | Counts | Fraction |
| --- | --- | --- | --- |
| Double Neutral<sup>a</sup> | Neither | 76 | 0.23 |
| Single Rheostat | Either Repression or Induction | 104 | 0.32 |
| Double Rheostat<sup>b</sup> | Both Repression and Induction | 27 | 0.08 |
| Single Toggle | Either Repression or Induction | 38 | 0.12 |
| Rheostat/Toggle<sup>b</sup> | Repression/Induction<sup>c</sup> | 4 | 0.01 |
| Unclassified<sup>d</sup> |  | 83 | 0.25 |
| LacI monomer<sup>e</sup> |  | 329 |  |

<sup>a</sup>Mapped on the LacI structure in Supplementary Figure 5.
<sup>b</sup>The double rheostat positions and the rheostat/toggle positions are two subgroups of multiplex rheostat positions.
<sup>c</sup>For induction to be detected in the Miller study, repression must be measurable; thus, positions that were toggle for repression and rheostat for induction could not be detected. These positions likely exist, but here would be classified with the repression toggle positions.
<sup>d</sup>Mapped on the LacI structure in Supplementary Figure 6.
<sup>e</sup>Not counting the C-terminal tetramerization domain; position 1 was used to determine the fraction of each type of position even though it was not substituted in the Miller study.

### Structural distributions of single and multiplex rheostat positions

Next, the locations of the rheostat and toggle positions were mapped onto the LacI structure. Figure 1 shows the combined repression and induction substitution outcomes. Repression rheostat positions were found in all domains and subdomains (Supplementary Figure 3). The locations of the induction rheostat positions were more limited (Supplementary Figure 4), but this may be an underestimate of their distribution (see Table 1 footnote). As a combined group, rheostat positions were not restricted to types of secondary structures, nor were they confined to either the interior or exterior of the protein (Figure 1). In contrast, toggle positions were concentrated in regions that directly contact bound DNA operator or allosteric sugar, and neutral positions were depleted in the ligand binding sites (Figure 1, Figure 3, and Supplementary Figures 3-5). When the single and multiplex rheostat positions were compared, striking differences were apparent. Single rheostat positions were present in all domains and subdomains, whereas the locations of multiplex rheostat positions clustered in the N-subdomain of the regulatory domains and around the binding sites for the allosteric ligand (Figure 1C). Several multiplex rheostat positions directly contacted ligand.

**Figure 3.**
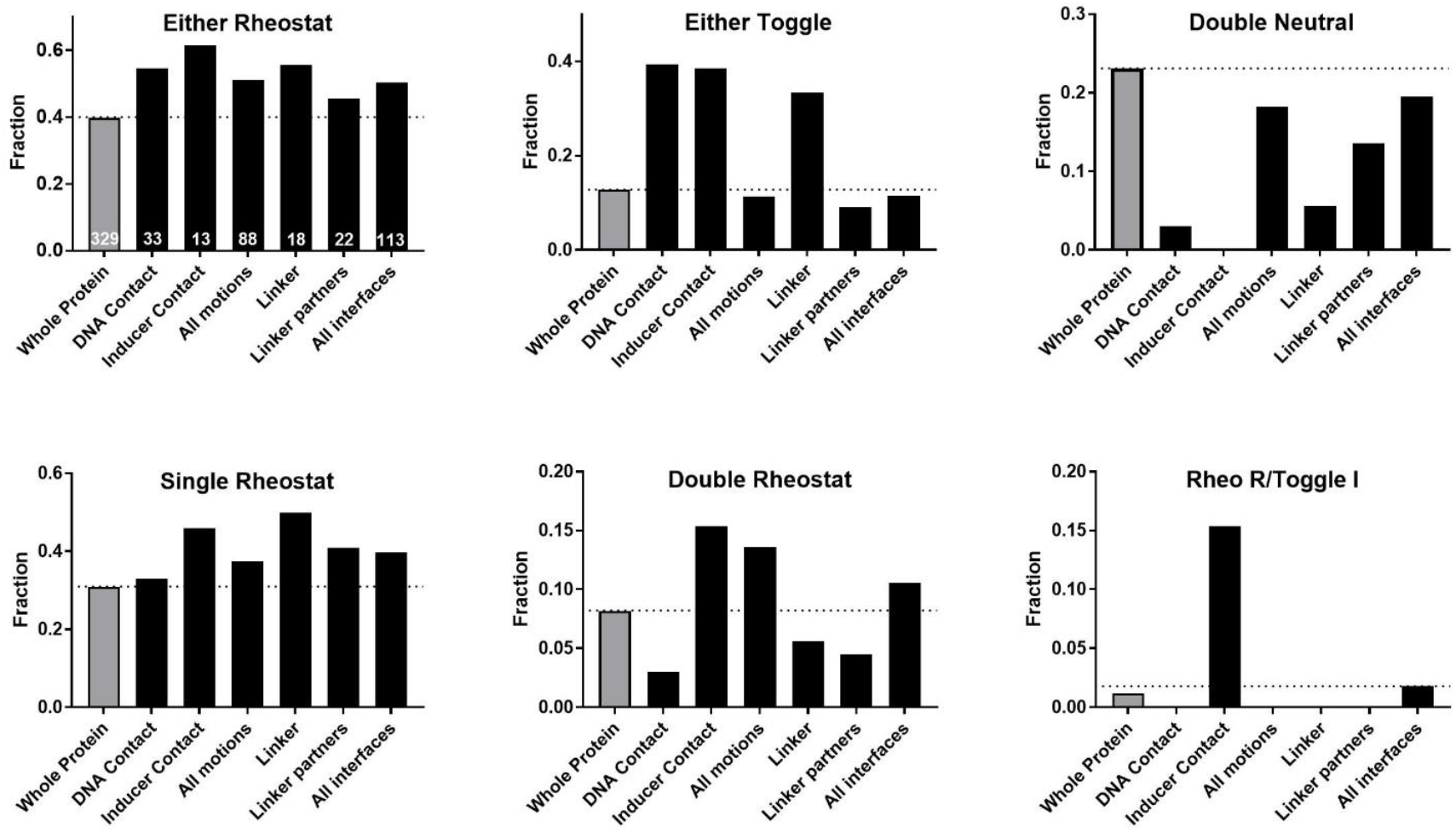
Fraction of positions in each LacI structural region that exhibit various rheostat, toggle, and neutral substitution outcomes. The various LacI regions involved in DNA or inducer binding sites and/or in motions associated with allosteric regulation are listed on the x axes. The total number of positions in each structural region is shown in white on the first panel. Dashed horizontal lines are to enable comparison with the fraction of each type of position in the “Whole Protein” (gray bars). “DNA Contact” and “Inducer Contact” comprise positions that directly contact ligand. “All motions” comprise positions that differ substantially on alternative LacI structural conformations (DNA bound, inducer bound, and apo-protein). The structural location of the “Linker” is shown on Figure 1A. “Linker partners” directly contact positions in the linker region. “All interfaces” comprises all positions that participate in inter-monomer interactions. Positions comprising each region are listed in Supplementary List 2; citations are in Methods; p values are in Supplementary Table 2. The plot for “Either Rheostat” counts all positions with a rheostat designation (both single and multiplex). The subclasses of rheostat categories are shown in the lower panels to highlight the different structural distribution of the double rheostat positions. Likewise, the plot for “Either Toggle” contains both single toggle positions and those that are rheostats for repression/toggle for induction. The latter category is also individually considered in the lower row of panels. Finally, the plot for “Double Neutral” counts positions that are neutral for both phenotypes.

**Figure 4.**
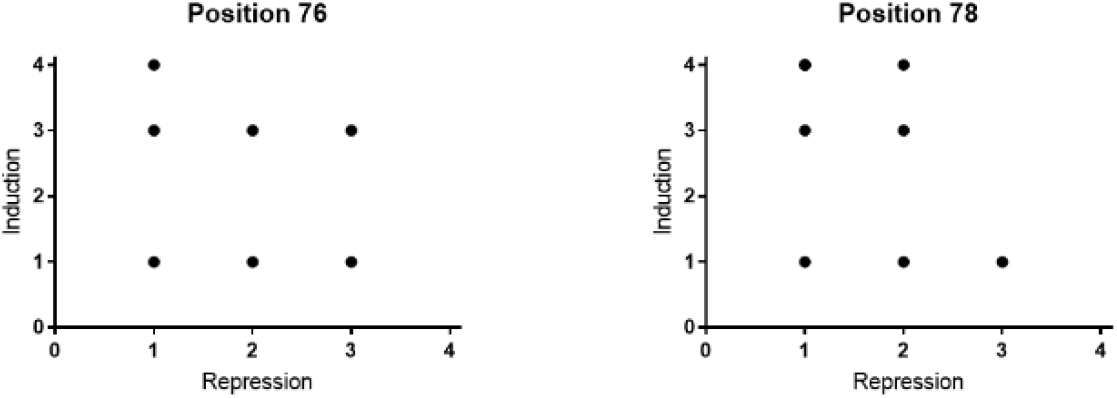
Correlation plots for all substitution phenotypes at double rheostat positions 76 and 78 show no correlation. This suggests that varied substitutions at the same location may differentially affect the contributing functional parameter(s) shown in Figure 2. The raw data for these plots are in Supplementary Table 3. Note that some data are stacked on top of each other in the plots and are thus not visible; other substitutions are missing from the plots because the induction phenotype could not be measured due to weak repression. Plots for other double rheostat positions are shown in Supplementary Figure 10. (Plots for the rheostat/toggle positions would not be informative; since most substitutions have the weakest induction phenotype, all data points would fall at the top of the plot).

**Figure 5.**
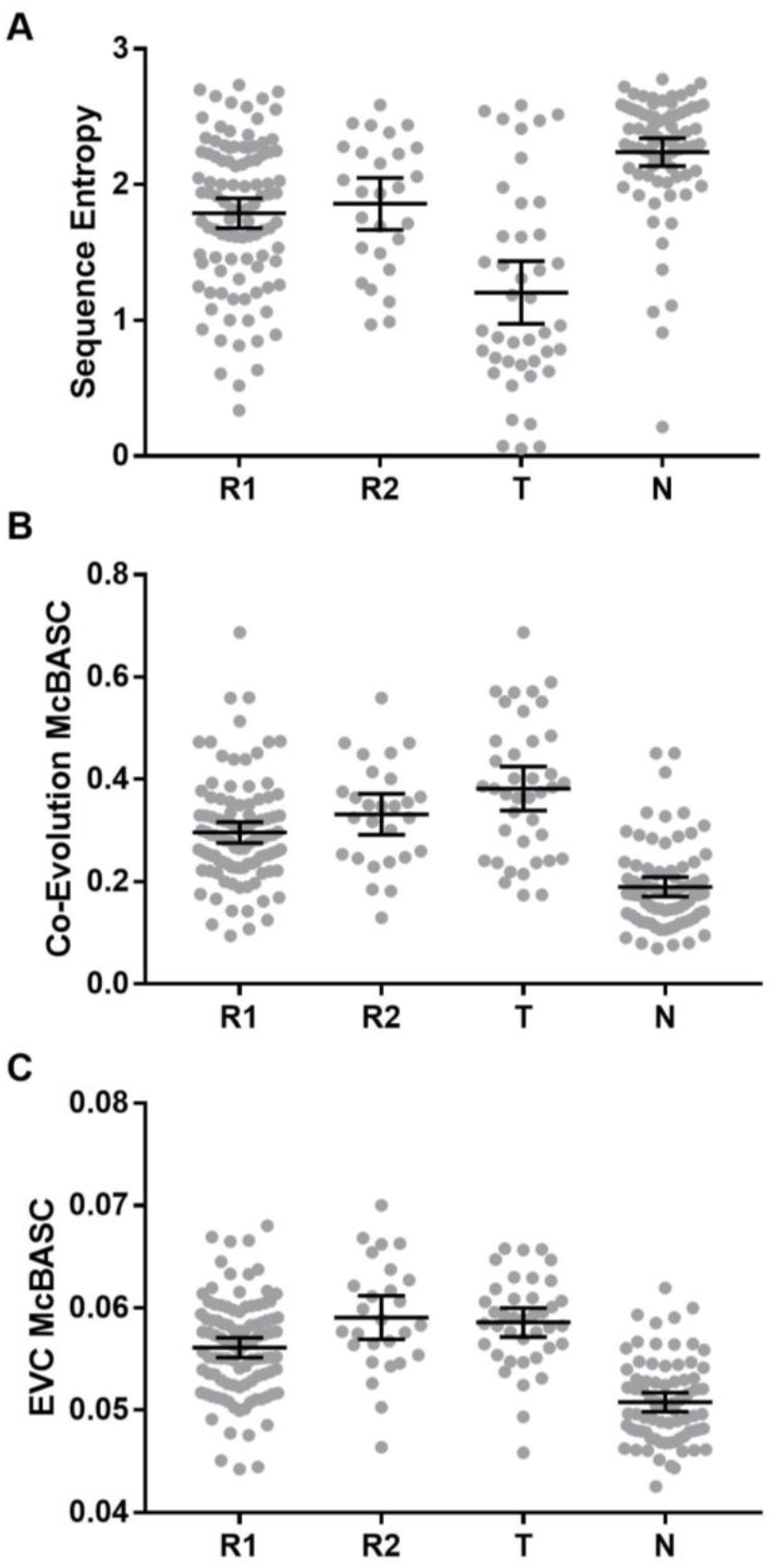
Sequence entropies, co-evolution, and eigenvector centrality scores for LacI functional groups. Various bioinformatic scores were determined for each LacI position. Score distributions were compared for the single rheostat positions (either repression or induction, (“R1”), double rheostat positions (“R2”), toggle positions (“T”), and neutral positions (“N”). The black bars show the mean and 95% confidence limits for the mean of each distribution.

To further examine the apparent clustering of the multiplex rheostat positions, we compared the distributions of the substitution outcomes across several LacI regions (Figure 3). The positions included in each region are listed in Supplementary List 2 and further described in Methods. In these comparisons, we considered all rheostat positions as a combined group (“either rheostat” comprises single rheostat positions for either phenotype and all multiplex rheostat positions), as well as the subclasses of single rheostat, double rheostat, rheostat/toggle positions.

Results in Figure 3 show that some regions appear to have a higher (or lower) density of various position classes than the whole protein. The p values for these comparisons are in Supplementary Table 2 and support many of these observations. In particular, the regions implicated in allosteric regulation and ligand binding had more rheostat positions (“either rheostat”) than the whole protein. The rheostat/toggle positions were located exclusively near the inducer binding site, consistent with the observation that many of the other toggle positions were located in binding sites. As expected, these functionally important regions of LacI were depleted of neutral positions. Finally, the positions that directly contact inducer sugars, positions involved in known protein motions, and positions in the monomer-monomer interfaces appear to be enriched for double rheostat positions relative to the whole protein. Although these differences do not have statistically significant p values (Supplementary Table 2), it is possible that significance was masked by the small overall number of double rheostat positions (27, Table 2) along with the small numbers of positions in some of the protein regions.

Among the allosteric regions, the linker region – where rheostat positions were first identified (18) – is of particular note. Since the linker positions are involved in multiple domain interfaces, we also hypothesized that their interface partners might also be rheostat positions. However, the linker interfaces partners were at most modestly enriched for any type of rheostat position relative to the whole protein (Figure 3). Thus, inter-domain interfaces are not necessarily “hot-spots” for rheostat positions, even when the domains contain allosterically linked ligand binding sites.

### Predicted stability changes at single and multiplex rheostat positions

As shown in Figure 2, the phenotypic Miller data are sensitive to changes in either function or protein stability. Likewise, we previously reasoned that substitutions at rheostat positions could affect protein function, protein stability, or both (see Figure 1 in (17)). Indeed, numerous studies have described protein substitutions with intermediate effects on stability (*e.g*. (52-56)), consistent with the presence of rheostat positions that alter protein stability. However, prior studies of LacI linker rheostat positions focused on those that modulated ligand binding (*e.g*. (51, 57, 58)(18)). Nevertheless, for other LacI regions, substitutions could diminish protein stability or alter assembly (summarized as “U” in Figure 2), which would manifest as diminished repression in the Miller data.

Thus, we attempted to separate effects on DNA binding from effects on stability using the FoldX algorithm (59) to predict stability changes. First, for all possible amino acid *substitutions* at every position in the LacI regulatory domain, we compared FoldX scores to repression phenotype (Supplementary Figures 7 and 8). Second, for all *positions* in the regulatory domain, we compared overall functional outcomes (Table 2) to overall stability outcomes that were derived by using RheoScale on FoldX scores (Supplementary Figure 9).

Two sets of control data are available to assess the predicted stability changes: First, neither repression neutral substitutions nor neutral positions (*i.e*., neutral in both repression and induction phenotypes) should experience a drastic loss of stability; otherwise they would not be able to perform repression. Second, positions with a large detrimental effect on stability should be assigned to the repression toggle category. Only the second criterion was met in FoldX scores. Many neutral substitutions and positions were incorrectly predicted to have large effects on stability, which leaves us reluctant to draw strong conclusions from these calculations. It is nevertheless interesting that substitutions at many of the single and multiplex rheostat positions were not predicted to have large effects on LacI stability. This is consistent with the finding of Bromberg and Rost, who noted that “many function changing mutations have no effect on stability” (60).

### Multiplex rheostat positions alter multiple functional parameters

We next considered whether the multiplex rheostat positions were obligate allosteric positions. That is, we attempted to ascertain whether the dual effects on repression and induction phenotypes arose solely from altering allosteric communication between the two binding sites. In theory, this does *not* have to be the case. A substitution can alter ligand binding without altering allosteric regulation (15, 22, 61-63). To illustrate this, a simplified thermodynamic scheme for LacI binding operator, inducer, and nonspecific genomic DNA is shown in Figure 2. If a substitution simultaneously altered DNA binding in the absence (ΔG_1_) and presence (ΔG_4_) of inducer in the same direction and magnitude (or altered inducer binding in the absence/presence of DNA), then the magnitude of allosteric response would not be altered. For a position that only altered binding, plots for all variants’ ΔG_1_ versus ΔG_4_ would correlate.

However, the Miller dataset can only be plotted as repression versus induction phenotypes. As detailed in the Methods and shown in Figure 2, these phenotypes comprise multiple events that can be represented with equations 1-4:

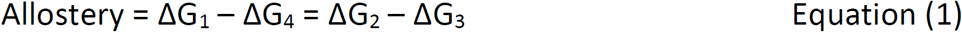

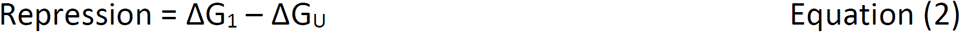

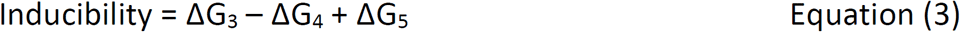

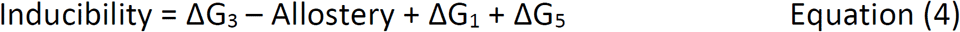

For the allosteric protein regions containing multiplex rheostat positions, we can make one simplification in equations 3 and 4: ΔG_5_ (non-specific binding) is unlikely to be altered by substitutions at these positions, because it is mediated by electrostatic interactions with the DNA binding domain (64). Even so, data for the induction phenotype does not discriminate changes in ΔG_3_ (inducer binding) from changes in allostery. Further, for the repression phenotype, we showed changes in ΔG_U_ cannot be reliably parsed from those in operator DNA binding (ΔG_1_) (Supplementary Figure 7 and Supplementary Figure 9). Thus, we cannot use the current data to ascertain whether the LacI multiplex rheostat positions are indeed allosteric positions. Similar limitations impair interpretation of “deep mutational scanning” experiments (67, 68)^1^ that having growing popularity for assessing substitution outcomes in other proteins.

Nevertheless, information can be gained by plotting induction versus repression for all substitutions at each double rheostat position.Representative plots are shown in Figure 4; the raw data is in Supplementary Table 3 and all plots are shown in Supplementary Figure 10. If all effects on the phenotypes were mediated by allosteric changes or any other single parameter, these plots should show a correlation. However, they instead show a striking *lack* of correlation. This is best explained if the individual substitutions differentially alter the functional parameters comprising each phenotype. This finding is consistent with the behaviors of multiplex rheostat position in pyruvate kinase, which also lacked correlation between five different binding and allosteric parameters (15).

### Bioinformatics characteristics of single and multiplex rheostat positions

As noted in the Introduction, we first identified the class of rheostat positions when we initiated studies of evolutionarily nonconserved positions. One of these studies suggested that nonconserved positions with strong phenotypic signals (defined further below) are indicative of rheostat positions (18). The Miller data provide additional opportunities to (i) test the general applicability of this and other bioinformatic properties across a whole protein and (ii) determine whether differences exist between single and multiplex positions.

“Nonconservation” is a generic term that encompasses a wide variety of amino acid changes. “Sequence entropy” is a measure of sequence conservation that incorporates both the range and frequency of amino acid change during evolution. In addition, at least three other patterns of change have been observed in sequence alignments for nonconserved positions: (i) positions with pairwise co-evolution, (ii) positions constrained by “interactions” with multiple positions (“eigen-vector centrality”; note that “interactions” are not necessarily direct structural contacts, as discussed in (45, 46)), and (iii) positions for which amino acid changes coincide with major branches in the protein family’s phylogenetic tree. Brief details about the representative analyses used to detect these patterns of change in the LacI/GalR sequences alignment are included in Methods. All analyses in this study utilized a curated sequence alignment previously generated for the LacI/GalR family that samples 34 major subfamilies of ortho- and paralogs (45-47).

The original rheostat positions identified in the LacI linker were nonconserved with strong phylogeny scores and poor co-evolution scores (18, 47). We hypothesized that these might be generalizable evolutionary features of rheostat positions, and we further wondered whether multiplex rheostat positions (specifically the double rheostat positions) would show a distinctive signal. (We reasoned that the “toggle” outcome of the four rheostat/toggle positions might dominate an evolutionary outcome; thus, these positions were grouped with the other toggle positions.) In the following comparisons, the substitution groups were designated as: single rheostat positions (“R1”), double rheostat positions (“R2”), toggle positions (“T”), and neutral positions (“N”). For these groups, we compared, contrasted, and combined the scores from 17 different sequences analyses (Figure 5, Figure 6, and Supplementary Figure 11). We next assessed whether thresholds in the bioinformatic scores could be identified to separate the substitution categories (Table 3, Supplementary Tables 4 and 5). Since statistical comparisons among four classes are very difficult to interpret (further discussed in Methods), we performed a three-group analysis by combining the R1 and R2 categories into the general class of rheostat positions (“R”) and compared them to positions in the T and N categories.

**Figure 6.**
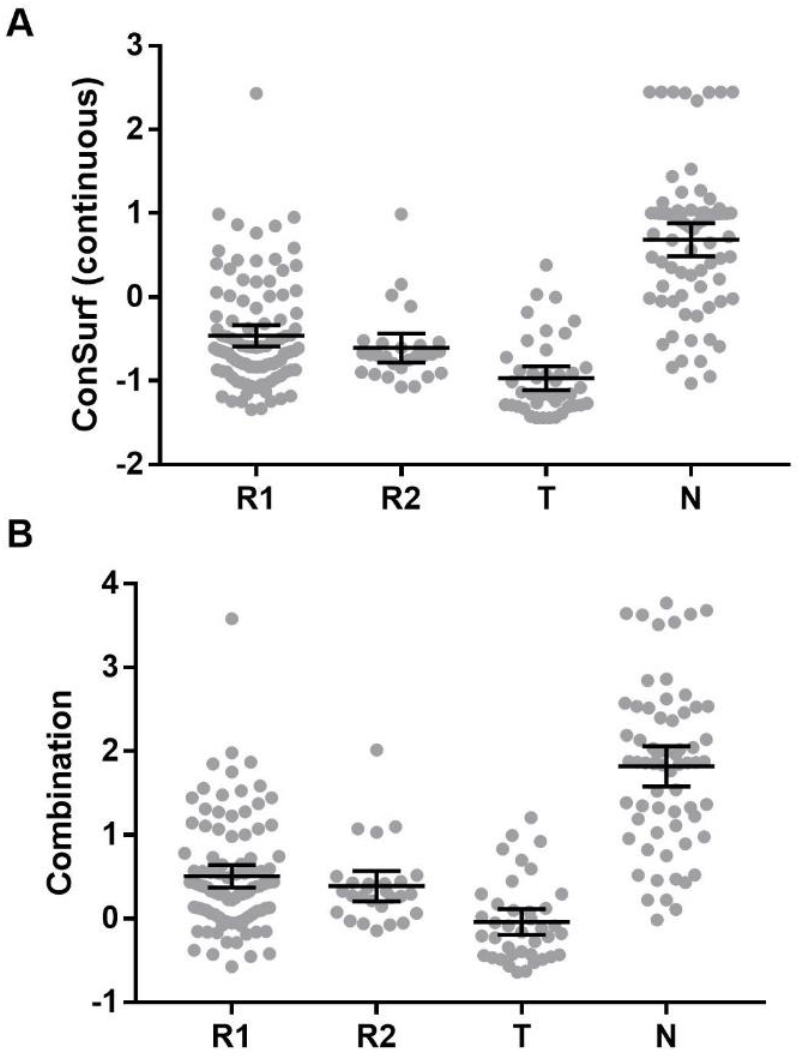
Phylogenetic and combined scores for LacI functional groups. (A) ConSurf calculations derive analog (continuous) scores for each position (shown here) and then discretize these scores into nine categories for its final presentation (not shown). The distributions of both continuous and discrete scores for the LacI rheostat, toggle, and neutral positions were examined in statistical analyses and showed some distinctions (Table 3). The distribution of discrete scores better identified neutral positions, whereas the distributions of continuous scores better separated toggle and rheostat positions. (B) Combination scores were calculated for each LacI position from the analyses listed in Supplementary Table 6.

**Table 3.** Statistical analyses of various algorithms’ abilities to discriminate N, R, and T positions in LacI^a^

| Analysis | TC (T)<sup>b</sup> | TC (R) | TC (N) | FC R\|T<sup>c</sup> | FC N\|T | FC T\|R | FC N\|R | FC T\|N | FC R\|N | Thresholds:C1,<br>C2 |
| --- | --- | --- | --- | --- | --- | --- | --- | --- | --- | --- |
| ConSurf (continuous) | 0.6323 | 0.5566 | 0.8276 | 0.2794 | 0.0882 | 0.2082 | 0.2351 | 0.0299 | 0.1425 | -1.008, 0.1661 |
| ConSurf (discrete)<sup>d</sup> | 0.7985 | 0.5421 | 0.6964 | 0.1657 | 0.0357 | 0.2157 | 0.2421 | 0.0619 | 0.2417 | 4.6407, 7.8375 |
| Combination | 0.6351 | 0.5518 | 0.7981 | 0.3024 | 0.0625 | 0.2316 | 0.2166 | 0.1672 | 0.0348 | 1.1381, 2.1954 |
| funTRP | 0.7381 | 0.3307 | 0.6842 | 0.1667 | 0.0952 | 0.4016 | 0.2677 | 0.0789 | 0.2368 | N/A |

<sup>a</sup>Results for other bioinformatic analyses, along with VUS values, are in Supplementary Tables 4 and 5. VUS analyses are analogous to the area under ROC curves (see Methods).
<sup>b</sup>TC (i): Given that the truth was i, what was the probability that a position was correctly classified as i, with i corresponding to N, R, and T, as indicated. The threshold values used to carry out these calculations are in the last column.
<sup>c</sup>FC (i|j): Given that the truth was j, what was the probability that a position was misclassified as i, with i and j corresponding to N, R, and T, as indicated. The threshold values used to carry out these calculations are in the last column.
<sup>d</sup>The threshold values shown for discrete ConSurf scores were determined after applying kernel estimates. In practice, since the nature of these ConSurf scores is discrete, thresholds are rounded to 4.5 and 7.5.

First, comparison of sequence entropy scores showed that most rheostat positions were indeed nonconserved. However, both single and double rheostat positions exhibited broad ranges of sequence entropies that overlapped with each other (Figure 5A). In agreement with textbook expectations, toggle positions generally exhibited lower sequence entropies and neutral positions generally exhibited higher sequence entropies. However, no clear thresholds separated the different types of positions (Supplementary Table 5). Indeed, numerous toggle positions exhibited high sequence entropies and a few neutral positions exhibited low scores (Figure 5).

Next, we compared the maximum pairwise co-evolution scores. The algorithm with the best discrimination is shown in Figure 5B; results from other analyses are in Supplementary Figure 11. None of the methods showed significant differences among the score distributions for single and double rheostat positions or toggle positions. The distributions of co-evolutionary scores for neutral positions, however, were often lower than those of toggle or rheostat positions. For the “eigenvector centrality” measure of multiple constraints, the McBASC algorithm did show some discrimination for single and double rheostat positions (p = 0.0137) (Figure 5 and Supplementary Figure 11). The neutral positions again exhibited a lower distribution of scores in several of these analyses.

None of the phylogenetic methods discriminated the single and double rheostat positions. However, all of the phylogenetic analyses showed better separation of rheostat (R1+ R2), toggle, and neutral positions (Figure 6, Supplementary Figure 11, Supplementary Tables 4 and 5) than the other analyses, consistent with our prior observations (18). The best performer among the phylogenetic methods was ConSurf (Figure 6; Table 3; (65, 66)).

Since several of the methods weakly separated rheostat (R1+R2), toggle, and neutral positions, we next tested whether multivariable combinations of analyses could provide better discrimination. No approach was identified for distinguishing R1 from R2, but a union set comprising seven analyses showed reasonable separation (Figure 7). Although we did not require the union set to sample all types of analyses, the best set comprised two co-evolutionary methods, two eigenvector centrality methods, and all three phylogeny methods (Supplementary Table 6). However, given Probability of being N the additional information used in this analysis, we were surprised that the combination method was no better than ConSurf alone (Table 3, Supplementary Figure 12).

**Figure 7.**
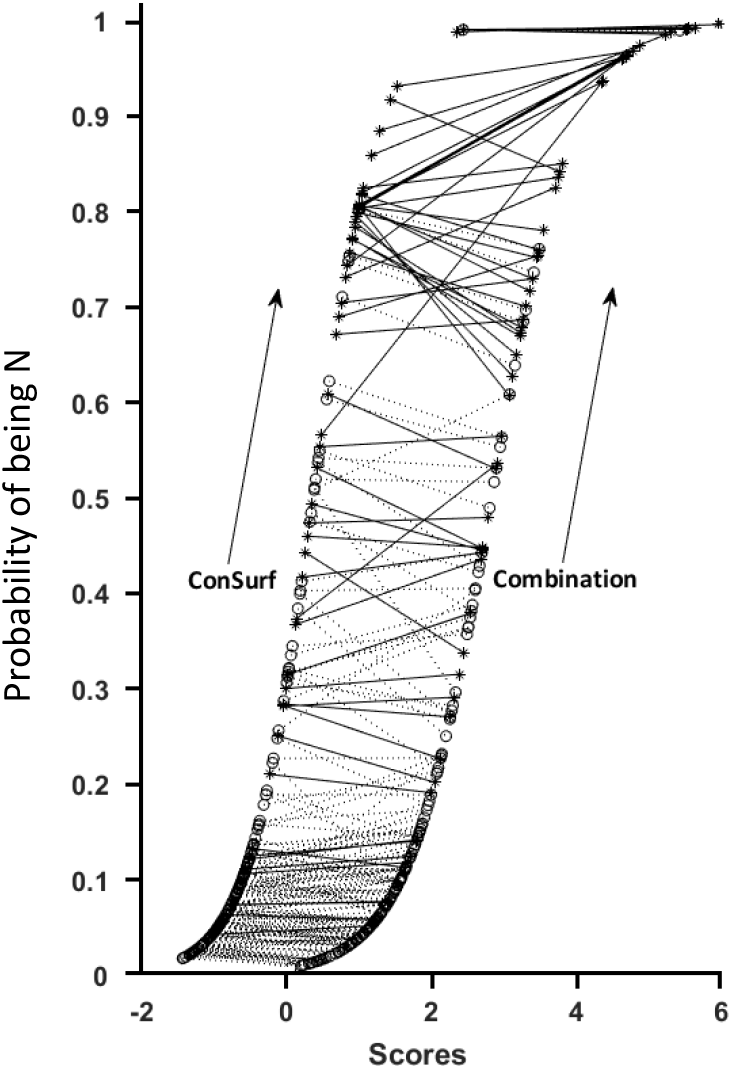
The probability of position neutrality from ConSurf (left sigmoid) and combination (right sigmoid) analyses. Each pair of dots represents one LacI position, and the connecting lines allow ConSurf score/probability to be directly compared to the combination score/probability. The dark symbols and lines correspond to neutral positions; the light symbols and lines correspond to non-neutral positions.

Nevertheless, we noted a persistent trend between ConSurf and the combination method in the distributions of neutral positions relative to the other types of positions (Supplementary Figure 13). Receiver-operator characteristic (ROC) curves from this analysis suggested that ConSurf and the combined method behaved identically (Supplementary Figure 12).

However, when we changed the query from “What fraction of positions are predicted to be in their true category?” to “What is the probability that a certain score correctly predicts neutral substitution behavior?”, the combined method outperformed ConSurf (Figure 7). That is, the combined scores predicted neutrality with high confidence (*e.g*. >0.8) for more positions than did the ConSurf scores. The potential significance of this observation is further considered in the Discussion.

### LacI position predictions with fuNTRp

Since the double rheostat positions clustered in distinct structural regions and the various bioinformatic analyses showed some ability to separate the different classes of positions, we next considered whether combining these features could be useful for their identification. The “fuNTRp” algorithm from the Bromberg laboratory combines structural and sequence alignment features and uses machine learning trained on data from deep mutational scanning studies (69). fuNTRp uses 10 input features: seven are structural, one is genetic, and two are derived from sequence analyses, including the phylogenetic analysis ConSurf that performed well for LacI (Figure 6). In fact, ConSurf was the top contributing feature in their algorithm (69).

We submitted the LacI sequence to the fuNTRp webserver and compared predictions to the rheostat, toggle and neutral behaviors determined from the Miller lab phenotypes (Table 3). fuNTRp was not better than sequence analyses alone at predicting rheostat or toggle positions, and it was comparable for predicting LacI neutral positions. Likewise, predictions for the two rheostat groups were not significantly different: R1 were correctly predicted 31 of 100 times, for a success rate of 0.31 with confidence intervals (0.22-0.40); R2 were correctly predicted 11 out of 27 times, for a success rate of 0.41 with confidence intervals (0.22-0.59). Two differences may contribute to this difference: ConSurf in fuNTRp used an automatically-generated a sequence alignment for LacI, whereas our studies used a highly curated sequence alignment (45-47). fuNTRp may also use a newer version of ConSurf (70) than was used in our studies (71).

## Discussion

Although LacI has a high overall density of rheostat positions (≥40%), only ∼20% of rheostat positions showed multiplex substitution outcomes for the repression and induction phenotypes. This whole-protein study for LacI complements and expands the more limited study of pyruvate kinase positions (15, 20): In both proteins, the multiplex rheostat positions were (i) structurally coincident with allosteric regions and (ii) substitution effects on functional parameters were not correlated. We find these results to be very intriguing and propose that understanding the biophysical underpinnings of rheostat positions can be informative for understanding allosteric regulation, and *vice versa*.

One key consideration for future biophysical studies will be to determine any dynamics changes that arise from substitutions at rheostat positions. Initial dynamics studies were recently completed for rheostat positions in the LacI linker region. In this work, (i) substitutions at multiplex rheostat position 52 showed long-range effects on the dynamics of positions that directly contact DNA, and (ii) the set of linker rheostat positions showed coupling patterns that were distinct from linker toggle and neutral positions (72). Further, these types of dynamics computations have the potential to illuminate epistasis and allosteric regulation (75) and long-distance substitution effects on function (76). Protein dynamics have been shown to change during evolution (*e.g*. (73, 74)). Thus, this may be a general mechanism by which rheostat substitutions lead to functional modulation without grossly distorting the overall protein structure, and it will be imperative to determine whether these dynamics characteristics extends to other multiplex rheostat positions in LacI and other proteins.

When we explored the bioinformatic characteristics of single and double rheostat positions, there was very little to suggest any distinguishing evolutionary properties. These analyses were better at discriminating the combined-rheostat, toggle, and neutral substitution outcomes, but the extensive overlap among their score distributions prevented these bioinformatic properties from being highly predictive for individual positions. This may reflect an inherit limitation of sequence data: Any pattern associated with functional change is likely to be confounded by signals arising from the multiple competing pressures at play during protein evolution (77). In addition, most extant homologs differ at multiple positions, which may have significant non-additivity (“epistasis”) that confounds detection of signals associated with single amino acid changes (*e.g*. (11, 78-84)). Nevertheless, these approaches may be useful for identifying sets of positions that are enriched in each substitution class for further experimental consideration.

Another limiting factor to the use of sequence alignments would be if the rheostat/toggle/neutral character of some positions changed during evolution. Indeed, we previously observed that evolutionarily constrained positions are in different locations on the otherwise common architecture of the LacI/GalR subfamilies (46). Similarly, when we compared rheostat/toggle/neutral substitution outcomes in 10 engineered LacI/GalR homologs (18), the outcomes for analogous positions sometimes varied. For example, position 60 acted as a repression rheostat in four homologs but was repression neutral position in one homolog and weakly rheostatic in others (20). In a second example, position 50 acted as a repression rheostat in four homologs but as a toggle in a four other homologs (20). In a third example, position 52 consistently behaved as a repression rheostat position (20). Thus, some positions appear to change their overall substitution outcomes, but a subset of rheostat positions may be “universal” for the whole family.

Despite these potential limits to the information contained in sequence alignments, the results of this study suggest several possible improvements that could be pursued. First, we should determine what features of ConSurf facilitates its being the best performer for predicting the class of substitution outcomes. Second, although fuNTRp was not as effective as ConSurf alone for identifying LacI rheostat positions, the two analyses used different LacI/GalR sequence alignments. Thus, fuNTRp performance might be improved by allowing the use of curated sequence alignments. Third, the fuNTRp training set included several intrinsically-disordered proteins, whereas LacI is a globular-soluble protein. If globular-soluble, intrinsically-disordered, and integral-membrane proteins have different proportions of rheostat, neutral, and toggle positions, this might erroneously bias fuNTRp predictions. Future implementations of fuNTRp might be improved by using three distinct training sets/analysis pipelines for the different protein types. Experimental studies are underway in example integral membrane and intrinsically disordered proteins to begin exploring this possibility.

Finally, although not the original focus of this study, these LacI results provided another opportunity to assess the success of various methods in identifying positions that are neutral for overall function. This is of general utility for personalized medicine, since variants at neutral positions are unlikely to result in clinically significant pathology. Indeed, the presumed prevalence of neutral positions is a major problem in molecular genetic pathology. Whole genome and whole exome sequencing commonly reveal >10,000 protein-coding variants between individuals (85, 86); thus, extensive bioinformatic filtering is required to separate a single pathological variant from the many neutral (or minimally deleterious) variants detected.

The identification of functionally neutral positions is not as simple as is commonly assumed. For example, in human liver pyruvate kinase, the “common” attributes of neutral positions (high sequence entropy, surface exposure, and insensitivity to alanine substitutions) were not sufficient to identify neutral positions with high confidence. However, a combination score derived from several types of sequence analyses did successfully identify both neutral and near-neutral positions in pyruvate kinase (23). The current work with LacI provides a second, successful demonstration of this general approach, as well as a second example for which high sequence entropy and surface exposure were insufficient to identify neutral positions. Although the full set of LacI neutral positions did not exhibit a distinct threshold that discriminated them from rheostat and toggle positions, a subset of neutral positions was identified with high confidence (Figure 7). The Bromberg lab previously developed the machine learning algorithm, SNAP, to predict which amino acid substitutions are neutral (87, 88) that has performed well in the community experiment CAGI (89). Machine learning algorithms such as SNAP and fuNTRp might be improved by including more types of sequence analyses when predicting the locations of neutral positions and neutral substitutions.

## Conclusion

In conclusion, this work provides a second example in which the locations of multiplex rheostat and allosteric positions showed intriguing overlap. The single and multiplex rheostat positions could not be discriminated by bioinformatic means, although a subset of neutral positions could be identified with high probability. In both LacI and pyruvate kinase, multiplex rheostat positions showed uncorrelated outcomes on the various functional parameters that comprise the protein’s activity, suggesting that changes to the underlying molecular biophysics were complex. These studies further highlight the need to understand the biophysics underlying functional outcomes at single and multiplex rheostat positions before they can be rationally exploited in other systems for protein engineering or personalized medicine.

## Methods

### Interpretation of in vivo phenotypes

The Miller data comprises the allosterically-related repression and induction phenotypes for 12-13 substitutions at nearly every position in tetrameric LacI (24, 25). Data from the low-resolution Miller study are in excellent agreement with quantitative data from a high-resolution *in vivo* repression study for substitutions in the linker region in dimeric LacI, for which levels of folded, functional protein were assessed and found to be unchanged (18). The Miller lab’s *in vivo* phenotypes also show good agreement with extensive *in vitro* biophysical measurements of DNA binding, inducer binding, and/or allosteric response (*e.g*. (35, 39, 41, 51, 90-93) and summarized in (50)). These comparisons showed that repression and induction phenotypes are sensitive to changes in one or more aspects of LacI function, which are summarized in the simplified schematic of Figure 2.

The repression phenotype is accomplished through the formation of “LO_2_” (LacI bound to two operator DNA sequences; Figure 2) and would be altered by changes in any of several steps. First, substitutions that altered binding affinities for DNA operator (ΔG_1_ on Figure 2) would alter repression. Known examples with this behavior include substitutions at position 52, which resulted in to altered DNA binding affinities (42) that highly correlated with a second, high-resolution *in vivo* repression study (18, 57). Second, because a homodimer is required for high affinity DNA binding (*e.g*. (94)), the repression phenotype would be altered by substitutions that changed protein assembly or stability (summarized here as “U” and ΔG_U_). For example, substitutions at position 282 altered the monomer-dimer equilibrium and thereby altered DNA binding (93, 95) and diminished repression (24, 25). Third, diminished repression could arise from an artefact of the amber suppression approach used to create the variants (24, 25), which would be translated as truncated LacI incapable of repression; the prevalence of this occurrence in the Miller data has not been assessed.

Likewise, changes in the inducibility phenotype could arise from multiple functional parameters (Figure 2). To dissect these contributions, it is most convenient to consider the “LO_2_” form that prevents transcription by RNA-polymerase (Figure 2). Starting from this complex, inducibility requires binding inducer ligand (“I”), dissociation of the operator DNA (“O”), and binding to non-specific genomic DNA (“N”). (*In vivo*, folded LacI is always bound to specific or nonspecific DNA and seldom free in the cytoplasm (96)). Thus, the inducibility phenotype could be altered if an amino acid substitution altered inducer binding affinity (ΔG_3_). An example substitution with altered inducer binding affinity is the D149E variant (35). All inducibility assays were carried out at IPTG concentrations that were saturating for wild-type LacI. Nevertheless, the ∼10-fold reduction in D149E inducer binding was scored as the weakest inducibility phenotype in the Miller data (24, 25), which means this assay should be sensitive to smaller changes (and insensitive to larger changes, thereby likely over-estimating toggle characteristics).

The inducibility phenotype would also be altered if a substitution altered allosteric response, that is, the difference in DNA affinities between the conditions of zero and saturating inducer (ΔG_1_ – ΔG_4_; Figure 2). An example substitution with wild-type inducer binding affinity but altered allosteric response is the K84L variant (41, 97, 98); as expected, this variant had an impaired induction phenotype in the Miller dataset (24, 25). A third contribution to inducibility is the competition between the operator and nonspecific genomic DNA for binding to the LacI-inducer complex (ΔG_5_ on Figure 2). Although it has been estimated for wild-type LacI (99-101), nonspecific binding has not yet been assessed for any LacI variant, most likely due to the technical challenges associated with measuring the weak non-specific DNA binding affinity and the fact that different non-specific sequences have slightly different affinities (99, 100).

### Assigning Rheostat, Toggle, and Neutral Substitution Behaviors

Instead of thinking about the role of individual amino acid *side chains* (*i.e*. “residues”), we currently consider the overall role of each *position* within a protein. Such an assessment requires characterizing multiple amino acid variants at each position. Although it would be ideal to have all 20 amino acids at each position, our prior experiences suggest that the overall substitution role of a position can be generally assessed from 10-12 substitutions per position (20). For LacI, the Miller lab used a low-resolution assay to assess outcomes for 12-13 substitutions at most positions in tetrameric LacI (24, 25), which neatly satisfies the 10-12 substitutions needed to classify the rheostatic nature of a position.

To quantify the aggregate substitution behaviors of individual protein positions, we used the RheoScale calculator (20). This calculator uses histogram analyses to assess the toggle-like, rheostatic, and neutral character of each position. “Neutral” scores reflect the fraction of substitutions that are equivalent to wild-type function. “Rheostat” scores reflect the fraction of the total, accessible functional range that was accessed by at least one substitution. “Toggle” scores reflect the fraction of substitutions that are greatly damaging to function.

The RheoScale calculator was designed to be used with quantitative data but can be adapted for qualitative data. In their work with LacI, the Miller lab assigned repression phenotypes to one of four qualitative categories: One category encompassed the tight repressors (including wild-type LacI); two categories of intermediate repression were designated; a fourth category was used for weak or dead repressors. Inducibility phenotypes were likewise assigned to one of four categories, with wild-type again falling in the strongest inducibility category. Thus, following a prior example (72), we assigned these four categories numerical values (1, 2, 3, and 4) and used RheoScale to calculate rheostat, toggle, and neutral scores for each position for each of the two phenotype. Transformed data and histograms for example positions are shown in Supplementary Figure 2.

Once histograms are constructed, RheoScale analyses can employ alternative analytic approaches. In this LacI study, rheostat scores were calculated using the method that gives more weight to bins with intermediate values (20). Calculated rheostat, toggle, and neutral scores for each position’s repression and induction are reported in Supplementary Table 1. Next, we considered significance thresholds for these scores. Since the use of low-resolution experimental data limited histogram analyses to four bins, the thresholds previously established for high-resolution data were inappropriate. Furthermore, and again because of the low-resolution experimental data, we had the most confidence in classifying each position with its dominant substitution outcome rather than trying to assess any intermediate behaviors along the neutral-rheostat-toggle spectrum. These classification results are summarized in Table 1 and Table 2 and listed in the Supplementary List 1. Assignments were made using the following criteria:

First, we identified positions for which all substitutions were in the wild-type-like “strong repression” or “strong induction” categories. This identified which positions were neutral for each of the two phenotypes (Table 1). Note that this is likely an overestimation of truly neutral positions, since the “strong” phenotype categories span a wide range of experimental outcomes (24, 25). For example, the wild-type strong repression bin spans a range that is at least 50-fold, if not larger. Nevertheless, positions for which all substitutions fall in the “strong” bin should *not* be classified as “rheostat” positions: Their variants could not span half of the available functional range, which is the minimum criterion previously used to designate rheostat positions (15, 20).

Second, we identified positions for which more than 75% of the non-wild-type substitutions were in the weak/nonfunctional classification and no more than two substitutions were in the strong category (Table 1). This threshold was slightly more stringent than one used in (16) because it is in better agreement with results from the high-resolution *in vivo* repression study (18). Positions that satisfied this criterion were designated as “toggle” for either repression or induction phenotypes.

For the remaining positions, we designated those with rheostat scores greater than 0.6 as rheostat positions (Table 1). This rheostat score threshold is higher than in previous studies, which used a threshold of 0.5 (20): For high resolution data, two of the ways a rheostat score of 0.5 could be achieved was by (i) half of the substitutions causing intermediate functional outcomes spanning the entire possible range, or (ii) the range of outcomes spanning at least half the possible functional range. However, for the low-resolution phenotype data used herein, a rheostat score of 0.5 might not span half the available range. Thus, we decided to use a more stringent threshold of 0.6. Even with the more stringent threshold, 40% of LacI positions were classified as rheostat positions for at least one of the two phenotypes (Table 2; Figure 1).

Finally, we assessed the composite substitution behavior for each position (Table 2 and Supplementary List 1). Neutral positions must be neutral for both phenotypes (23). The “single rheostat” category was used for positions that only altered one of the two measured phenotypes (the other phenotype was neutral). For multiplex positions that affected both phenotypes, the “double rheostat” category was assigned to positions that rheostatically altered both phenotypes; four positions were identified as being rheostat for repression and toggle for inducer (“rheostat/toggle). Thirty-eight positions exhibited toggle outcomes for one phenotype; if the toggle phenotype was assigned to induction, then repression was neutral (or unclassified); if the toggle phenotype occurred for repression, then effects on induction could not be measured and are thus unknown. Positions with toggle repression/rheostat induction likely exist but could not be detected in these experiments. Finally, 83 positions could not be assigned to a substitution category for either phenotype (Supplementary Figure 6) and thus are not further considered in this work.

### Structural analyses

Amino acid positions at functionally important regions were compiled from a variety of published studies. Citations to various structural studies include (32, 38, 40, 48, 64, 102-108) and key structural features from each study are compiled in the AlloRep database (www.allorep.org ; (50)). A large number of biochemical studies have also been performed by the Matthews’ and other laboratories for LacI variants (*e.g*. (35, 39, 41, 51, 90-93, 109, 110)); a full list of these variants and their functional effects are summarized in the AlloRep database (50).

For this work, we extracted the following positions from the AlloRep database: (i) All positions that participate in the monomer-monomer interfaces between two subunits of any available structure for any available liganded state; (ii) all positions that form the interface between the linker region and the top of the regulatory domain; (iii) all positions that directly contact DNA; (iv) all positions that directly contact inducer IPTG, anti-inducer ONPF (orthonitrophenyl-beta-d-fucopyranoside), or neutral ligand glycerol. In addition, positions identified to be involved in LacI motions or implicated in the allosteric regulation were compiled; these include (iv) positions in the “core pivot” region (39), (v) those identified by targeted molecular dynamics simulations of the transition from the DNA-to IPTG-bound structures (38) and (vi) positions with changes at the N-subdomain and/or that alter allosteric regulation (35, 90, 91, 104, 107). Note that the various lists (Supplementary List 2) have some overlap among their sets of positions.

Structure figures in this manuscript were created using PDB 1EFA of dimeric LacI bound to *lacO*^*sym*^ DNA and anti-inducer ONPF (104); structures were rendered with UCSF Chimera (111). *In silico energetics calculations*

The crystallographic structure 1LBI for the tetrameric lactose repressor core (comprising 4 regulatory domains and 4 tetramerization domains) was downloaded from the Protein Databank (43) and analyzed using FoldX (59). This structure was chosen because it includes the small tetramerization interface and lacks the DNA and effector ligands. The structure was preprocessed using the FoldX “RepairPDB” command, which locally minimizes the force field at “high energy” residues in the structure and those with risk of incorrect rotamer assignment. A FoldX position scan was thereafter conducted for ever residue on every chain, which substitutes each residue to all 19 other residues to determine the change in Gibbs free energy (ΔΔG). Energetic changes for a given position and substitution variant were not identical when different (monomer) chains were mutated. We therefore averaged the ΔΔG across all four chains in the tetrameric structure. The distribution of results in shown in Supplementary Figure 7. The predicted stabilities were analyzed with RheoScale to determine an overall role for each position in protein stability; further details of this calculation and resulting scores are presented in Supplementary Figure 9.

### Bioinformatic score calculations

The sequence alignment for the LacI/GalR family was previously reported and used to generate various evolutionary bioinformatics scores for each amino acid position in the alignment (45-47). This alignment contained 351 representative sequences of LacI/GalR paralogs from 34 subfamilies; sequence identities ranged from 99% to ∼15%. For this work, we used the “whole family” sequence alignment. Preliminary analyses using only sequences containing the “YPAL” linker motif (47) did not exhibit noticeable differences from whole family analyses. Analyses could not be performed on the subfamily of LacI orthologs (sequence identities from 99% to ∼40%) because it contains too few sequences (45); for other proteins, subfamily analyses might provide additional insights (46).

A brief summary of the various types of sequence analyses is as follows:

1. Sequence entropy calculations estimate conservation using an information theoretic approach (Shannon entropy) to quantify the distribution of observed amino acids at each position (112). In addition to quantifying the overall amino acid variability observed at each position, this calculation discriminates between the following two scenarios: When all 20 amino acids are observed at a given position in an alignment with 100 sequences (i) each amino acid could be equally represented (5 occurrences each, perfectly nonconserved) or (ii) one amino acid could occur 81 times, with the other amino acids represented once each (highly – although not perfectly – conserved). Sequence entropy scores are different for these two and other intermediate conditions.
2. Co-evolutionary analyses estimate the extent to which two positions vary together during evolution. For example, if a specific mutation at position A is always correlated with a specific mutation at position B, then positions A and B are said to “co-evolve”. Numerous mathematical frameworks have been developed to quantify co-evolutionary behaviors. As previously reported (and as occurs in other protein families), the scores assigned to any individual pairs of positions do not agree well among different co-evolution algorithms (46, 113), even when additional procedures were used to subtract evolutionary “noise” from the calculations (46). Since no method has been shown to find more “important” positions than any other method, we previously used five, mathematically-divergent co-evolution algorithms for our studies (45, 46): (i) Observed Minus Expected Squared (OMES; (114, 115)), which is based on Chi-squared-like goodness of fit); (ii) Explicit Likelihood of Subset Covariation (ELSC; (116)) and (iii) Statistical Coupling Analysis (SCA; (117)), which are based on a subset perturbation approach; (iv) McLachlan-based Substitution Correlation (McBASC; (118-120)), which is based on coordinated changes within physiochemical classes; and (v) Z-Normalized Mutual Information (ZNMI; (113)), which uses an information theoretic approach. Although several newer co-evolutionary analyses have been developed, the focus of the field has been on improving amino acid contact prediction (reviewed in (121)). We decided not to use these versions because, since many rheostat positions do not contact each other, we reasoned that these algorithms would impose unsuitable constraints. Since co-evolution scores are assigned to pairs of positions, each individual position has *n*-1 co-evolution scores, where *n* is the number of positions (columns) in the sequence alignment being analyzed. This is difficult to map on a structure or to compare to functional outcomes. Thus, for the current analyses, we assigned each position its maximal co-evolution score (maximal edge weight, “MEW”, (46)). We also used MEW scores to generate and assess a composite coevolution scores (45, 46), to determine whether the combination of algorithms could amplify a signal for various classes of LacI positions.
3. Another type of sequence analysis is derived from co-evolutionary methods: “Eigenvector centrality” (EVC) uses a network-based approach identify “central” positions that have the greatest degree of connectivity within a weighted co-evolutionary network (45). These positions can be thought of as being the most constrained overall – by evolutionary “interactions” (not necessarily structural) with several other positions – as opposed to having the highest single constraint from a partner position. Since eigenvector centrality scores were derived from co-evolutionary scores (above), we again generated five sets of EVC scores for the LacI/GalR family along with a composite score generated from all EVC analyses (45).
4. Several analyses incorporate phylogenetic trees in their analyses: “Consurf” (65, 66) uses a phylogenetic tree to estimate the conservation of a position based on its evolutionary rate. “Evolutionary trace analysis” (ETA) also uses a phylogenetic tree to identify positions that diverge earlier in evolutionary history (122, 123). “Two entropies analysis – Objective” (TEA-O) is a third phylogenetic tree-based method that calculates sequence entropy at multiple phylogenetic levels to identify positions that are globally conserved (TEA-O “conserved”) along with those that are non-conserved globally but conserved within subsets of the tree (TEA-O “specificity”) (124).

These various methods were previously used to generate 17 different sets of scores for each of the amino acid positions in LacI (45-47). To demonstrate that different types of analyses highlight different positions, all possible pairs of score sets were plotted against each other (*e.g*. Supplementary Figure 14) and Pearson correlation coefficients were determined (Supplementary Figure 15). As previously noted, the eigenvector centrality analyses showed the best within-class agreement (45).

### Statistical comparisons of experimental results and bioinformatic scores

We divided each of the 17 bioinformatic scores sets into subsets corresponding to the scores of rheostat (R, comprising both single R1 and double R2), toggle (T), and neutral (N) positions. (Since only four LacI rheostat/toggle positions were identified, and reasoning that the toggle outcome would dominate any signal in a sequence alignment, these positions were grouped with the other toggle positions for these analyses.) The distributions of the scores for the four subsets were then compared to determine how well they discriminated the experimental rheostat, toggle, and neutral substitution outcomes. Ideally, these comparative analyses would be performed for the four different categories (single rheostat, double rheostat, toggle, and neutral). However, four-dimensional ROC analyses are difficult to calculate, extremely hard to interpret, and impossible to visualize. Thus, the performance of each score type was evaluated through three-dimensional analysis using ROC surfaces; when possible, the single (R1) and double (R2) rheostat positions were considered separately in downstream analyses.

Three-dimensional analyses of ROC surfaces are analogous to two-dimensional analyses of ROC curves. In the three-dimensional analyses, the volume under the ROC surface (VUS) can yield values within the interval [1/6, 1]. The value of 1/6 corresponds to an uninformative predictor, whereas the value of 1 corresponds to a perfect predictor (125). VUS were determined for all 17 score sets. Confidence intervals for the VUS were derived through the percentile bootstrap resampling method, using 1000 bootstrap samples.

Next, we explored whether a combination of bioinformatic score sets could be identified that had better separation of the classes than any single score set. Since exploring all possible combinations of 17 scores was intractable, we narrowed down the number of scores sets by the following steps. In narrowing down the score sets, we separately considered the R1 and R2 groups in addition to T and N, to allow as many possible signals as might exist. Two highly correlating eigenvector centrality measures (Supplementary Figure 15) were excluded to further simplify analyses. We next applied forward stepwise elimination through logistic regression for all six separate pairs of responses derived from the set (R1, R2, N, T), *i.e*. (R1 *vs* R2), (R1 *vs* N), … and (N *vs* T) (Supplementary Table 6). The selection process was performed with SPSS statistical software. From these results, a final union set of seven bioinformatic score sets was used to create a “combination” score for each LacI position. To generate this score from the component bioinformatic scores, we determined which coefficients of a linear combination maximized the VUS of the ROC surface. The final equation obtained was

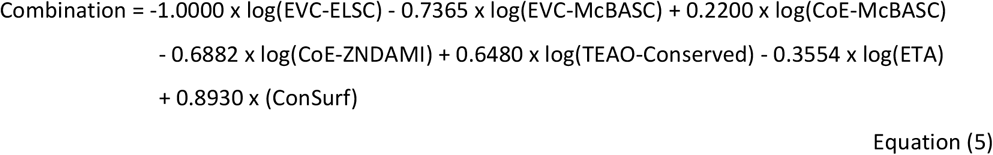

The confidence interval of the VUS corresponding to the combination score was derived through the percentile bootstrap with 1000 bootstrap samples. Note that this combinatorial analysis was derived solely to determine the potential value of combining different types of bioinformatic analyses for predicting the locations of rheostat, toggle, and neutral positions. This combinatorial score was *not* subjected to external validation, nor do we expect this empirical equation to extrapolate to scores sets for other protein families.

For all analyses, we determined the generalized Youden index (126, 127), all three class rates at the Youden based optimal threshold pair of points, and all six false classification rates. For the optimization of the Youden index, we considered kernel-based estimates of the densities of each group that are based on gaussian kernels. The analysis was performed using MATLAB 2019b. In addition, for ConSurf and combinatorial score, we determined pairwise ROC curves that refer to all possible pairs of N, R, T (Supplementary Figure 12).

Finally, another comparison of the ConSurf and combined scores is shown in Figure 7, which plots the probabilities of being neutral (“N”) for each range of scores, as opposed to “not-N”. These probabilities were obtained from logistic regression models using MATLAB R2019 and the equations:

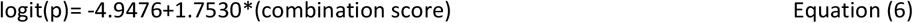

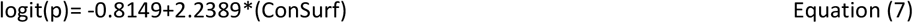

### fuNTR predictions

fuNTRp is a machine learning algorithm that uses structural and bioinformatic information to predict the rheostat, toggle, and neutral substitution outcomes for each position in a protein (69). The seven structural features used by the algorithm include (i) the observed amino acid side chain chemistry, size, and charge and (ii) predictions about each position’s solvent accessibility, secondary structure, residue flexibility, disorder. One included genetic feature was based on the “number of possible nsSNPs (all codons)” (69). The two features derived from sequence analyses included an automatic implementation of ConSurf (which does not use a curated multiple sequence alignment) and the “MSA ratio” (which was defined as the “fractions of residue amino acid per MSA column” (69)). Thus, information from pairwise co-evolution and eigenvector centrality scores were not included in fuNTRp analyses. As reported by Miller et al. (69), each of the ten chosen features contributed different amounts to the final algorithm output. The LacI sequence was submitted to the *fuNTRp* website (https://services.bromberglab.org/funtrp/) to generate predictions about the locations of rheostat, toggle, and neutral positions.

## Data Availability

Data underlying this manuscript and full lists of scores and results are included in the Supplementary Material to this manuscript. The Miller data were published as a Supplement to (Suckow, et al. 1996) and can also be found at www.allorep.org.

## Acknowledgements

This work was supported by the National Institute of General Medicine at the National Institutes of Health (grant numbers GM115340 to AWF, GM118589 to LSK and AWF, and P20GM130423 to LB and LSK as part of the Kansas Institute for Precision Medicine) and by the W.M. Keck Foundation (LSK and AWF). We thank Dr. Kathleen Matthews (Rice University) for discussions about the thermodynamic relationships in Figure 2 and Equations 1-4 and Ms. Kaitlin DeHart (KUMC) for comments on the manuscript.

## Footnotes

1 “Deep mutational scanning” is an experimental design that relies on phenotype competition for high-throughput analyses of variant libraries for the protein of interest. In this approach, the protein variants are expressed in some cellular context that is sensitive to the protein’s function. Using the pool of cells, a variant’s representation in next-generation sequencing is used to infer its prevalence in the cell population, which infers protein phenotype, which is in turn a composite of many functional and structural parameters that could be altered by amino acid substitution (67, 68). That is, variants with high sequence abundance are presumed to have good functionality, whereas variants with low sequence abundance are presumed to have poor functionality. These experiments rely upon only one composite phenotype comprising many thermodynamic parameters and, as of yet, do not detect multiplex rheostat positions.

