## Supplementary for "“Multiplex” rheostat positions cluster around allosterically critical regions of the lactose repressor protein"

Supplement to:

**A whole-protein analysis of the lactose repressor protein reveals that "multiplex" rheostat positions cluster around allosterically critical regions.**

Leonidas E. Bantis[1], Daniel J. Parente[2], Aron W. Fenton[3], Liskin Swint-Kruse[3*]

**Supplementary Figure 1.** Textbook substitution rules and example substitution outcomes at toggle, rheostat, and neutral protein positions.

**Supplementary Table 1.** RheoScale neutral, rheostat, and toggle scores for LacI positions. Due to the size of this table, it is located at the end of this document.

**Supplementary Figure 2.** Substitution phenotype outcomes, resulting histograms, and RheoScale scores for representative LacI positions.

**Supplementary Lists 1.** LacI positions comprising each type of substitution class.

**Supplementary Figure 3.** Locations of repression rheostat positions on the LacI structure.

**Supplementary Figure 4.** Locations of induction rheostat positions on the LacI structure.

**Supplementary Figure 5.** Locations of double neutral positions on the LacI structure.

**Supplementary Figure 6.** Locations of unclassified positions on the LacI structure.

**Supplementary Lists 2.** LacI positions comprising various functional regions.

**Supplementary Table 2.** Statistical comparisons of substitution outcomes among various LacI regions.

**Supplementary Figure 7.** FoldX calculations for the LacI variants as a function of repression phenotype.

**Supplementary Figure 8.** FoldX threshold study and Supplementary Methods.

**Supplementary Figure 9.** RheoScale analyses of position FoldX scores, stratified by functional groups.

**Supplementary Table 3.** Substitution phenotypes at double rheostat positions.

**Supplementary Figure 10.** Phenotype correlation plots for double rheostat positions.

**Supplementary Figure 11.** Distributions of various bioinformatic scores for R1, R2, T, and N functional groups.

**Supplementary Table 4.** VUS for bioinformatic analyses.

**Supplementary Table 5.** ROC analyses and true/false rates of all bioinformatic analyses.

**Supplementary Table 6.** Stepwise logistic regression used to generate the combined analysis.

**Supplementary Figure 12.** ROC for ConSurf and combined scores.

**Supplementary Figure 13.** Binary comparisons (neutral vs non-neutral) of ConSurf and combined scores.

**Supplementary Figure 14.** Correlation matrices for various bioinformatic analyses.

**Supplementary Figure 15.** Pearson correlation coefficients among bioinformatic analyses.

**Supplementary References.**

**Supplementary Figure 1.** Textbook rules (A) and example substitution outcomes for (B) toggle, (C) rheostat, and (D) neutral protein positions. (A) Decades of protein mutagenesis studies have led to several textbook expectations for amino acid substitution outcomes. However, most historical experiments have inadvertently focused on evolutionarily conserved positions and/or relied upon alanine substitutions (*e.g.* (1)), leading to the expectation that substitution outcomes will resemble those shown in panel B. (B-D) These hypothetical examples show idealized substitution outcomes that can arise when multiple amino acid substitutions are made at a single protein position (here, 15 medium-gray bars). The "function" of wild-type protein is shown with light gray bars; small black bars represent functionally "dead" protein. Textbook rules (A) predict the "toggle" substitution outcome (B), in which only amino acid substitutions that are chemically similar to the wild-type amino acid allow function (tall medium-gray bars). Toggle outcomes may be common at evolutionarily conserved positions. (C) At rheostat positions, substitutions show a wide range of functional strengths, including function "better" than wild-type; other noncanonical substitution outcomes observed at rheostat positions are detailed below the figure. (D) Neutral positions tolerate any substitution without altering function.

A short video presentation about rheostat positions is available at:
https://www.youtube.com/watch?v=835ATtoZsko

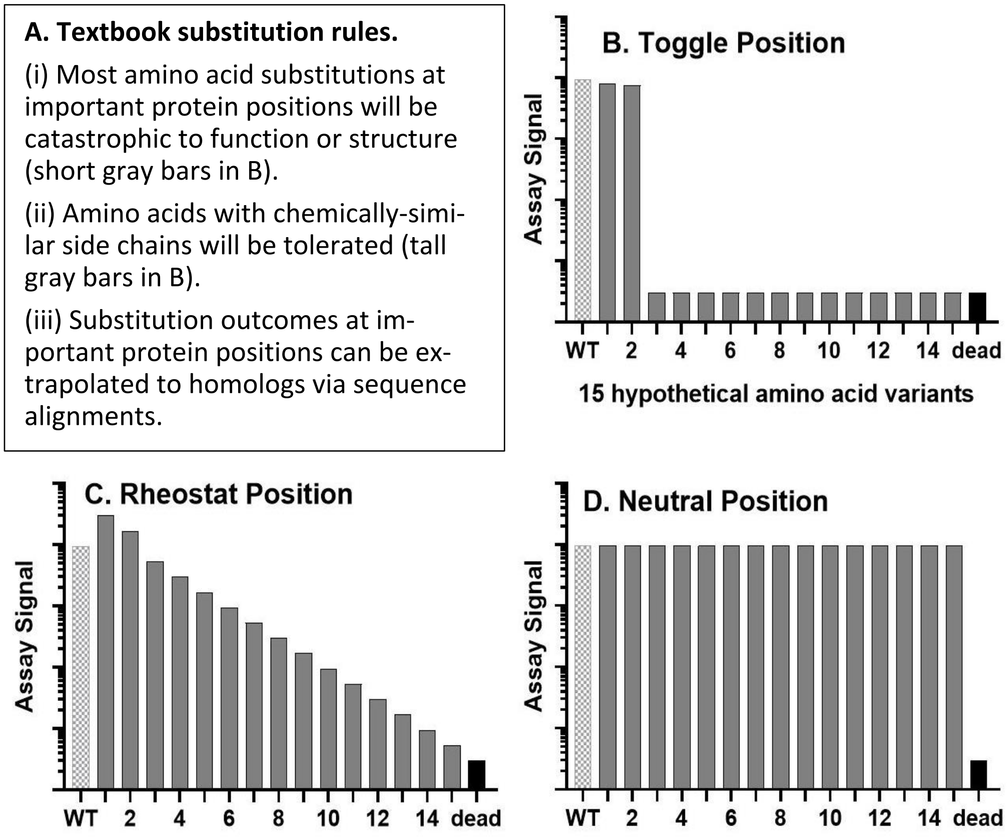


**Rheostat positions do not follow the textbook substitution rules.** Among the experimental studies of rheostat positions cited in the main text, the functionality of specific amino acid substitutions often did *not* correlate with either the general side chain chemistry (*e.g.* hydrophobicity or size) or the observed evolutionary frequency. This is counter to expectations (i) and (ii) in (A). Further, when the same substitution was created at analogous positions in multiple homologs, functional outcomes differed widely, counter to expectation (iii) in (A) (2). Not surprisingly, since these rheostat positions did not follow the textbook assumptions upon which prediction algorithms are built, their substitution outcomes were very poorly predicted by 16 different computational algorithms (3).

**Supplementary Table 1.** RheoScale neutral, rheostat, and toggle scores calculated for each LacI position using the Miller lab data.

Due to the size of this table, it is located as the final pages of this supplement (after the Supplementary References).

**Supplementary Figure 2.** Phenotype outcomes (left), resulting histograms (middle), and RheoScale scores (right) for representative positions in LacI. RheoScale scores for both phenotypes of all positions are in Supplementary Table 1.

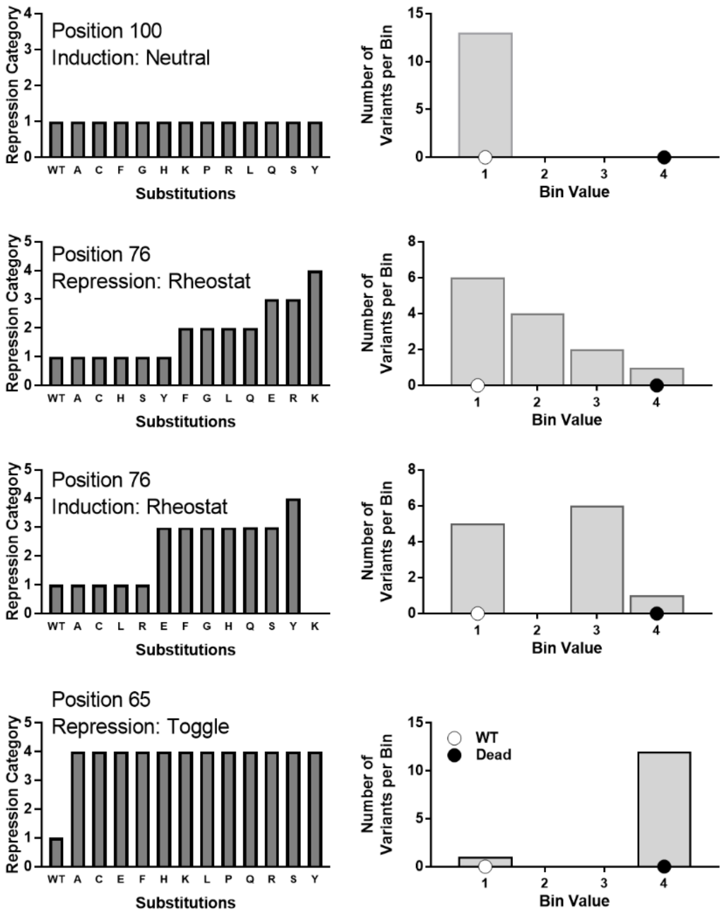


Neutral score = 1.00
Rheostat score = 0.17 (one bin occupied)
Toggle score = 0.00

Neutral score = 0.42
Rheostat score = 1.00
Toggle score = 0.08

Neutral score = 0.36
Rheostat score = 0.67
Toggle score = 0.09

Neutral score = 0.00
Rheostat score = 0.33
Toggle score = 1.00

**Supplementary Lists 1.** LacI positions comprising each substitution class (summarized in Tables 2 and 3 of main text)

Repression rheostats: 3, 7, 8, 14, 15, 24, 25, 29, 33, 34, 35, 41, 42, 45, 48, 51, 55, 58, 59, 68, 76, 81, 87, 98, 118, 122, 128, 148, 166, 182, 191, 226, 257, 259, 265, 284, 292, 301, 319, 321, 323, 4, 52, 60, 72, 78, 79, 84, 99, 117, 127, 221, 6, 17, 31, 37, 50, 67, 71, 77, 83, 94, 114, 119, 121, 124, 133, 136, 147, 150, 159, 161, 169, 171, 173, 174, 183, 185, 187, 188, 194, 218, 220, 222, 229, 233, 239, 244, 246, 249, 251, 254, 255, 256, 263, 264, 267, 269, 271, 273, 275, 278, 281, 285, 289, 296, 300, 304, 328

Repression toggles: 22, 38, 53, 56, 65, 268, 19, 247, 21, 47, 54, 252, 272, 326, 9, 20, 23, 10, 16, 179, 225, 286, 297, 5, 30, 13, 18, 49, 57, 243

Induction rheostats: 84, 192, 200, 319, 69, 107, 114, 136, 139, 188, 289, 292, 295, 4, 52, 61, 66, 68, 72, 73, 74, 76, 78, 79, 80, 92, 96, 97, 98, 99, 110, 125, 127, 142, 149, 150, 159, 160, 161, 194, 195, 196, 246, 273, 278, 296, 301, 304, 329,

Induction toggles: 75, 88, 193, 276, 191, 248, 275, 197, 95, 220, 187, 293

Double neutral: 32, 62, 91, 100, 101, 103, 105, 106, 108, 109, 111, 112, 115, 126, 130, 131, 132, 134, 135, 138, 144, 153, 155, 158, 162, 164, 165, 167, 175, 176, 180, 181, 186, 189, 190, 199, 204, 206, 207, 208, 209, 211, 212, 214, 215, 216, 217, 234, 236, 237, 238, 240, 258, 260, 261, 266, 277, 294, 302, 306, 307, 308, 309, 310, 311, 312, 313, 314, 315, 316, 317, 318, 320, 322, 325, 327

Either rheostat (both single and multiplex): 3, 7, 8, 14, 15, 24, 25, 29, 33, 34, 35, 41, 42, 45, 48, 51, 55, 58, 59, 68, 76, 81, 84, 87, 98, 118, 122, 128, 148, 166, 182, 191, 192, 200, 226, 257, 259, 265, 284, 292, 301, 319, 321, 323, 4, 52, 60, 69, 72, 78, 79, 99, 107, 114, 117, 127, 136, 139, 188, 221, 289, 295, 6, 17, 31, 37, 50, 61, 66, 67, 71, 73, 74, 77, 80, 83, 92, 94, 96, 97, 110, 119, 121, 124, 125, 133, 142, 147, 149, 150, 159, 160, 161, 169, 171, 173, 174, 183, 185, 187, 194, 195, 196, 218, 220, 222, 229, 233, 239, 244, 246, 249, 251, 254, 255, 256, 263, 264, 267, 269, 271, 273, 275, 278, 281, 285, 296, 300, 304, 328, 329

Either Toggle (both single and rheostat/toggle): 22, 38, 53, 56, 65, 75, 88, 193, 268, 19, 247, 276, 21, 47, 54, 191, 248, 252, 272, 326, 9, 20, 23, 275, 10, 16, 179, 197, 225, 286, 297, 5, 30, 95, 220, 13, 18, 49, 57, 187, 243, 293

Rheostat positions for only a single functional parameter (either repression or induction): 3, 6, 7, 8, 14, 15, 17, 24, 25, 29, 31, 33, 34, 35, 37, 41, 42, 45, 48, 50, 51, 55, 58, 59, 60, 61, 66, 67, 69, 71, 73, 74, 77, 80, 81, 83, 87, 92, 94, 96, 97, 107, 110, 117, 118, 119, 121, 122, 124, 125, 128, 133, 139, 142, 147, 148, 149, 160, 166, 169, 171, 173, 174, 182, 183, 185, 192, 195, 196, 200, 218, 221, 222, 226, 229, 233, 239, 244, 249, 251, 254, 255, 256, 257, 259, 263, 264, 265, 267, 269, 271, 281, 284, 285, 295, 300, 321, 323, 328, 329

Multiplex rheostat: Double rheostat (for both repression and induction): 4, 52, 68, 72, 76, 78, 79, 84, 98, 99, 114, 127, 136, 150, 159, 161, 188, 194, 246, 273, 278, 289, 292, 296, 301, 304, 319

Multiplex rheostat: Repression Rheostat/Induction Toggle: 187, 191, 220, 275

Unclassified: 2, 11, 12, 26, 27, 28, 36, 39, 40, 43, 44, 46, 63, 64, 70, 82, 85, 86, 89, 90, 93, 102, 104, 113, 116, 120, 123, 129, 137, 140, 141, 143, 145, 146, 151, 152, 154, 156, 157, 163, 168, 170, 172, 177, 178, 184, 198, 201, 202, 203, 205, 210, 213, 219, 223, 224, 227, 228, 230, 231, 232, 235, 241, 242, 245, 250, 253, 262, 270, 274, 279, 280, 282, 283, 287, 288, 290, 291, 298, 299, 303, 305, 324

**Supplementary Figure 3.** The locations of positions that showed rheostat (magenta ribbons) and toggle (green ball-and-stick) substitution outcomes in the repression phenotype data are mapped onto the LacI structure 1efa. As previously noted by Miller and colleagues (4, 5), the DNA binding domain contains many toggle positions. Based on prior biochemical studies of the C-subdomain interface (*e.g.* (6, 7)), we surmise that the toggle positions in the C-subdomain likely alter dimerization and/or protein stability.

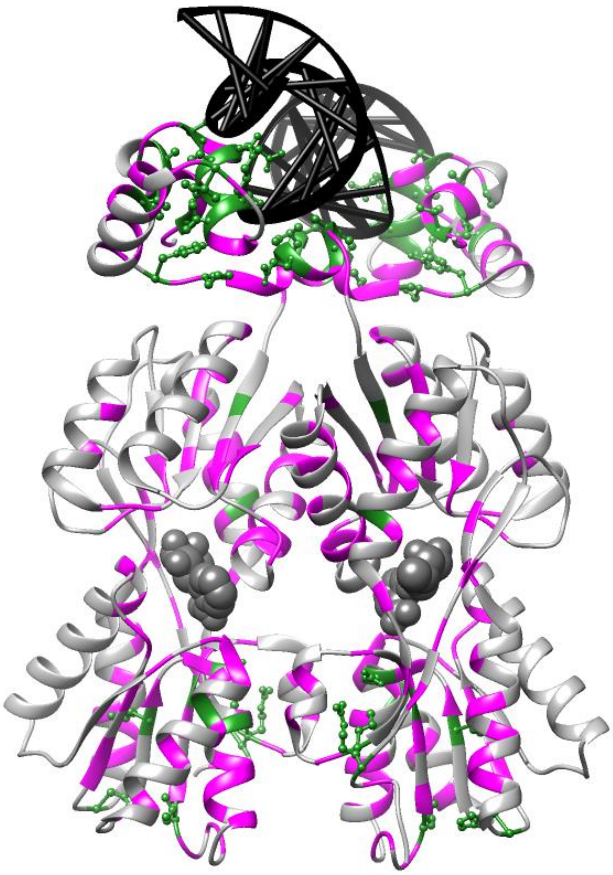

**Supplementary Figure 4**. The locations of positions that showed rheostat (magenta ribbon) and toggle (green ball-and-stick) substitution outcomes in the induction phenotype data are mapped onto the LacI structure 1efa.

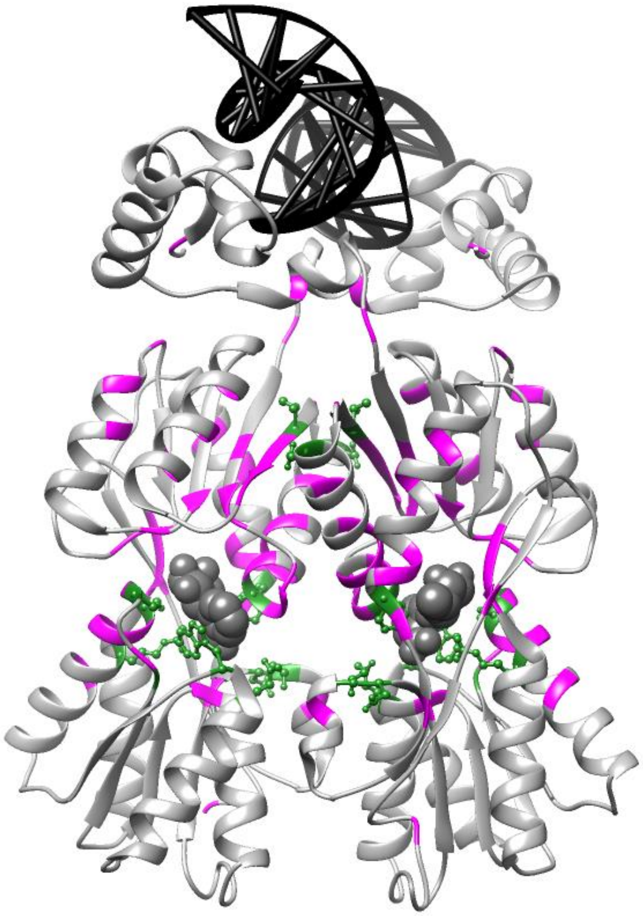

**Supplementary Figure 5.** The locations of positions that showed neutral substitution outcomes (blue ball-and-stick) for both repression and induction phenotypes are mapped onto the LacI structure 1efa. Many of these double neutral positions were solvent exposed, but not all solvent exposed positions were neutral.

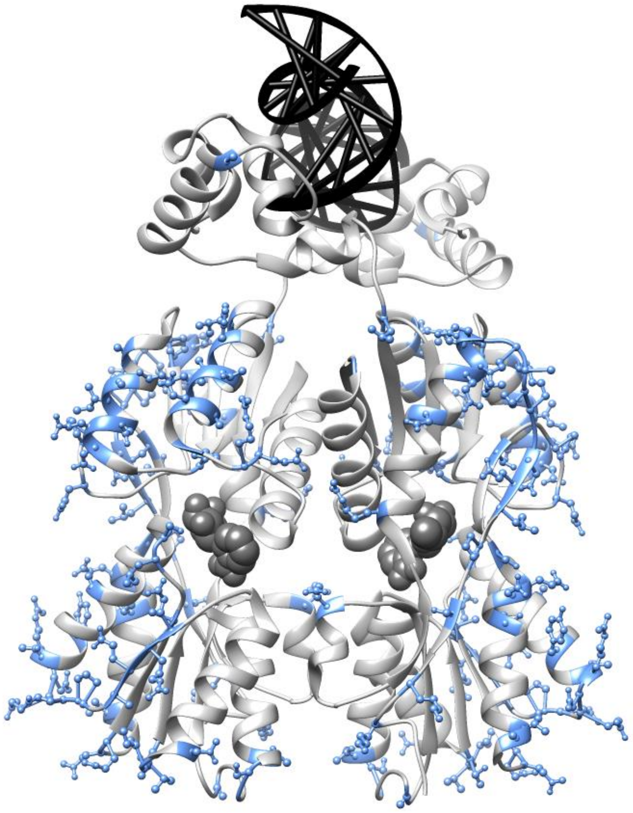

**Supplementary Figure 6.** The locations of the unclassified positions (yellow ribbon) are mapped onto the structure of LacI 1efa. These positions are distributed across the protein. These positions all showed some sensitivity to substitution, but higher resolution experimental data and/or additional substitutions are needed to classify their substitution behaviors.

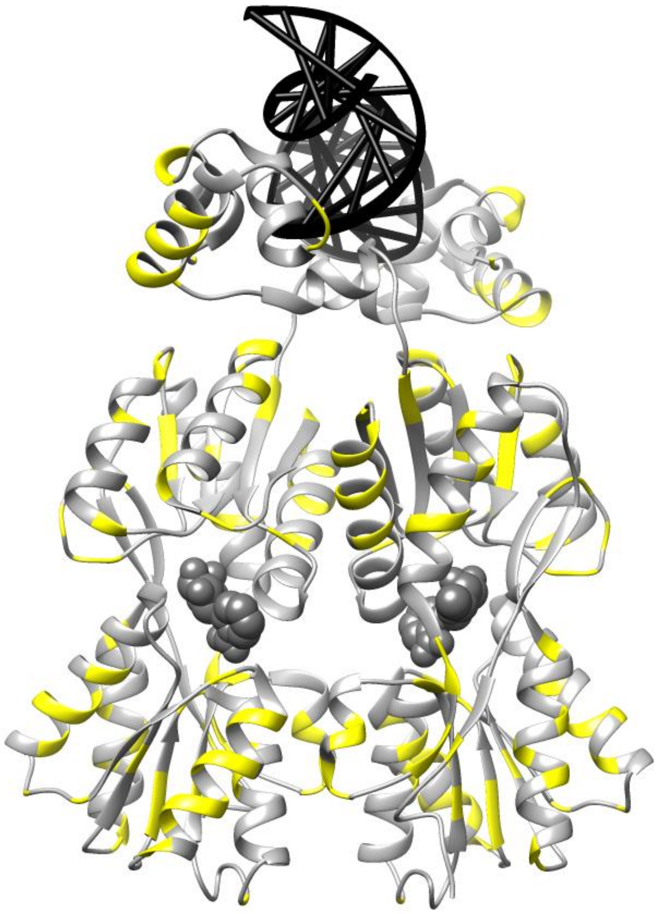

**Supplementary Lists 2.** LacI positions comprising each structural region used to construct Figure 3 of main text. Note that some positions fall into more than one of these structural regions.

Linker interfaces: 4,6,8*,9*,12,24,63,91,92,93,94,95,112,113,115,116,117,118,119,120,139,141

Linker: 45-62

DNA contacts: 3, 4, 5, 6, 7, 14, 15, 16, 17, 18, 19, 21, 22, 24, 25, 28, 29, 30, 31, 32, 34, 47, 48, 49, 50, 53, 54, 56, 57, 58, 59, 60, 118

Inducer contacts, 125, 148, 149, 160, 161, 191, 193, 197, 220, 246, 248, 274, 291

All motions: 4, 6, 8, 9, 12, 24, 45, 46, 47, 48, 49, 50, 51, 52, 53, 54, 55, 56, 57, 58, 59, 60, 61, 62, 63, 64, 66, 67, 69, 70, 71, 72, 73, 74, 77, 78, 80, 81, 83, 84, 85, 88, 90, 91, 92, 93, 94, 95, 96, 97, 98, 99, 100, 101, 102, 103, 104, 105, 106, 107, 108, 109, 110, 111, 112, 113, 114, 115, 116, 117, 118, 119, 120, 139, 141, 161, 162, 163, 164, 290, 291, 292, 293, 295, 296, 318, 319, 320

All cross-monomer interfaces: 2, 3, 4, 10, 15, 16, 17, 20, 24, 25, 27, 35, 39, 40, 43, 44, 46, 47, 48, 49, 50, 51, 52, 53, 55, 56, 63, 67, 69, 70, 71, 72, 73, 74, 77, 78, 80, 81, 84, 85, 88, 93, 94, 95, 96, 97, 98, 99, 100, 101, 102, 103, 104, 105, 106, 107, 108, 109, 110, 111, 112, 113, 114, 115, 116, 117, 118, 124, 126, 127, 128, 132, 133, 135, 136, 138, 139, 141, 142, 188, 190, 191, 219, 220, 221, 222, 223, 226, 230, 233, 234, 238, 248, 251, 252, 254, 255, 258, 259, 260, 261, 262, 263, 267, 277, 278, 280, 281, 282, 283, 284, 285, 294

*Positions 8 and 9 are present in the linker interface and thus also appear in the "all motions" list (because the linker interface changes upon DNA binding). However, positions 8 and 9 form linker interface contacts with their own chain (intra-monomer contacts). Thus, these positions are not listed in the in "all cross-monomer interfaces" (inter-monomer) list.

**Supplementary Table 2.** Statistical comparisons for the fraction positions with each substitutions class in each protein region, as compared to the fraction in the whole protein (yellow highlight). “n” is the number of positions in each region.

| | Total n | Fraction of type | Z value | p-value |
|---|---|---|---|---|
| **Either Rheostat** | | | | |
| Whole protein | 329 | 0.398 | | |
| DNA contact | 33 | 0.545 | 1.73051494 | 0.084 |
| Inducer contact | 13 | 0.615 | 1.60125574 | 0.109 |
| All motions | 88 | 0.511 | 2.172577726 | 0.030 |
| Linker | 18 | 0.556 | 1.365621018 | 0.172 |
| Linker partners | 22 | 0.455 | 0.541837749 | 0.588 |
| All interfaces | 113 | 0.504 | 2.311224838 | 0.021 |
| **Either Toggle** | | | | |
| Whole protein | 329 | 0.128 | | |
| DNA contact | 33 | 0.394 | 4.573781139 | <0.001 |
| Inducer contact | 13 | 0.385 | 2.773585589 | 0.006 |
| All motions | 88 | 0.114 | -0.393102807 | 0.694 |
| Linker | 18 | 0.333 | 2.603315993 | 0.009 |
| Linker partners | 22 | 0.091 | -0.51945728 | 0.603 |
| All interfaces | 113 | 0.115 | -0.413636969 | 0.679 |
| **Double Neutral** | | | | |
| Whole protein | 329 | 0.231 | | |
| DNA contact | 33 | 0.030 | 0.949643018 | 0.3423 |
| Inducer Contact | 13 | 0.000 | -0.397359707 | 0.6911 |
| All motions | 88 | 0.182 | 14.64607191 | <0.001 |
| Linker | 18 | 0.056 | 1.7144303 | 0.0864 |
| Linker partners | 22 | 0.136 | 5.341508579 | <0.001 |
| All interfaces | 113 | 0.195 | 17.8657485 | <0.001 |
| **Double Rheostat** | | | | |
| Whole protein | 329 | 0.082 | | |
| DNA contact | 33 | 0.030 | -1.088759724 | 0.276 |
| Inducer contact | 13 | 0.154 | 0.946184667 | 0.344 |
| All motions | 88 | 0.136 | 1.846319383 | 0.065 |
| Linker | 18 | 0.056 | -0.402051173 | 0.688 |
| Linker partners | 22 | 0.045 | -0.632535344 | 0.527 |
| All interfaces | 113 | 0.106 | 0.929869915 | 0.352 |
| **Single Rheostat** | | | | |
| Whole protein | 329 | 0.310 | | |
| DNA contact | 33 | 0.330 | 0.248417149 | 0.804 |
| Inducer contact | 13 | 0.460 | 1.169385065 | 0.242 |
| All motions | 88 | 0.375 | 1.318406393 | 0.187 |
| Linker | 18 | 0.500 | 1.742948124 | 0.081 |
| Linker partners | 22 | 0.409 | 1.004017177 | 0.315 |
| All interfaces | 113 | 0.398 | 2.022630266 | 0.043 |
| **Rheostat/Toggle** | | | | |
| Whole protein | 329 | 0.012 | | |
| DNA contact | 33 | 0.000 | -0.633095346 | 0.527 |
| Inducer contact | 13 | 0.154 | 4.702089868 | <0.001 |
| All motions | 88 | 0.000 | -1.03384037 | 0.301 |
| Linker | 18 | 0.000 | -0.4675719 | 0.640 |
| Linker partners | 22 | 0.000 | -0.516920185 | 0.605 |
| All interfaces | 113 | 0.018 | 0.585762246 | 0.558 |

**Supplementary Figure 7.** FoldX calculations (8) for LacI variants in the Miller dataset. (A-B) Stability changes were calculated for all 19 substitutions at all positions in the LacI regulatory domains and compared to repression phenotypes. The structure used was 1lbi, which comprises four chains of the homotetrameric LacI regulatory and tetramerization domains in the absence of bound ligand (9).

A total of 3299 variants were analyzed: 2453 with the wild-type "+" phenotype, 191 with the intermediate "+/-" phenotype, 84 with even weaker "-/+" and 571 with the poorest "-" repression phenotype). For clarity on the plots, a small number* of substitutions with $\Delta\Delta G > 15$ are not shown. In general, the distributions correlated with repression phenotype: strong repressors exhibited a lower distribution of FoldX scores (more stable) and weak repressors showed higher scores (less stable). This trend is statistically significant: Kruskal-Wallis $X^2 = 781.39$; $p < 0.001$ for "Raw Predictions". For reference, $\Delta\Delta G = 3$ (dotted line) is shown as an energy that is likely to be de-stabilizing. (The effects of varied significance thresholds are explored below in Supplementary Figure 8.) However, >20% of variants that repress strongly ("+" phenotype) are incorrectly predicted to be severely destabilized, with $\Delta\Delta G > 3$. This limits the usefulness of FoldX as a method for separating variants that affect stability from variants that affect other aspects of function.

(A) "Raw Predictions" comprise the average score assigned across the four chains for each position. (B) "Outlier removed/tight confidence" scores exclude statistical outliers, as follows: In the raw scores, sometimes either (i) the score for one chain disagreed with the other three chains or (ii) the 95% confidence interval for predicted stability was wide. We considered the possibility that these effects might be calculation anomalies and thereby introduce noise into the analysis. However, removal of these data points (see Supplementary Methods on the next page) did not affect the interpretation of the results.

*For "Raw Predictions" data, 60 data "+" points with $\Delta\Delta G > 15$ were excluded from the plots, as were 10 "+/-", 4 "-/+" and 107 "-". For "Outlier-removed" data, 43 "+" data points were excluded from the plots, as were 8 "+/-", 4 "-/+" and 108 "-".

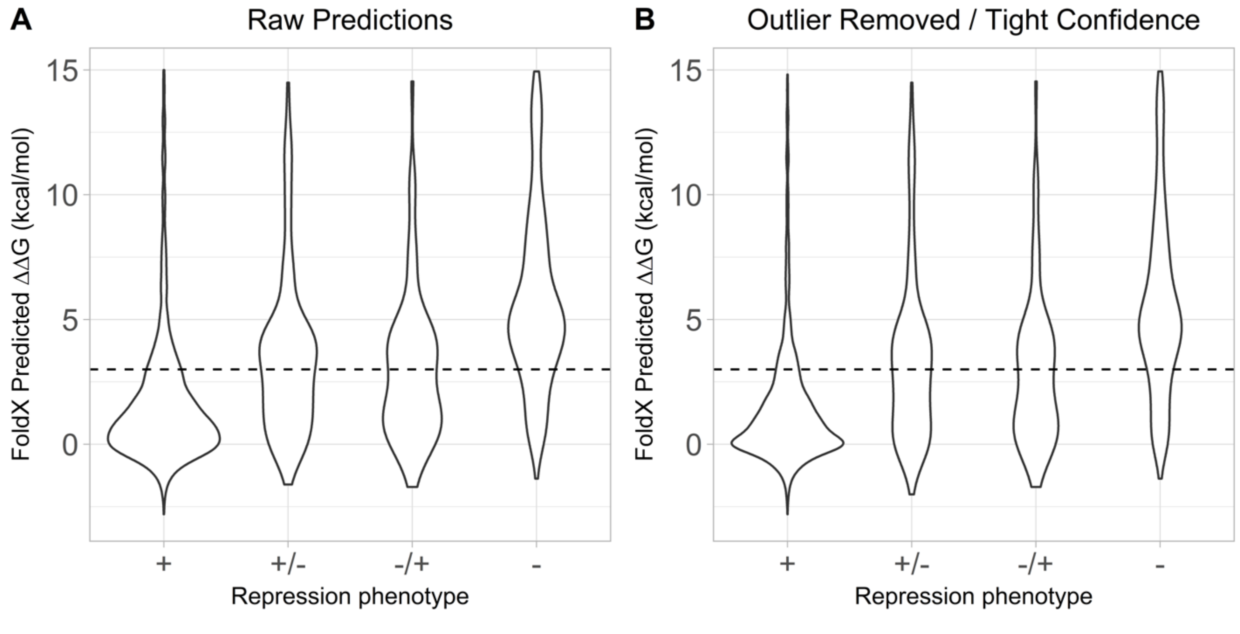

**Supplementary Figure 8**. Although Supplementary Figure 7 uses a value of ΔΔG > 3 as an example de-stabilizing energy change, the true threshold energy at which LacI de-stabilization results in changed phenotype is unknown. Indeed, the published literature on FoldX appears to lack consensus about what change in free energy is required for protein destabilization. Arguments have been raised that there is no one-size-fits-all-proteins destabilization threshold (10). Various studies have considered ΔΔG thresholds between 1.84 kcal/mol and 5 kcal/mol (11-13). More generally, analysis of experimentally-determined ΔΔG values in the ProTherm database demonstrates that most amino acid changes have ΔΔG values in the range -3 to 5 kcal/mol, and that there is also a sharp drop-off in the histogram past even 2.5 kcal/mol (14) The ΔΔG thresholds chosen for the "outlier" analysis in Supplementary Figure 7B and Supplementary Figure 8B – 2 kcal/mol and 3kcal/mol – were chosen to fall within the general range of likely-destabilizing energies.

Nevertheless, LacI appears to be an unusually stable protein (15). Thus, for the data plotted in Supplementary Figure 7A and B, we also considered the fraction of repressors that were predicted to be de-stabilized as a function of increasingly stringent free energy thresholds. At all thresholds, and as expected for successful predictions, repressors with greater *in vivo* repression were associated with greater predicted stability *in silico*.

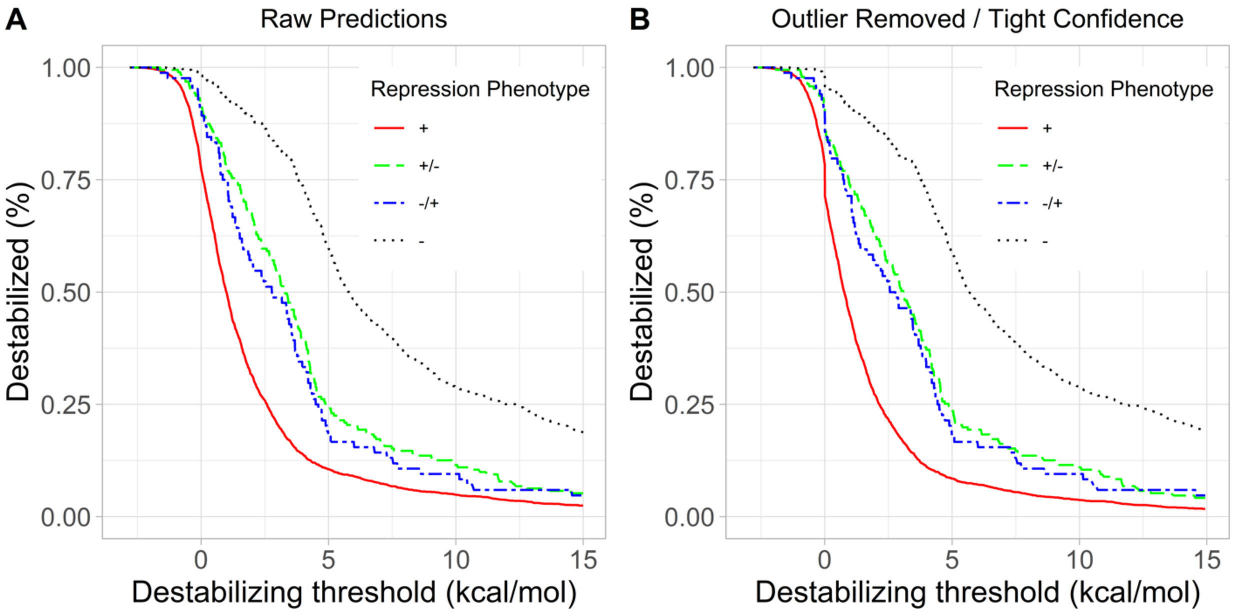


**Supplementary Methods: FoldX outlier removal.** FoldX calculations for one chain in the LacI tetramer structure sometimes disagreed with calculations for the other three chains. Notably, the 4 chains are not perfectly symmetric in each structure. Thus, one interpretation of the outliers is that the chain was accessing a conformation for which the substitution was truly destabilizing. An alternative interpretation is that the outlier chain was due to a calculation artefact; as such, these outliers could explain why some "wild-type" like repressors were falsely predicted to have greatly diminished stability.

Thus, to identify outliers, the inter-quartile range (IQR) of the four ΔΔG was first determined. Any ΔΔG below the first quartile - 1.5 * IQR or exceeding the third quartile + 1.5 * IQR was classified as an outlier and removed from the four-chain average. After outlier removal, we again assessed whether variants agreed among the remaining chains, based on their 95% confidence intervals. For this process, we first hypothesized that substitution variants may start becoming destabilized at ΔΔG > 2 kcal/mol and are more likely to be destabilized at ΔΔG > 3 kcal/mol. For extremely destabilizing mutants (ΔΔG > 3 kcal/mol), it does not matter precisely how destabilizing they are (*e.g.* they unfold all the same whether ΔΔG is 5 kcal/mol or 50 kcal/mol). Thus, for extremely destabilizing mutants (ΔΔG > 3 kcal/mol), we excluded variants only if the 95% confidence interval included ΔΔG < 2 kcal/mol (*i.e.* contained some values that were not clearly destabilizing). For stabilizing (ΔΔG < 0) or not extremely destabilizing (ΔΔG < 3 kcal/mol), mutants were excluded from further analyses if the half-width of their 95% confidence interval exceeded 1.5 kcal/mol, which is 75% of the change required to change classification from stabilizing (ΔΔG < 0 kcal/mol) to destabilizing (ΔΔG > 2 kcal/mol).

For 1lbi, 213 low-energy (ΔΔG < 3) and 152 high-energy variants were excluded due to wide confidence intervals, and 5555 mutants remained to be further analyzed.

**Supplementary Figure 9.** RheoScale analyses of stability changes predicted by FoldX, stratified by phenotype substitution classes (Table 2).

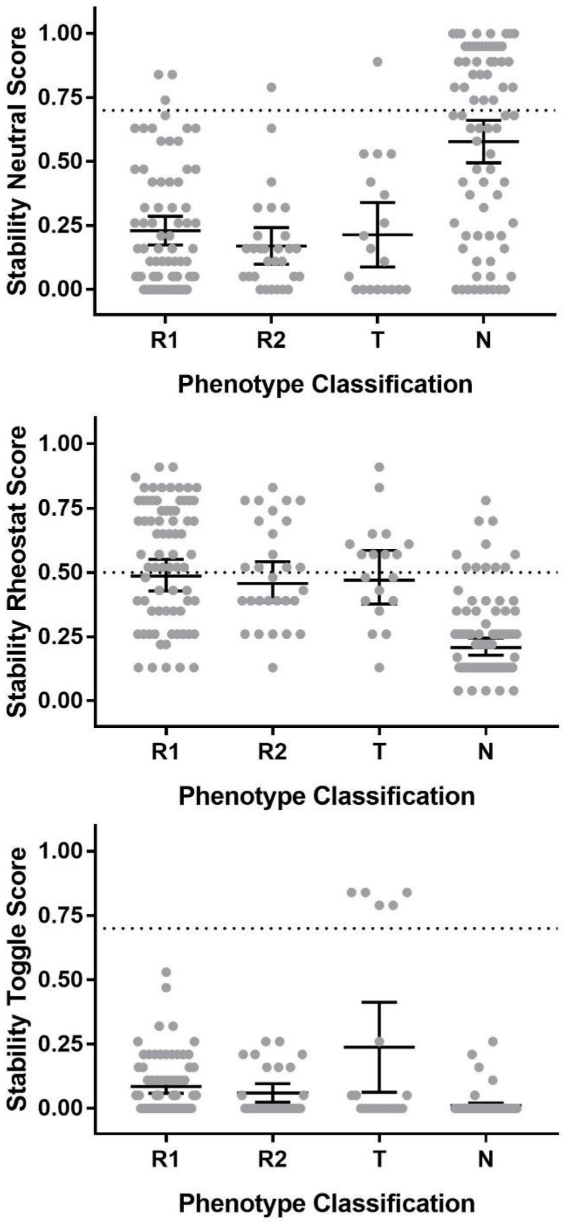


To assess whether each position was predicted to be a rheostat, toggle, or neutral position for stability, the stability scores generated by FoldX were analyzed using the RheoScale calculator. As such, each position's RheoScale scores (Y axes) were derived from all 20 possible substitutions.

Methods: Analysis parameters included a bin number of 10 and a "max override" value of 15. The latter prevents the small percent of positions that were predicted to be extremely destabilizing from dominating analyses. Since FoldX predicts ΔΔG values, stabilities were analyzed without an additional log conversion. The dotted horizontal lines on each panel correspond to empirically observed significance thresholds for RheoScale scores, as determined for other datasets (16, 17).

Results: RheoScale stability scores were then compared to the phenotype substitution outcomes identified for LacI positions (Table 2). "R1" corresponds to the group of single rheostat positions. "R2" are the double rheostat positions; "T" are toggle positions and "N" are double neutral positions.

Two control conditions are available to assess the predicted stability changes: First, phenotypically neutral substitutions for repression should have only small predicted changes in stability (Supplementary Figure 7, above) and phenotypically double neutral positions should have low rheostat and toggle stability scores (this figure, middle and lower panels). Second, positions with high toggle scores for stability should be phenotypically assigned to the toggle category (lower panel). Note that toggle phenotypes can arise from functional effects that are distinct from stability. Only the second criterion was met; several neutral substitutions and neutral positions were incorrectly predicted to have large effects on stability. This leaves us reluctant to draw strong conclusions from these calculations. Nevertheless, it is interesting to note that substitutions at many of the single (R1) and double (R2) rheostat positions are not predicted to have large effects on LacI stability (as indicated by low rheostat and toggle scores).

**Supplementary Table 3.** Phenotypes for each amino acid substitution at double rheostat positions (Figure 7 of main text and Supplementary Figure 10). A value of "1" corresponds to the wild-type phenotype. For many of the variants with poorest repression ("4"), induction could not be assessed (blank values).

| Substitution | Repression | Induction |
|---|---|---|
| 4A | 1 | 4 |
| 4C | 1 | 3 |
| 4E | 3 | 1 |
| 4F | 1 | 1 |
| 4H | 1 | 1 |
| 4L | 2 | 1 |
| 4P | 1 | 4 |
| 4R | 1 | 1 |
| 4S | 1 | 4 |
| 4G | 2 | 1 |
| 4K | 1 | 1 |
| 4Q | 2 | 1 |
| 4Y | 1 | 3 |
| 52A | 1 | 4 |
| 52C | 1 | 4 |
| 52E | 3 | 1 |
| 52F | 1 | 1 |
| 52G | 2 | 1 |
| 52H | 1 | 4 |
| 52K | 3 | 1 |
| 52L | 1 | 3 |
| 52P | 1 | 1 |
| 52Q | 1 | 1 |
| 52R | 3 | 1 |
| 52S | 1 | 3 |
| 52Y | 1 | 1 |
| 68A | 1 | 1 |
| 68C | 1 | 1 |
| 68E | 1 | 4 |
| 68F | 2 | 1 |
| 68G | 1 | 1 |
| 68H | 2 | 1 |
| 68K | 4 | |
| 68L | 2 | 3 |
| 68P | 2 | 1 |
| 68Q | 3 | 1 |
| 68R | 4 | |
| 68S | 1 | 1 |
| 68Y | 2 | 3 |
| 72C | 1 | 1 |
| 72E | 1 | 1 |
| 72F | 1 | 4 |
| 72G | 1 | 1 |
| 72H | 1 | 1 |
| 72K | 1 | 1 |
| 72L | 2 | 1 |
| 72P | 3 | 1 |
| 72Q | 1 | 1 |
| 72R | 1 | 1 |
| 72S | 1 | 1 |
| 72Y | 1 | 3 |
| 76A | 1 | 1 |
| 76C | 1 | 1 |
| 76E | 3 | 3 |
| 76F | 2 | 3 |
| 76G | 2 | 3 |
| 76H | 1 | 3 |
| 76K | 4 | |
| 76L | 2 | 1 |
| 76Q | 2 | 3 |
| 76R | 3 | 1 |
| 76S | 1 | 3 |
| 76Y | 1 | 4 |
| 78A | 2 | 3 |
| 78C | 1 | 4 |
| 78E | 3 | 1 |
| 78F | 1 | 1 |
| 78G | 1 | 3 |
| 78H | 1 | 3 |
| 78P | 2 | 4 |
| 78R | 1 | 4 |
| 78K | 1 | 4 |
| 78L | 2 | 1 |
| 78S | 1 | 1 |
| 78Y | 2 | 1 |
| 79A | 1 | 1 |
| 79C | 1 | 1 |
| 79E | 2 | 1 |
| 79F | 1 | 3 |
| 79G | 2 | 1 |

| Substitution | Repression | Induction |
|---|---|---|
| 79H | 1 | 4 |
| 79K | 2 | 3 |
| 79L | 1 | 3 |
| 79P | 2 | 3 |
| 79Q | 2 | 1 |
| 79R | 3 | 3 |
| 79S | 1 | 1 |
| 79Y | 2 | 3 |
| 84A | 1 | 4 |
| 84C | 1 | 4 |
| 84E | 1 | 3 |
| 84F | 1 | 4 |
| 84G | 1 | 4 |
| 84H | 1 | 3 |
| 84P | 3 | 2 |
| 84R | 1 | 1 |
| 84L | 2 | 4 |
| 84Q | 3 | 3 |
| 84S | 1 | 4 |
| 84Y | 1 | 4 |
| 98A | 2 | 4 |
| 98C | 1 | 1 |
| 98E | 2 | 3 |
| 98F | 1 | 3 |
| 98G | 3 | 4 |
| 98H | 2 | 4 |
| 98K | 4 | |
| 98L | 2 | 1 |
| 98P | 1 | 3 |
| 98Q | 1 | 4 |
| 98R | 1 | 3 |
| 98S | 2 | 3 |
| 98Y | 2 | 1 |
| 99A | 2 | 1 |
| 99C | 1 | 1 |
| 99E | 2 | 1 |
| 99F | 1 | 4 |
| 99G | 3 | 1 |
| 99H | 2 | 1 |
| 99K | 1 | 4 |
| 99L | 1 | 3 |
| 99P | 2 | 4 |
| 99Q | 1 | 1 |
| 99R | 1 | 4 |

| Substitution | Repression | Induction |
|---|---|---|
| 99S | 1 | 1 |
| 99Y | 1 | 1 |
| 114A | 1 | 1 |
| 114C | 1 | 1 |
| 114E | 4 | |
| 114F | 1 | 2 |
| 114G | 1 | 1 |
| 114H | 1 | 1 |
| 114K | 4 | |
| 114P | 4 | |
| 114Q | 1 | 1 |
| 114R | 4 | |
| 114S | 1 | 1 |
| 114Y | 2 | 3 |
| 127A | 1 | 4 |
| 127C | 1 | 1 |
| 127E | 1 | 4 |
| 127F | 1 | 1 |
| 127G | 1 | 4 |
| 127H | 1 | 3 |
| 127K | 3 | 3 |
| 127L | 2 | 3 |
| 127Q | 1 | 4 |
| 127R | 1 | 3 |
| 127S | 1 | 3 |
| 127Y | 1 | 1 |
| 136A | 1 | 1 |
| 136C | 1 | 1 |
| 136E | 1 | 1 |
| 136F | 2 | 1 |
| 136G | 2 | 1 |
| 136H | 1 | 3 |
| 136K | 4 | 2 |
| 136L | 1 | 1 |
| 136P | 4 | |
| 136Q | 1 | 3 |
| 136R | 4 | 2 |
| 136S | 1 | 1 |
| 136Y | 1 | 3 |
| 150A | 1 | 1 |
| 150C | 1 | 4 |
| 150E | 4 | 3 |
| 150F | 4 | |
| 150G | 1 | 1 |

| Substitution | Repression | Induction |
|---|---|---|
| 150H | 2 | 1 |
| 150K | 1 | 1 |
| 150L | 1 | 3 |
| 150P | 4 | 3 |
| 150Q | 1 | 3 |
| 150R | 4 | |
| 150S | 1 | 3 |
| 150Y | 1 | 3 |
| 159A | 1 | 1 |
| 159C | 1 | 1 |
| 159E | 1 | 1 |
| 159F | 1 | 4 |
| 159G | 1 | 3 |
| 159H | 1 | 1 |
| 159K | 4 | |
| 159L | 1 | 3 |
| 159P | 3 | 3 |
| 159Q | 1 | 3 |
| 159R | 1 | 4 |
| 159S | 1 | 3 |
| 159Y | 1 | 3 |
| 161A | 4 | |
| 161C | 4 | |
| 161E | 4 | 3 |
| 161G | 4 | 4 |
| 161H | 1 | 3 |
| 161K | 4 | |
| 161L | 1 | 3 |
| 161P | 4 | 4 |
| 161Q | 4 | 3 |
| 161R | 4 | |
| 161S | 3 | 4 |
| 161Y | 3 | 1 |
| 162A | 1 | 1 |
| 188A | 1 | 1 |
| 188C | 1 | 1 |
| 188E | 1 | 1 |
| 188F | 4 | |
| 188G | 1 | 3 |
| 188H | 1 | 1 |
| 188K | 1 | 2 |
| 188L | 1 | 1 |
| 188Q | 1 | 1 |
| 188R | 1 | 3 |

| Substitution | Repression | Induction |
|---|---|---|
| 188S | 1 | 1 |
| 188Y | 2 | 1 |
| 194C | 1 | 3 |
| 194E | 1 | 4 |
| 194F | 4 | |
| 194G | 1 | 1 |
| 194H | 2 | 1 |
| 194K | 4 | |
| 194L | 1 | 1 |
| 194P | 1 | 4 |
| 194Q | 1 | 4 |
| 194R | 4 | |
| 194S | 1 | 1 |
| 194Y | 1 | 4 |
| 246A | 1 | 3 |
| 246C | 1 | 4 |
| 246E | 1 | 4 |
| 246F | 4 | |
| 246G | 1 | 1 |
| 246H | 1 | 4 |
| 246K | 3 | 3 |
| 246L | 3 | 3 |
| 246P | 4 | |
| 246Q | 1 | 4 |
| 246R | 1 | 4 |
| 246S | 1 | 3 |
| 246Y | 3 | 3 |
| 273A | 1 | 4 |
| 273C | 1 | 4 |
| 273E | 4 | |
| 273F | 1 | 4 |
| 273G | 1 | 4 |
| 273H | 1 | 3 |
| 273K | 4 | |
| 273P | 4 | |
| 273R | 4 | |
| 273L | 1 | 1 |
| 273Q | 2 | 4 |
| 273S | 1 | 4 |
| 278A | 1 | 4 |
| 278C | 1 | 3 |
| 278E | 1 | 4 |
| 278F | 3 | 1 |
| 278G | 1 | 1 |

| Substitution | Repression | Induction |
|---|---|---|
| 278H | 1 | 1 |
| 278K | 4 | |
| 278L | 1 | 4 |
| 278P | 1 | 1 |
| 278Q | 1 | 1 |
| 278R | 4 | |
| 278S | 1 | 1 |
| 278Y | 1 | 1 |
| 289A | 4 | |
| 289C | 1 | 1 |
| 289E | 4 | |
| 289F | 3 | 3 |
| 289G | 4 | |
| 289H | 4 | 2 |
| 289K | 4 | |
| 289L | 1 | 1 |
| 289P | 4 | |
| 289Q | 4 | |
| 289R | 4 | |
| 289S | 4 | |
| 289Y | 4 | |
| 292A | 1 | 1 |
| 292C | 1 | 1 |
| 292E | 1 | 3 |
| 292F | 4 | |
| 292G | 1 | 2 |
| 292H | 2 | 1 |
| 292K | 3 | 1 |
| 292L | 4 | 2 |
| 292P | 1 | 1 |
| 292Q | 1 | 3 |
| 292R | 2 | 1 |
| 292S | 1 | 1 |
| 292Y | 4 | |
| 296A | 1 | 1 |
| 296C | 1 | 1 |
| 296E | 1 | 1 |
| 296F | 1 | 4 |
| 296G | 1 | 1 |
| 296H | 1 | 3 |
| 296P | 3 | 3 |
| 296Q | 1 | 1 |

| Substitution | Repression | Induction |
|---|---|---|
| 296R | 4 | |
| 296Y | 1 | 4 |
| 296K | 1 | 4 |
| 296S | 1 | 1 |
| 301A | 1 | 1 |
| 301C | 1 | 1 |
| 301E | 4 | |
| 301F | 1 | 2 |
| 301G | 3 | 1 |
| 301H | 1 | 1 |
| 301K | 4 | |
| 301L | 1 | 1 |
| 301P | 4 | 2 |
| 301Q | 4 | |
| 301R | 4 | |
| 301S | 2 | 4 |
| 301Y | 1 | 2 |
| 304A | 1 | 3 |
| 304C | 1 | 1 |
| 304E | 1 | 1 |
| 304F | 1 | 1 |
| 304G | 2 | 1 |
| 304H | 1 | 3 |
| 304K | 4 | |
| 304P | 4 | |
| 304R | 2 | 4 |
| 304Q | 1 | 3 |
| 304S | 1 | 3 |
| 304Y | 1 | 1 |
| 319A | 1 | 2 |
| 319C | 1 | 1 |
| 319E | 1 | 3 |
| 319F | 1 | 4 |
| 319G | 3 | 1 |
| 319H | 1 | 3 |
| 319K | 3 | 1 |
| 319P | 4 | |
| 319R | 2 | 3 |
| 319Q | 1 | 3 |
| 319S | 2 | 3 |
| 319Y | 1 | 3 |

**Supplementary Figure 10.** Correlation plots for all substitution phenotypes at each double rheostat position show no correlation. This suggests that varied substitutions at the same location may differentially affect the composite functional parameter(s) shown in Figure 2. The raw data for these plots are in Supplementary Table 3, above. Note that some data are stacked on top of each other in the plots and are thus not visible; other substitutions are missing from the plots because the induction phenotype could not be measured for substitutions with weak repression.

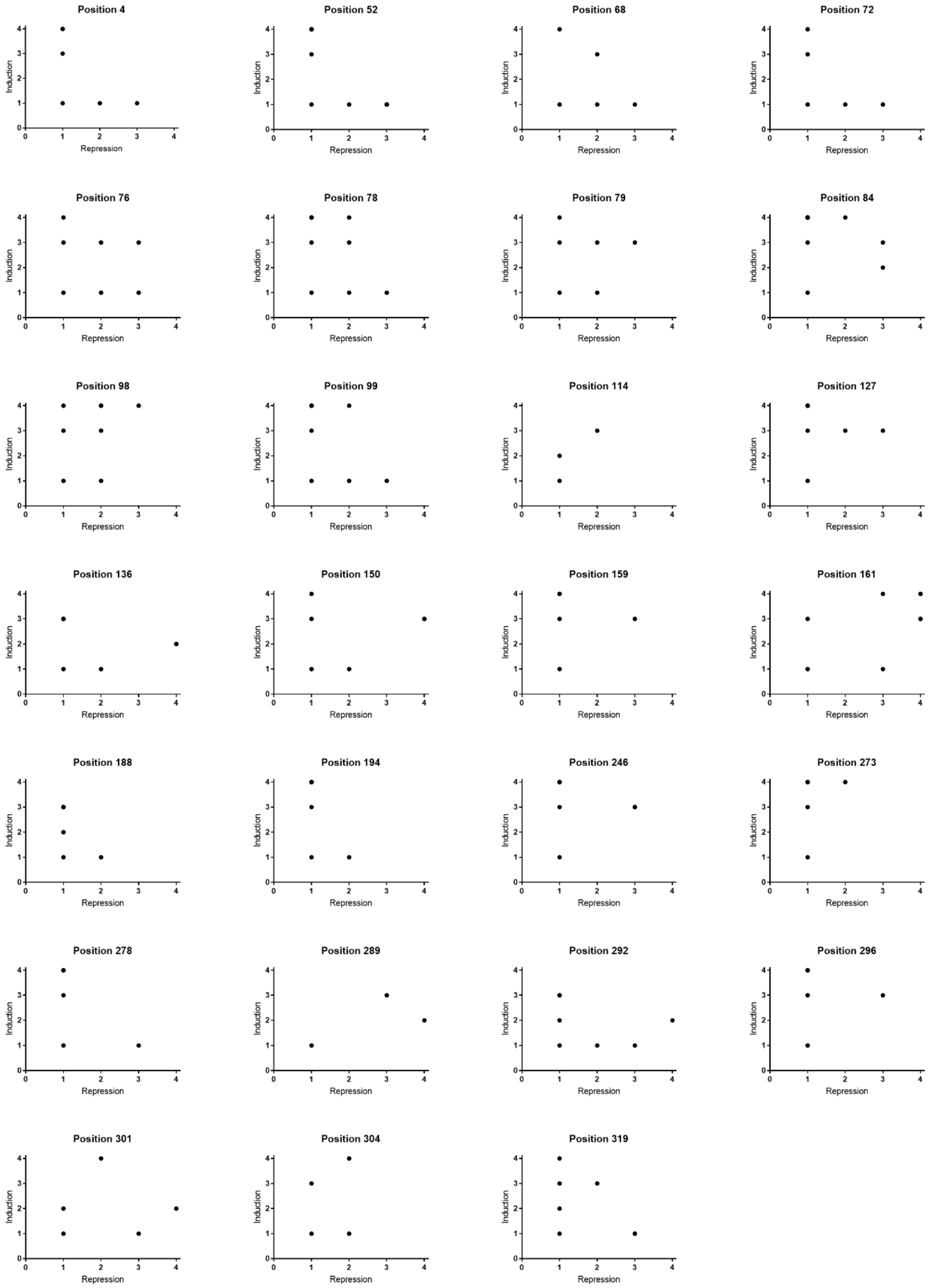

**Supplementary Figure 11**. Distributions of bioinformatic scores for single (R1), double (R2), toggle, and double neutral positions were compared for 16 analyses of the LacI/GalR family. Four plots are shown per page. The notation “351” indicates that the sequence alignment used in analyses represented the full LacI/GalR family (see Methods, main text). Averages (red dots) and their 95% confidence limits (blue lines) are shown for each distribution. Each of the Y axes corresponds to the score range for each different method. On the x axes labels, numbers in parentheses indicate the number of LacI positions shown experimentally to occur in each group. At the bottom of each page, p values for comparisons of the single (R1) and double (R2) rheostat positions are listed; statistically significant p values are highlighted with red text. The plots for ConSurf and the score created from combining seven analyses are shown in main text Figure 5.

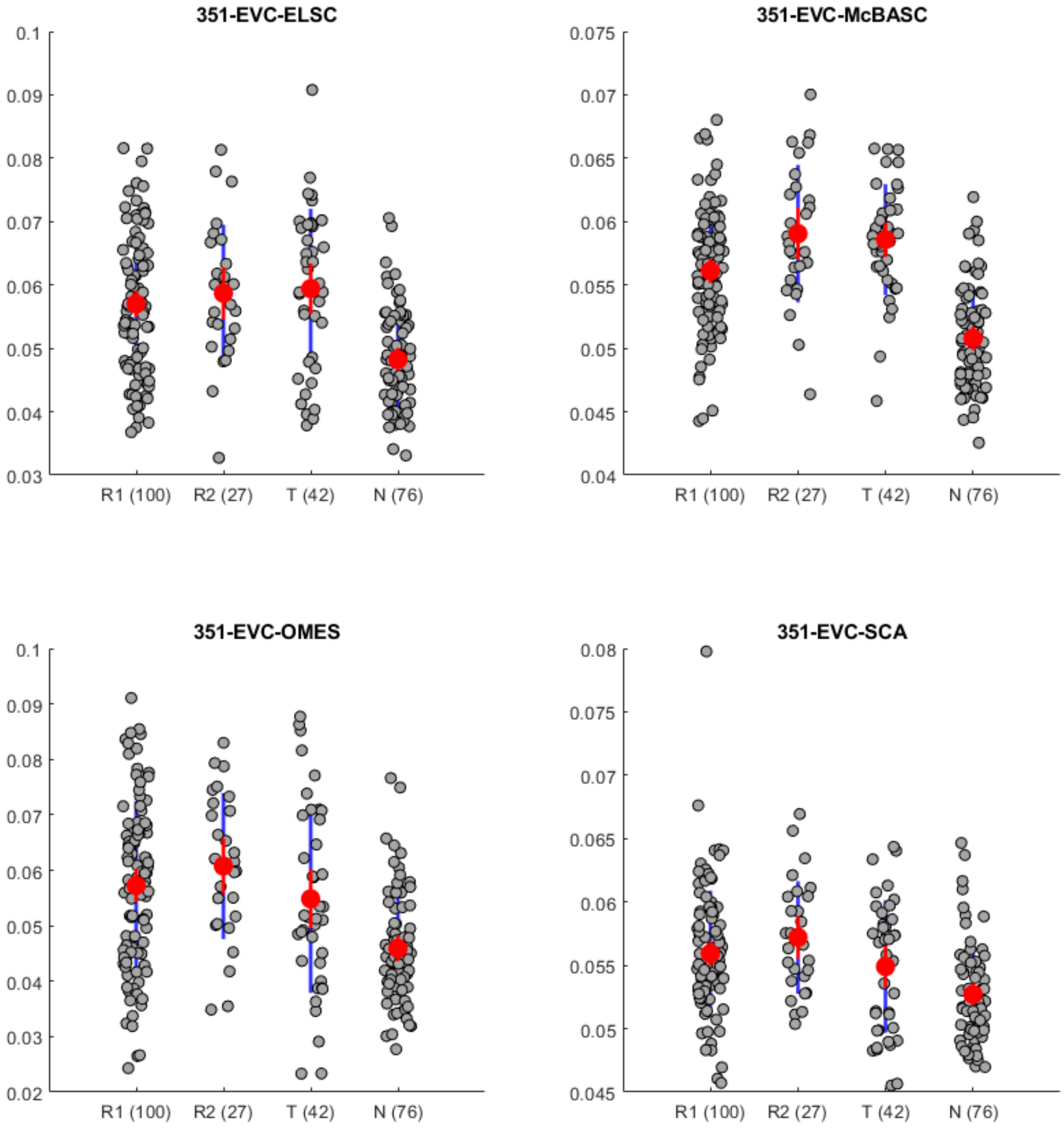


P-values for comparing R1 vs R2 for eigenvector centrality (“EVC”) analyses ELCS, McBASC, OMES, and SCA are respectively: 0.4315, 0.0137, 0.2770, and 0.2020.

**Supplementary Figure 11, cont.**

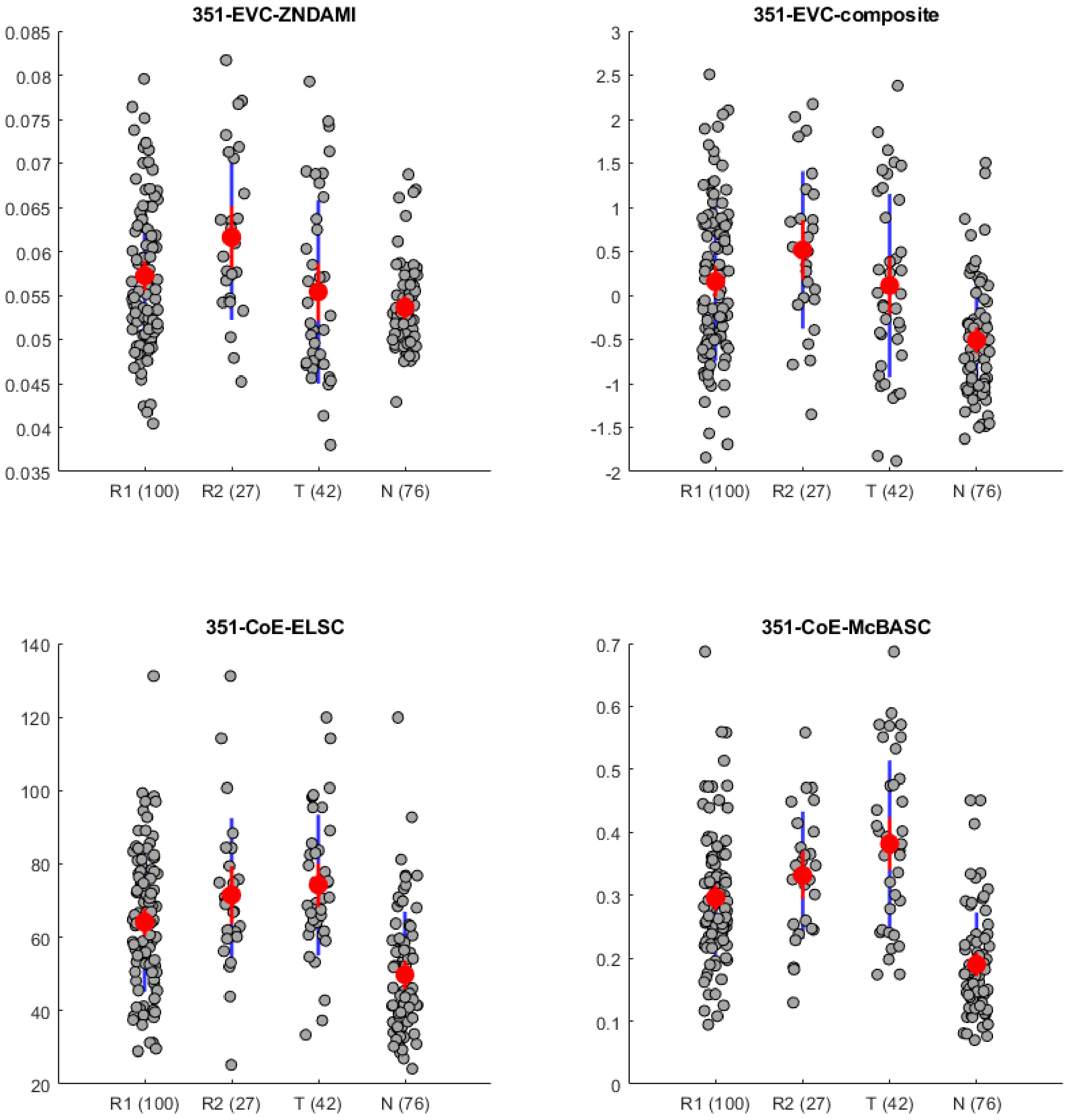


P-values for comparing R1 vs R2 for eigenvector centrality ("EVC") analyses ZNDAMI and the composite eigenvector centrality score (derived from the 5 eigenvector centrality scores), as well as the pairwise co-evolutionary ("Co-E") methods ELSC and McBASC are respectively: 0.0316, 0.0709, 0.1292, and 0.0677.

**Supplementary Figure 11, cont.**

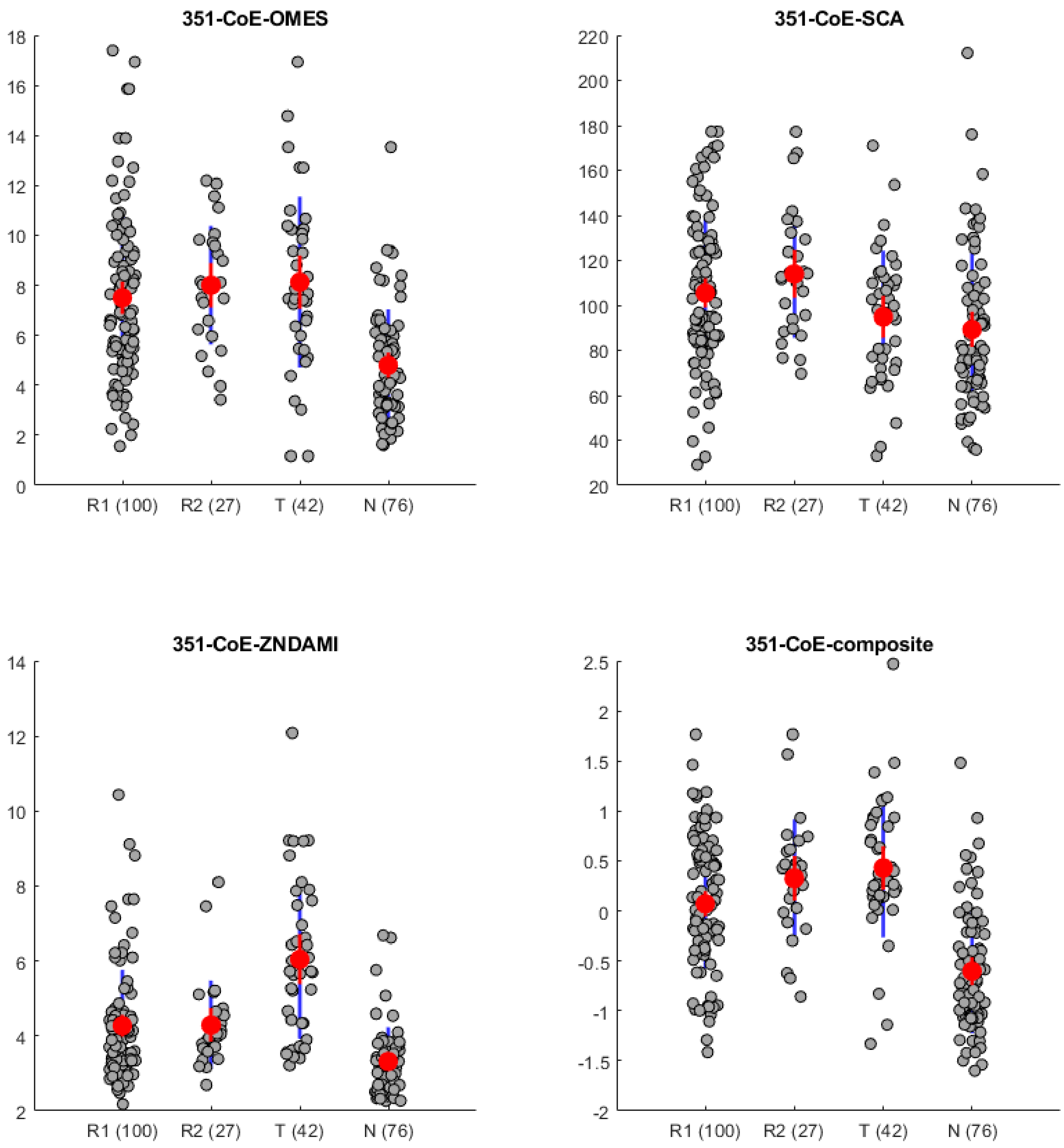


P-values for comparing R1 vs R2 for the pairwise co-evolutionary ("Co-E") methods OMES, SCA, and ZNDAMI, and the composite Co-E score (derived from the 5 Co-E scores) are respectively: 0.2351, 0.2094, 0.4443, and 0.0806.

**Supplementary Figure 11, cont.**

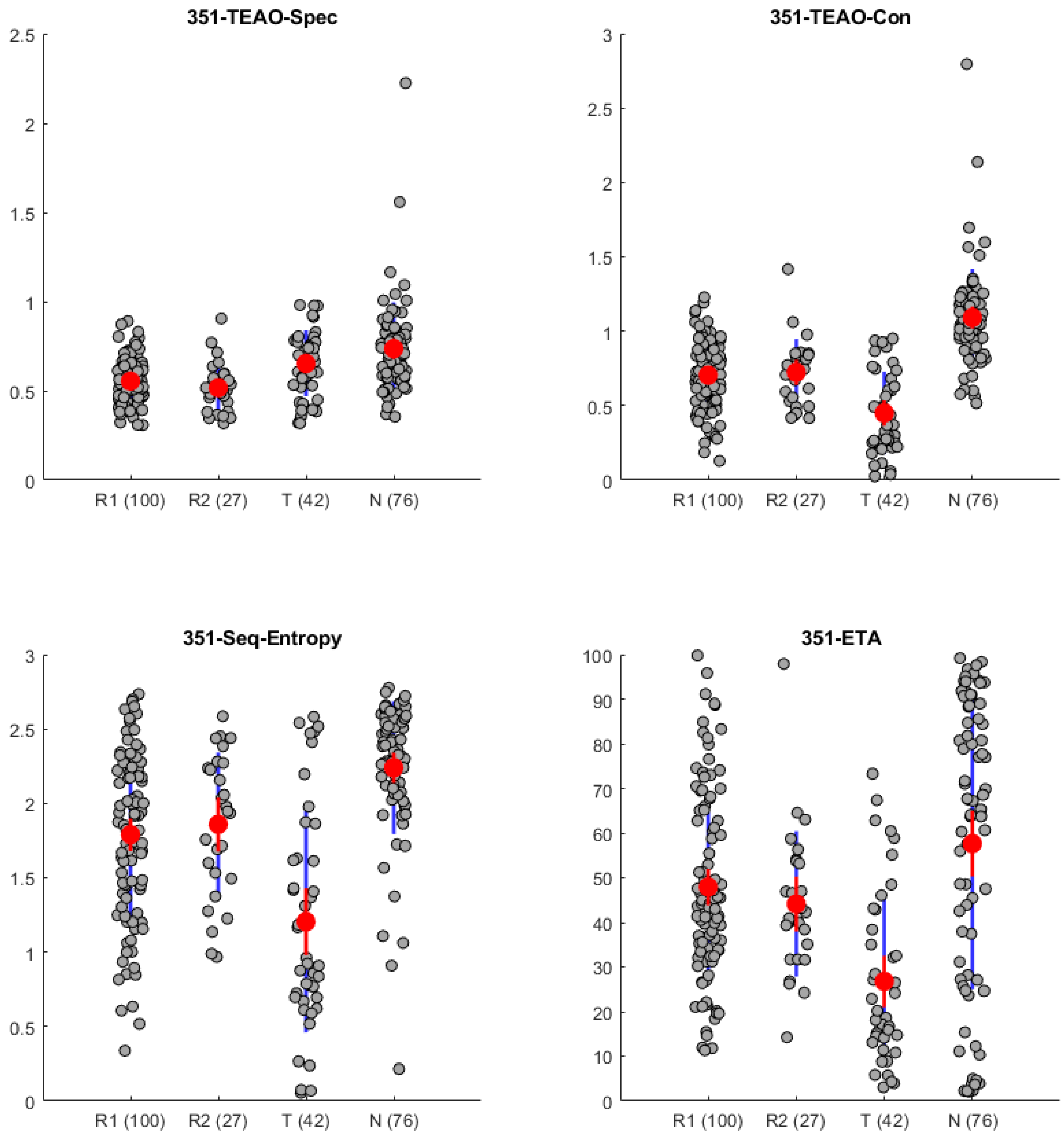


P-values for comparing R1 vs R2 for TEAO-Specificity, TEAO-Conserved, Sequence Entropy, and Evolutionary Trace Analysis scores are respectively: 0.1123, 0.9507, 0.6061, and 0.4332.

**Supplementary Table 4.** VUS for bioinformatic analyses. The cutoff thresholds used to separate the three outcomes are list in Supplementary Table 5.

| Analysis | Outcome Order* | Empirical VUS** |
|---|---|---|
| EVC-ELSC | N<R<T, | 0.3531(0.2686,0.4458) |
| EVC-McBASC | N<R<T, | 0.4585(0.3704,0.5421) |
| EVC-OMES | N<T<R, | 0.2832(0.2127,0.3561) |
| EVC-SCA | N<R<T, | 0.2736(0.1880,0.3641) |
| EVC-ZNDAMI | T<N<R | 0.3125(0.2227,0.4122) |
| EVC-composite*** | N<T<R | 0.2807(0.1997,0.3697) |
| CoE-ELSC | N<R<T | 0.4024(0.3118,0.4953) |
| CoE-McBASC | N<R<T | 0.5208(0.4194,0.6200) |
| CoE-OMES | N<R<T | 0.3739(0.2931,0.4621) |
| CoE-SCA | N<T<R | 0.2832(0.2182,0.3543) |
| CoE-ZNDAMI | N<R<T | 0.5375(0.4444,0.6290) |
| CoE-composite*** | N<R<T | 0.4655(0.3724,0.5541) |
| TEAO-Spec | R<T<N | 0.3085(0.2406,0.3887) |
| TEAO-Con | T<R<N | 0.6301(0.5347,0.7234) |
| Seq-Entropy | T<R<N | 0.5288(0.4316,0.6315) |
| ETA | T<R<N | 0.4579(0.3663,0.5462) |
| ConSurf (Continuous) | T<R<N | 0.6583(0.5639,0.7438) |
| ConSurf (Discrete) | T<R<N | 0.6494(0.5585,0.7317) |
| Combination | T<R,N | 0.6700(0.5780,0.7563) |

*The outcome order differs for each analysis because the algorithms use different rank orderings; thus, the distribution averages for the three types of experimental groups also occur in different rank orders.

**VUS: Volume under the surface. For a data set with three categories, this measure is analogous to the area under a ROC curve (see Methods). High numbers indicate better discrimination; the maximum value is 1. Numbers in parentheses are the 95% confidence limits of the VUS.

***Composite scores for each LacI position were generated from the mean of the Z-normalized scores of the 5 eigenvector centrality or the 5 pairwise co-evolution analyses.

**Supplementary Table 5.** ROC analysis and true/false class rates at the Youden based optimized operating point along with the corresponding cutoffs for subclasses 1, 2, and 3*.

| Analysis | J** | TC1[‡] | TC2 | TC3 | FC2\|1[‡‡] | FC3\|1 | FC1\|2 | FC3\|2 | FC2\|3 | FC1\|3 | Cutoff1[‡‡‡] | Cutoff2 |
|---|---|---|---|---|---|---|---|---|---|---|---|---|
| EVC-ELSC | 0.2354 | 0.7747 | 0.2394 | 0.4567 | 0.1685 | 0.0568 | 0.4227 | 0.3379 | 0.3544 | 0.1889 | 0.0521 | 0.0596 |
| EVC-McBASC | 0.3266 | 0.7544 | 0.1727 | 0.7261 | 0.1263 | 0.1193 | 0.2869 | 0.5404 | 0.1259 | 0.1480 | 0.0542 | 0.0544 |
| EVC-OMES | 0.1932 | 0.7604 | 0.0412 | 0.5849 | 0.0404 | 0.1992 | 0.4875 | 0.4713 | 0.3761 | 0.0391 | 0.0479 | 0.0556 |
| EVC-SCA | 0.1711 | 0.6226 | 0.3380 | 0.3816 | 0.2543 | 0.1230 | 0.2927 | 0.3693 | 0.4066 | 0.2119 | 0.0543 | 0.0573 |
| EVC-ZNDAMI | 0.2387 | 0.2563 | 0.7675 | 0.4537 | 0.3885 | 0.3553 | 0.0705 | 0.1620 | 0.1019 | 0.4444 | 0.0484 | 0.0587 |
| EVC-comp’ | 0.1938 | 0.7768 | 0.4342 | 0.1766 | 0.2018 | 0.0214 | 0.4206 | 0.1452 | 0.4736 | 0.3497 | -0.2168 | 1.2676 |
| CoE-ELSC | 0.2644 | 0.6704 | 0.0702 | 0.7881 | 0.0641 | 0.2655 | 0.3124 | 0.6174 | 0.1500 | 0.0619 | 62.7344 | 75.6295 |
| CoE-McBASC | 0.3980 | 0.7256 | 0.5508 | 0.5196 | 0.2310 | 0.0434 | 0.2150 | 0.2341 | 0.1467 | 0.3337 | 0.2279 | 0.3835 |
| CoE-OMES | 0.2388 | 0.7581 | 0.2660 | 0.4535 | 0.1606 | 0.0813 | 0.3611 | 0.3729 | 0.2869 | 0.2595 | 6.1450 | 10.3754 |
| CoE-SCA | 0.1371 | 0.5227 | 0.4377 | 0.3137 | 0.2906 | 0.1867 | 0.3822 | 0.1801 | 0.2647 | 0.4216 | 69.1527 | 115.2316 |
| CoE-ZNDAMI | 0.4068 | 0.7729 | 0.3914 | 0.6493 | 0.1662 | 0.0609 | 0.4102 | 0.1984 | 0.1629 | 0.1878 | 3.8871 | 4.9798 |
| CoE-comp | 0.3274 | 0.6853 | 0.1882 | 0.7814 | 0.1360 | 0.1787 | 0.2310 | 0.5808 | 0.1004 | 0.1182 | -0.4718 | 0.0713 |
| TEAO-Spec | 0.2073 | 0.8097 | 0.2898 | 0.3150 | 0.1442 | 0.0461 | 0.5262 | 0.1840 | 0.4011 | 0.2839 | 0.6804 | 0.9822 |
| TEAO-Con | 0.4749 | 0.5534 | 0.6408 | 0.7555 | 0.3600 | 0.0867 | 0.1551 | 0.2041 | 0.0055 | 0.2390 | 0.4091 | 0.9028 |
| Seq-Entropy | 0.3886 | 0.5819 | 0.4086 | 0.7868 | 0.2318 | 0.1863 | 0.1897 | 0.4017 | 0.0532 | 0.1600 | 1.1067 | 2.0445 |
| ETA | 0.3648 | 0.6297 | 0.6060 | 0.4939 | 0.3109 | 0.0594 | 0.1925 | 0.2014 | 0.2532 | 0.2529 | 27.9217 | 73.2643 |
| ConSurf (Continu-ous) | 0.5083 | 0.6323 | 0.5566 | 0.8276 | 0.2794 | 0.0882 | 0.2082 | 0.2351 | 0.0299 | 0.1425 | -1.0008 | 0.1661 |
| ConSurf (Discrete) | 0.5185 | 0.7985 | 0.5421 | 0.6964 | 0.1657 | 0.0357 | 0.2157 | 0.2421 | 0.0619 | 0.2417 | 4.6407 | 7.8375 |
| Combination | 0.4593 | 0.6351 | 0.5518 | 0.7981 | 0.3024 | 0.0625 | 0.2316 | 0.2166 | 0.0348 | 0.1672 | 1.1381 | 2.1954 |

*Substitution outcome classes 1, 2, and 3 are defined in Supplementary Table 4 above. Note that the order differs for each type of analysis because the algorithms use different rank orderings and the distribution averages for the three types of experimental groups occur in different rank orders.
**J is the generalized Youden index (18, 19).
[‡]TC(i): Using the cutoffs listed and given that the truth was class i, what is the probability that a position was correctly classified as i.
[‡‡]FC(i|j): Using the cutoffs listed, and given that the truth was class j, what is the probability that a position was misclassified as i.
[‡‡‡]"Cutoff 1" is the score threshold that best separates class 1 from class 2. "Cutoff2" is the score threshold that best separates class 2 from class 3.

**Supplementary Table 6**. Stepwise logistic regression used to generate the combined analysis.

A. Logistic regression of 14 analyses. (Two eigenvector centrality measures were excluded because their scores highly correlated). Logistic regression was carried out for four groups of positions (single rheostat R1, double rheostat R2, toggle T, and double neutral N) to allow for all possible discriminatory signals. The various analyses identified are indicate in the table with numbers for which a key is listed below. Based on these results, we proceeded with forward elimination that yielded the same sets of analyses for every group separation, regardless of whether we use Wald, LR or Conditional selection (highlighted with green color).

| Group separation | Backward Wald | Backward LR | Backward Conditional | Forward Wald | Forward LR | Forward Conditional |
|---|---|---|---|---|---|---|
| **R1-R2** | 1, 5, 17 | 1, 5, 17 | 1, 5, 17 | 2 | 2 | 2 |
| **R1-T** | 6, 10, 11, 13 | 6, 10, 11, 13 | 6, 10, 11, 13 | 11, 16 | 11,16 | 11,16 |
| **R1-N** | 4, 14, 15 | 4, 14, 15 | 4, 14, 15 | 17 | 17 | 17 |
| **R2-T** | 1, 4, 5, 6, 8, 10, 11, 16 | 1,2,4,5,7,8,10,11,13, 15,16,17 | All except for 12 and 14 | 1,8,11,14,17 | 1,8,11,14,17 | 1,8, 11, 14, 17 |
| **R2-N** | 2,3,5,6,17 | 2,3,5,6,14 | 2,3,5,6,14 | 2,14 | 2,14 | 2,14 |
| **T-N** | 4,5,6,9,13,14,15 | 4,5,6,9,13,14,15 | 4,5,6,9,13,14,15 | 17 | 17 | 17 |

B. Logistic regression after merging the R1 and R2 groups into the "R" category.

| Group separation | Backward Wald | Backward LR | Backward Conditional | Forward Wald | Forward LR | Forward Conditional |
|---|---|---|---|---|---|---|
| **R-T** | 6,8,10,11,13,14 | 6,8,10,11,13,14 | 6,8,10,11,13,14 | 11,16 | 11,16 | 11,16 |
| **R-N** | 2,14 | 2,14 | 2,14 | 17 | 17 | 17 |
| **T-N** | 4,5,6,9,13,14,15 | 4,5,6,9,13,14,15 | 4,5,6,9,13,14,15 | 17 | 17 | 17 |

Numbering key

1: Eigenvector centrality ("EVC")-ELSC; 2: EVC-McBASC; 3: EVC-OMES; 4: EVC-SCA; 5: EVC-ZNDAMI; 6: EVC-composite;
7: CoEvolution ("CoE")-ELSC; 8: CoE-McBASC; 9: CoE-OMES; 10: CoE-SCA; 11: CoE-ZNDAMI; 12: CoE-composite;
13: TEAO-Spec; 14: TEAO-Con; 15: Seq-Entropy; 16: ETA; 17: ConSurf (Continuous)

**Supplementary Figure 12.** ROC curves for Consurf scores (top) and the combination scores derived from combining seven analyses (bottom) for discriminating the rheostat, toggle, and double neutral substitution outcomes in LacI. Note that scores for rheostat and toggle positions were poorly discriminated. The AUC for each ROC is listed below.

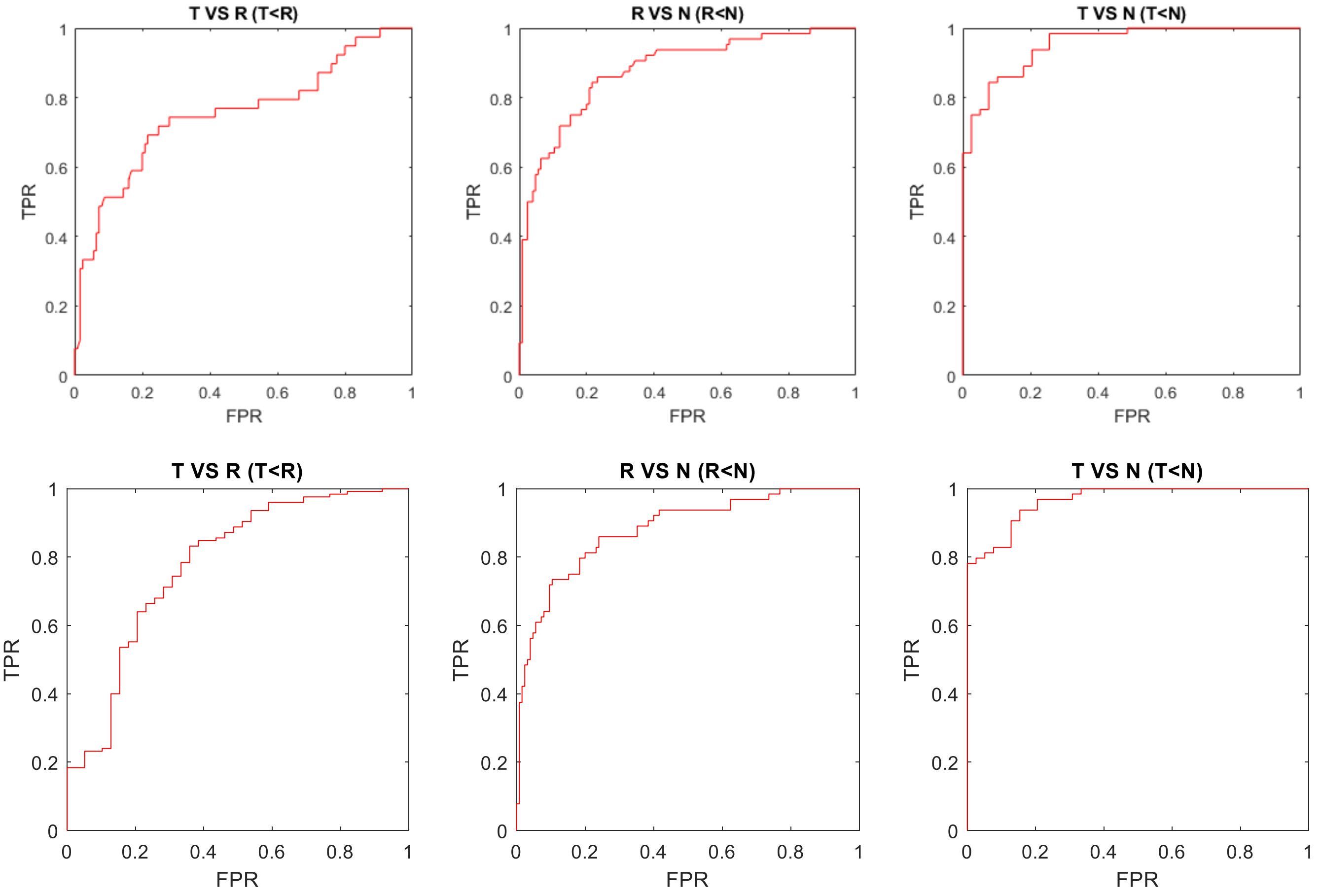


| Pairwise ROCs | AUC (95% CI) for T VS R | AUC (95% CI) for T VS R | AUC (95% CI) for T VS N |
|---|---|---|---|
| ConSurf | 0.7537(0.6555-0.8520) | 0.8791(0.8269-0.9312) | 0.9539(0.9188-0.9891) |
| Combined | 0.7797(0.6920-0.8674) | 0.8804(0.8288-0.9319) | 0.9663(0.9382-0.9945) |

**Supplementary Figure 13**. Both ConSurf and the combination method (derived from combining 7 analyses) show some discrimination of double neutral ("N") and non-neutral ("non-N") positions, especially at the highest score values. (Non-neutral positions comprise all rheostat and toggle positions.) Note that the difference in the two groups appears larger for the combination method than for ConSurf. This difference is quantified in Figure 6 of the main text

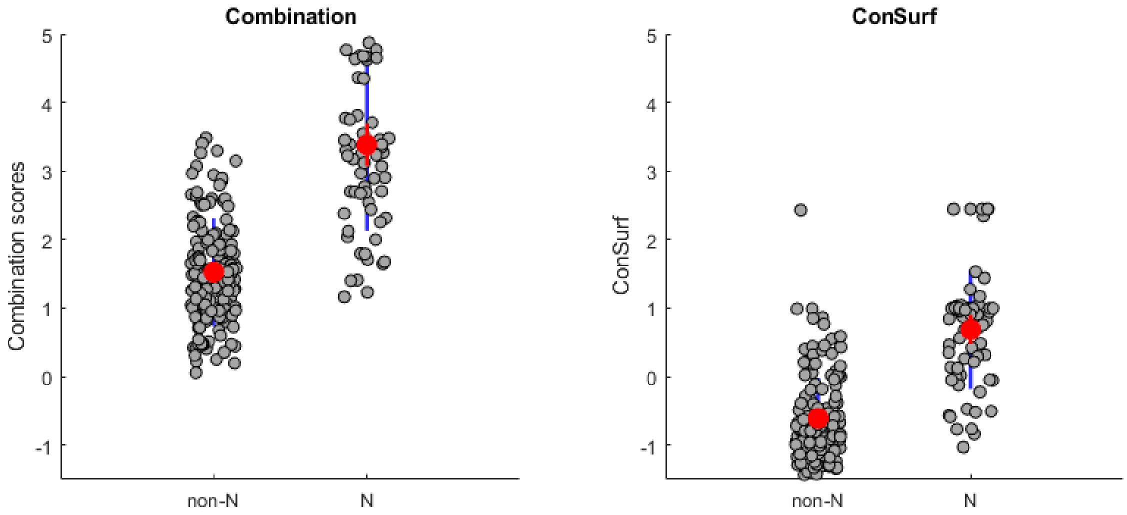

**Supplementary Figure 14.** Correlation matrix of scores from the seven sequence analyses selected to create the combination score (calculated with Equation 1 in main text). The histograms on the diagonal show the distributions of scores for individual methods. Some analyses rank positions with high to low scores; other analyses invert the ranking. Pearson correlation coefficients are shown for each plot; the red line is the result of linear regression. Supplementary Figure 15 shows a heatmap of all correlation coefficients for all pairs of all seventeen sequence analyses used in this study. On the plots below, the scatter and low correlation between many pairs of scores indicates that various analyses each highlight different sets of positions as "important".

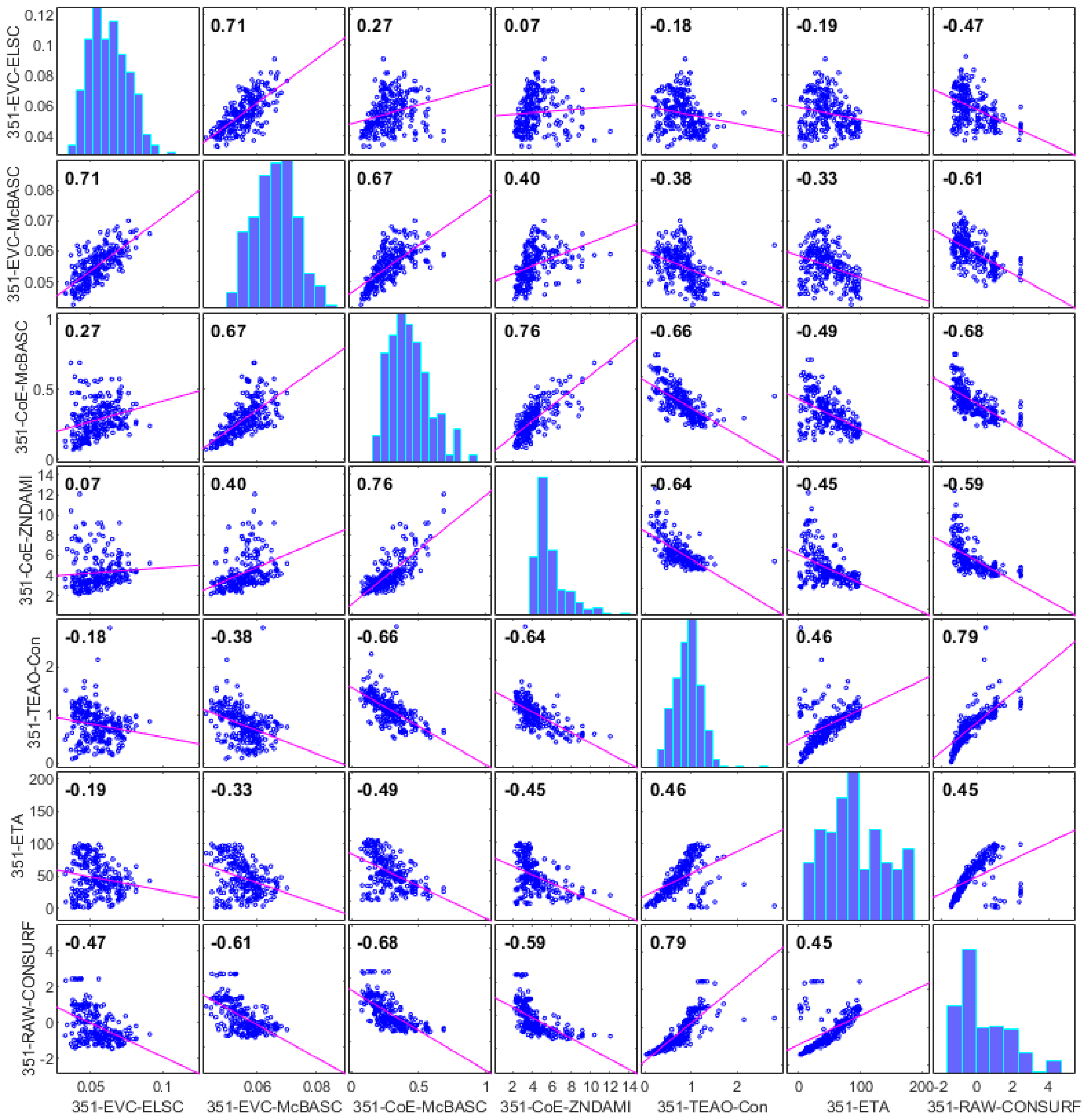

**Supplementary Figure 15**. Heatmap of the Pearson correlation coefficients for all seventeen sequence analyses. Example of the correlation plots are shown above in Supplementary Figure 14. On the heatmap below, the main types of analyses are indicated with dashed boxes. See Methods in the main text for more detail. Analyses within each method class generally show better agreement than do analyses between method classes.

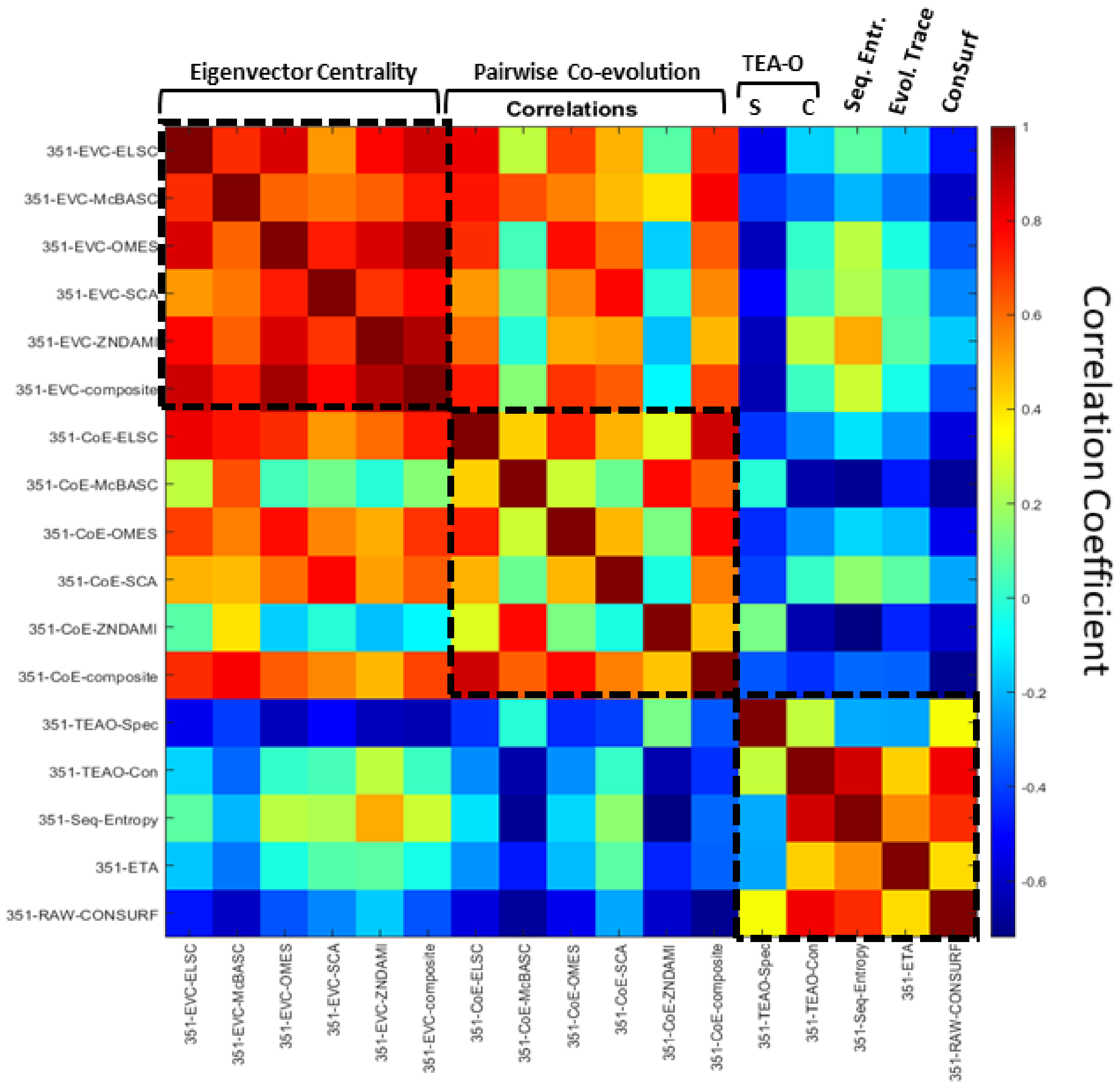

**Supplementary Table 1.** RheoScale neutral, rheostat, and toggle scores calculated for each LacI position using the Miller lab data. The thresholds used assign each position's dominant substitution outcome (or as "unclassified") are described in Methods. The dominant substitution outcome for each position is in Supplementary Lists 1.

n = number of substitutions available, including the wild-type amino acid

"FALSE" = too few substitutions for RheoScale to calculate a score (usually because the substitution abolished repression and induction could not be measured).

**Repression Phenotype**

| Position | n | Neutral | Weighted Rheostat | Toggle |
|---|---|---|---|---|
| 2 | 13 | 0.92 | 0.50 | 0.00 |
| 3 | 13 | 0.75 | 1.00 | 0.08 |
| 4 | 14 | 0.69 | 0.83 | 0.00 |
| 5 | 14 | 0.00 | 1.00 | 0.77 |
| 6 | 13 | 0.25 | 0.67 | 0.42 |
| 7 | 13 | 0.00 | 1.00 | 0.08 |
| 8 | 14 | 0.08 | 1.00 | 0.54 |
| 9 | 14 | 0.00 | 1.00 | 0.85 |
| 10 | 13 | 0.08 | 0.67 | 0.83 |
| 11 | 13 | 0.92 | 0.33 | 0.08 |
| 12 | 13 | 0.92 | 0.33 | 0.08 |
| 13 | 13 | 0.08 | 0.67 | 0.75 |
| 14 | 13 | 0.50 | 1.00 | 0.17 |
| 15 | 14 | 0.23 | 1.00 | 0.62 |
| 16 | 13 | 0.00 | 0.67 | 0.83 |
| 17 | 13 | 0.00 | 0.67 | 0.08 |
| 18 | 13 | 0.00 | 1.00 | 0.75 |
| 19 | 14 | 0.00 | 0.67 | 0.92 |
| 20 | 14 | 0.00 | 0.67 | 0.85 |
| 21 | 13 | 0.00 | 0.67 | 0.92 |
| 22 | 13 | 0.00 | 0.33 | 1.00 |
| 23 | 14 | 0.00 | 0.67 | 0.85 |
| 24 | 14 | 0.69 | 1.00 | 0.15 |
| 25 | 14 | 0.08 | 1.00 | 0.31 |
| 26 | 13 | 0.92 | 0.33 | 0.08 |
| 27 | 13 | 0.92 | 0.50 | 0.00 |
| 28 | 13 | 1.00 | 0.17 | 0.00 |
| 29 | 13 | 0.42 | 1.00 | 0.25 |
| 30 | 14 | 0.08 | 0.67 | 0.77 |
| 31 | 13 | 0.42 | 0.67 | 0.50 |
| 32 | 13 | 1.00 | 0.17 | 0.00 |
| 33 | 13 | 0.25 | 1.00 | 0.08 |
| 34 | 14 | 0.15 | 1.00 | 0.54 |
| 35 | 13 | 0.33 | 1.00 | 0.17 |

**Induction Phenotype**

| Position | n | Neutral | Weighted Rheostat | Toggle |
|---|---|---|---|---|
| 2 | 13 | 1.00 | 0.17 | 0.00 |
| 3 | 13 | 0.83 | 0.33 | 0.17 |
| 4 | 14 | 0.62 | 0.67 | 0.23 |
| 5 | FALSE | | | |
| 6 | 8 | 1.00 | 0.17 | 0.00 |
| 7 | 12 | 1.00 | 0.17 | 0.00 |
| 8 | 7 | 1.00 | 0.17 | 0.00 |
| 9 | FALSE | | | |
| 10 | FALSE | | | |
| 11 | 12 | 1.00 | 0.17 | 0.00 |
| 12 | 12 | 1.00 | 0.17 | 0.00 |
| 13 | FALSE | | | |
| 14 | 11 | 1.00 | 0.17 | 0.00 |
| 15 | 6 | 1.00 | 0.17 | 0.00 |
| 16 | FALSE | | | |
| 17 | 12 | 1.00 | 0.17 | 0.00 |
| 18 | FALSE | | | |
| 19 | FALSE | | | |
| 20 | FALSE | | | |
| 21 | FALSE | | | |
| 22 | FALSE | | | |
| 23 | FALSE | | | |
| 24 | 12 | 0.82 | 0.50 | 0.00 |
| 25 | 10 | 1.00 | 0.17 | 0.00 |
| 26 | 12 | 1.00 | 0.17 | 0.00 |
| 27 | 13 | 1.00 | 0.17 | 0.00 |
| 28 | 13 | 0.83 | 0.50 | 0.00 |
| 29 | 10 | 1.00 | 0.17 | 0.00 |
| 30 | FALSE | | | |
| 31 | 7 | 1.00 | 0.17 | 0.00 |
| 32 | 13 | 1.00 | 0.17 | 0.00 |
| 33 | 12 | 1.00 | 0.17 | 0.00 |
| 34 | 7 | 1.00 | 0.17 | 0.00 |
| 35 | 11 | 1.00 | 0.17 | 0.00 |

**Repression Phenotype**

| Position | n | Neutral | Weighted Rheostat | Toggle |
|---|---|---|---|---|
| 36 | 13 | 0.83 | 0.50 | 0.00 |
| 37 | 13 | 0.50 | 0.67 | 0.08 |
| 38 | 14 | 0.00 | 0.33 | 1.00 |
| 39 | 13 | 0.92 | 0.33 | 0.08 |
| 40 | 13 | 0.92 | 0.33 | 0.08 |
| 41 | 13 | 0.00 | 1.00 | 0.58 |
| 42 | 14 | 0.31 | 1.00 | 0.54 |
| 43 | 13 | 0.92 | 0.33 | 0.08 |
| 44 | 13 | 0.92 | 0.33 | 0.08 |
| 45 | 13 | 0.08 | 1.00 | 0.58 |
| 46 | 14 | 0.92 | 0.33 | 0.08 |
| 47 | 13 | 0.00 | 0.67 | 0.92 |
| 48 | 14 | 0.00 | 1.00 | 0.69 |
| 49 | 13 | 0.08 | 0.67 | 0.75 |
| 50 | 14 | 0.15 | 0.67 | 0.62 |
| 51 | 13 | 0.17 | 1.00 | 0.08 |
| 52 | 14 | 0.69 | 0.83 | 0.00 |
| 53 | 13 | 0.00 | 0.33 | 1.00 |
| 54 | 13 | 0.00 | 0.67 | 0.92 |
| 55 | 13 | 0.33 | 1.00 | 0.17 |
| 56 | 13 | 0.00 | 0.33 | 1.00 |
| 57 | 13 | 0.00 | 1.00 | 0.75 |
| 58 | 13 | 0.08 | 1.00 | 0.50 |
| 59 | 13 | 0.08 | 1.00 | 0.08 |
| 60 | 13 | 0.75 | 0.83 | 0.00 |
| 61 | 13 | 0.92 | 0.50 | 0.00 |
| 62 | 13 | 1.00 | 0.17 | 0.00 |
| 63 | 13 | 1.00 | 0.17 | 0.00 |
| 64 | 14 | 0.38 | 0.33 | 0.62 |
| 65 | 13 | 0.00 | 0.33 | 1.00 |
| 66 | 14 | 0.46 | 0.33 | 0.54 |
| 67 | 13 | 0.75 | 0.67 | 0.17 |
| 68 | 14 | 0.38 | 1.00 | 0.15 |
| 69 | 13 | 1.00 | 0.17 | 0.00 |
| 70 | 13 | 0.75 | 0.50 | 0.00 |
| 71 | 13 | 0.17 | 0.67 | 0.50 |
| 72 | 13 | 0.83 | 0.83 | 0.00 |
| 73 | 13 | 0.92 | 0.50 | 0.00 |
| 74 | 13 | 1.00 | 0.17 | 0.00 |
| 75 | 13 | 1.00 | 0.17 | 0.00 |
| 76 | 13 | 0.42 | 1.00 | 0.08 |
| 77 | 13 | 0.25 | 0.67 | 0.08 |
| 78 | 13 | 0.58 | 0.83 | 0.00 |

**Induction Phenotype**

| Position | n | Neutral | Weighted Rheostat | Toggle |
|---|---|---|---|---|
| 36 | 13 | 1.00 | 0.17 | 0.00 |
| 37 | 12 | 1.00 | 0.17 | 0.00 |
| 38 | FALSE | | | |
| 39 | 12 | 1.00 | 0.17 | 0.00 |
| 40 | 12 | 1.00 | 0.17 | 0.00 |
| 41 | 6 | 1.00 | 0.17 | 0.00 |
| 42 | 7 | 1.00 | 0.17 | 0.00 |
| 43 | 12 | 1.00 | 0.17 | 0.00 |
| 44 | 12 | 1.00 | 0.17 | 0.00 |
| 45 | 6 | 1.00 | 0.17 | 0.00 |
| 46 | 13 | 1.00 | 0.17 | 0.00 |
| 47 | FALSE | | | |
| 48 | 5 | 1.00 | 0.17 | 0.00 |
| 49 | FALSE | | | |
| 50 | 6 | 1.00 | 0.17 | 0.00 |
| 51 | 12 | 1.00 | 0.17 | 0.00 |
| 52 | 14 | 0.62 | 0.67 | 0.23 |
| 53 | FALSE | | | |
| 54 | FALSE | | | |
| 55 | 11 | 1.00 | 0.17 | 0.00 |
| 56 | FALSE | | | |
| 57 | FALSE | | | |
| 58 | 7 | 1.00 | 0.17 | 0.00 |
| 59 | 12 | 1.00 | 0.17 | 0.00 |
| 60 | 13 | 1.00 | 0.17 | 0.00 |
| 61 | 13 | 0.67 | 0.67 | 0.25 |
| 62 | 13 | 1.00 | 0.17 | 0.00 |
| 63 | 13 | 0.83 | 0.50 | 0.00 |
| 64 | 7 | 0.83 | 0.50 | 0.00 |
| 65 | FALSE | | | |
| 66 | 7 | 0.50 | 0.67 | 0.33 |
| 67 | 11 | 0.60 | 0.50 | 0.00 |
| 68 | 12 | 0.73 | 0.67 | 0.09 |
| 69 | 13 | 0.42 | 0.83 | 0.00 |
| 70 | 13 | 0.92 | 0.50 | 0.00 |
| 71 | 7 | 0.83 | 0.50 | 0.00 |
| 72 | 13 | 0.83 | 0.67 | 0.08 |
| 73 | 13 | 0.33 | 0.67 | 0.58 |
| 74 | 13 | 0.67 | 0.67 | 0.25 |
| 75 | 13 | 0.00 | 0.33 | 1.00 |
| 76 | 12 | 0.36 | 0.67 | 0.09 |
| 77 | 12 | 0.91 | 0.50 | 0.00 |
| 78 | 13 | 0.42 | 0.67 | 0.33 |

**Repression Phenotype**

| Position | n | Neutral | Weighted Rheostat | Toggle |
|---|---|---|---|---|
| 79 | 14 | 0.46 | 0.83 | 0.00 |
| 80 | 14 | 0.85 | 0.50 | 0.00 |
| 81 | 13 | 0.42 | 1.00 | 0.08 |
| 82 | 13 | 0.58 | 0.50 | 0.00 |
| 83 | 14 | 0.46 | 0.67 | 0.46 |
| 84 | 13 | 0.75 | 0.83 | 0.00 |
| 85 | 13 | 0.75 | 0.50 | 0.00 |
| 86 | 13 | 0.83 | 0.50 | 0.00 |
| 87 | 13 | 0.33 | 1.00 | 0.42 |
| 88 | 14 | 1.00 | 0.17 | 0.00 |
| 89 | 13 | 1.00 | 0.17 | 0.00 |
| 90 | 13 | 0.92 | 0.50 | 0.00 |
| 91 | 13 | 1.00 | 0.17 | 0.00 |
| 92 | 13 | 0.83 | 0.50 | 0.00 |
| 93 | 13 | 1.00 | 0.17 | 0.00 |
| 94 | 14 | 0.85 | 0.67 | 0.08 |
| 95 | 14 | 0.92 | 0.50 | 0.00 |
| 96 | 14 | 1.00 | 0.17 | 0.00 |
| 97 | 13 | 0.92 | 0.50 | 0.00 |
| 98 | 14 | 0.38 | 1.00 | 0.08 |
| 99 | 14 | 0.62 | 0.83 | 0.00 |
| 100 | 13 | 1.00 | 0.17 | 0.00 |
| 101 | 13 | 1.00 | 0.17 | 0.00 |
| 102 | 13 | 0.92 | 0.50 | 0.00 |
| 103 | 13 | 1.00 | 0.17 | 0.00 |
| 104 | 14 | 1.00 | 0.17 | 0.00 |
| 105 | 13 | 1.00 | 0.17 | 0.00 |
| 106 | 13 | 1.00 | 0.17 | 0.00 |
| 107 | 13 | 1.00 | 0.17 | 0.00 |
| 108 | 13 | 1.00 | 0.17 | 0.00 |
| 109 | 13 | 1.00 | 0.17 | 0.00 |
| 110 | 13 | 0.92 | 0.50 | 0.00 |
| 111 | 14 | 1.00 | 0.17 | 0.00 |
| 112 | 13 | 1.00 | 0.17 | 0.00 |
| 113 | 14 | 0.92 | 0.33 | 0.08 |
| 114 | 13 | 0.58 | 0.67 | 0.33 |
| 115 | 13 | 1.00 | 0.17 | 0.00 |
| 116 | 13 | 0.58 | 0.50 | 0.00 |
| 117 | 13 | 0.67 | 0.83 | 0.00 |
| 118 | 13 | 0.00 | 1.00 | 0.42 |
| 119 | 14 | 0.54 | 0.67 | 0.23 |
| 120 | 13 | 0.92 | 0.50 | 0.00 |
| 121 | 13 | 0.83 | 0.67 | 0.08 |

**Induction Phenotype**

| Position | n | Neutral | Weighted Rheostat | Toggle |
|---|---|---|---|---|
| 79 | 14 | 0.46 | 0.67 | 0.08 |
| 80 | 14 | 0.54 | 0.67 | 0.31 |
| 81 | 12 | 0.82 | 0.50 | 0.00 |
| 82 | 13 | 0.75 | 0.50 | 0.00 |
| 83 | 9 | 0.63 | 0.50 | 0.00 |
| 84 | 13 | 0.08 | 1.00 | 0.58 |
| 85 | 13 | 0.92 | 0.50 | 0.00 |
| 86 | 13 | 0.92 | 0.50 | 0.00 |
| 87 | 12 | 0.36 | 0.50 | 0.00 |
| 88 | 14 | 0.00 | 0.33 | 1.00 |
| 89 | 13 | 0.75 | 0.50 | 0.00 |
| 90 | 13 | 1.00 | 0.17 | 0.00 |
| 91 | 13 | 1.00 | 0.17 | 0.00 |
| 92 | 13 | 0.42 | 0.67 | 0.33 |
| 93 | 13 | 0.92 | 0.50 | 0.00 |
| 94 | 13 | 0.50 | 0.50 | 0.00 |
| 95 | 14 | 0.08 | 0.67 | 0.77 |
| 96 | 14 | 0.08 | 0.67 | 0.69 |
| 97 | 13 | 0.58 | 0.67 | 0.25 |
| 98 | 13 | 0.25 | 0.67 | 0.33 |
| 99 | 14 | 0.62 | 0.67 | 0.31 |
| 100 | 13 | 1.00 | 0.17 | 0.00 |
| 101 | 13 | 1.00 | 0.17 | 0.00 |
| 102 | 13 | 1.00 | 0.17 | 0.00 |
| 103 | 13 | 1.00 | 0.17 | 0.00 |
| 104 | 14 | 0.92 | 0.50 | 0.00 |
| 105 | 13 | 1.00 | 0.17 | 0.00 |
| 106 | 13 | 1.00 | 0.17 | 0.00 |
| 107 | 13 | 0.75 | 0.83 | 0.00 |
| 108 | 13 | 1.00 | 0.17 | 0.00 |
| 109 | 13 | 1.00 | 0.17 | 0.00 |
| 110 | 13 | 0.58 | 0.67 | 0.17 |
| 111 | 14 | 1.00 | 0.17 | 0.00 |
| 112 | 13 | 1.00 | 0.17 | 0.00 |
| 113 | 13 | 0.75 | 0.50 | 0.00 |
| 114 | 9 | 0.75 | 0.83 | 0.00 |
| 115 | 13 | 1.00 | 0.17 | 0.00 |
| 116 | 13 | 1.00 | 0.17 | 0.00 |
| 117 | 13 | 1.00 | 0.17 | 0.00 |
| 118 | 8 | 1.00 | 0.17 | 0.00 |
| 119 | 12 | 0.73 | 0.50 | 0.00 |
| 120 | 13 | 0.92 | 0.50 | 0.00 |
| 121 | 12 | 1.00 | 0.17 | 0.00 |

**Repression Phenotype**

| Position | n | Neutral | Weighted Rheostat | Toggle |
|---|---|---|---|---|
| 122 | 13 | 0.42 | 1.00 | 0.33 |
| 123 | 14 | 0.31 | 0.33 | 0.69 |
| 124 | 14 | 0.38 | 0.67 | 0.54 |
| 125 | 14 | 0.85 | 0.50 | 0.00 |
| 126 | 13 | 1.00 | 0.17 | 0.00 |
| 127 | 13 | 0.83 | 0.83 | 0.00 |
| 128 | 13 | 0.42 | 1.00 | 0.33 |
| 129 | 14 | 1.00 | 0.17 | 0.00 |
| 130 | 14 | 1.00 | 0.17 | 0.00 |
| 131 | 13 | 1.00 | 0.17 | 0.00 |
| 132 | 14 | 1.00 | 0.17 | 0.00 |
| 133 | 13 | 0.83 | 0.67 | 0.08 |
| 134 | 14 | 1.00 | 0.17 | 0.00 |
| 135 | 13 | 1.00 | 0.17 | 0.00 |
| 136 | 14 | 0.62 | 0.67 | 0.23 |
| 137 | 13 | 0.92 | 0.50 | 0.00 |
| 138 | 13 | 1.00 | 0.17 | 0.00 |
| 139 | 13 | 1.00 | 0.17 | 0.00 |
| 140 | 13 | 1.00 | 0.17 | 0.00 |
| 141 | 14 | 1.00 | 0.17 | 0.00 |
| 142 | 14 | 1.00 | 0.17 | 0.00 |
| 143 | 14 | 1.00 | 0.17 | 0.00 |
| 144 | 13 | 1.00 | 0.17 | 0.00 |
| 145 | 13 | 0.92 | 0.50 | 0.00 |
| 146 | 13 | 0.50 | 0.33 | 0.50 |
| 147 | 13 | 0.33 | 0.67 | 0.58 |
| 148 | 12 | 0.36 | 1.00 | 0.27 |
| 149 | 14 | 0.85 | 0.33 | 0.15 |
| 150 | 14 | 0.62 | 0.67 | 0.31 |
| 151 | 13 | 1.00 | 0.17 | 0.00 |
| 152 | 14 | 1.00 | 0.17 | 0.00 |
| 153 | 13 | 1.00 | 0.17 | 0.00 |
| 154 | 14 | 1.00 | 0.17 | 0.00 |
| 155 | 13 | 1.00 | 0.17 | 0.00 |
| 156 | 14 | 0.92 | 0.33 | 0.08 |
| 157 | 14 | 0.85 | 0.33 | 0.15 |
| 158 | 13 | 1.00 | 0.17 | 0.00 |
| 159 | 14 | 0.85 | 0.67 | 0.08 |
| 160 | 14 | 0.92 | 0.50 | 0.00 |
| 161 | 13 | 0.17 | 0.67 | 0.67 |
| 162 | 13 | 1.00 | 0.17 | 0.00 |
| 163 | 13 | 1.00 | 0.17 | 0.00 |
| 164 | 13 | 1.00 | 0.17 | 0.00 |

**Induction Phenotype**

| Position | n | Neutral | Weighted Rheostat | Toggle |
|---|---|---|---|---|
| 122 | 9 | 1.00 | 0.17 | 0.00 |
| 123 | 5 | 1.00 | 0.17 | 0.00 |
| 124 | 7 | 1.00 | 0.17 | 0.00 |
| 125 | 14 | 0.15 | 0.67 | 0.31 |
| 126 | 13 | 1.00 | 0.17 | 0.00 |
| 127 | 13 | 0.25 | 0.67 | 0.33 |
| 128 | 9 | 0.50 | 0.50 | 0.00 |
| 129 | 14 | 0.85 | 0.50 | 0.00 |
| 130 | 14 | 1.00 | 0.17 | 0.00 |
| 131 | 13 | 1.00 | 0.17 | 0.00 |
| 132 | 14 | 1.00 | 0.17 | 0.00 |
| 133 | 12 | 1.00 | 0.17 | 0.00 |
| 134 | 14 | 1.00 | 0.17 | 0.00 |
| 135 | 13 | 1.00 | 0.17 | 0.00 |
| 136 | 13 | 0.58 | 0.83 | 0.00 |
| 137 | 13 | 1.00 | 0.17 | 0.00 |
| 138 | 13 | 1.00 | 0.17 | 0.00 |
| 139 | 13 | 0.50 | 0.83 | 0.00 |
| 140 | 13 | 0.92 | 0.50 | 0.00 |
| 141 | 14 | 0.92 | 0.50 | 0.00 |
| 142 | 14 | 0.69 | 0.67 | 0.08 |
| 143 | 14 | 0.92 | 0.50 | 0.00 |
| 144 | 13 | 1.00 | 0.17 | 0.00 |
| 145 | 13 | 0.92 | 0.50 | 0.00 |
| 146 | 7 | 1.00 | 0.17 | 0.00 |
| 147 | 6 | 1.00 | 0.17 | 0.00 |
| 148 | 10 | 1.00 | 0.17 | 0.00 |
| 149 | 13 | 0.08 | 0.67 | 0.50 |
| 150 | 12 | 0.36 | 0.67 | 0.09 |
| 151 | 13 | 0.92 | 0.50 | 0.00 |
| 152 | 14 | 0.92 | 0.50 | 0.00 |
| 153 | 13 | 1.00 | 0.17 | 0.00 |
| 154 | 14 | 0.85 | 0.50 | 0.00 |
| 155 | 13 | 1.00 | 0.17 | 0.00 |
| 156 | 13 | 0.92 | 0.50 | 0.00 |
| 157 | 12 | 1.00 | 0.17 | 0.00 |
| 158 | 13 | 1.00 | 0.17 | 0.00 |
| 159 | 13 | 0.33 | 0.67 | 0.17 |
| 160 | 14 | 0.69 | 0.67 | 0.15 |
| 161 | 7 | 0.13 | 0.67 | 0.38 |
| 162 | 13 | 1.00 | 0.17 | 0.00 |
| 163 | 13 | 0.83 | 0.50 | 0.00 |
| 164 | 13 | 1.00 | 0.17 | 0.00 |

**Repression Phenotype**

| Position | n | Neutral | Weighted Rheostat | Toggle |
|---|---|---|---|---|
| 165 | 14 | 1.00 | 0.17 | 0.00 |
| 166 | 13 | 0.17 | 1.00 | 0.67 |
| 167 | 14 | 1.00 | 0.17 | 0.00 |
| 168 | 13 | 0.92 | 0.50 | 0.00 |
| 169 | 13 | 0.83 | 0.67 | 0.08 |
| 170 | 13 | 0.67 | 0.33 | 0.33 |
| 171 | 13 | 0.42 | 0.67 | 0.50 |
| 172 | 13 | 0.92 | 0.33 | 0.08 |
| 173 | 13 | 0.75 | 0.67 | 0.08 |
| 174 | 13 | 0.33 | 0.67 | 0.42 |
| 175 | 14 | 1.00 | 0.17 | 0.00 |
| 176 | 13 | 1.00 | 0.17 | 0.00 |
| 177 | 13 | 0.92 | 0.50 | 0.00 |
| 178 | 13 | 0.92 | 0.50 | 0.00 |
| 179 | 13 | 0.00 | 0.67 | 0.83 |
| 180 | 13 | 1.00 | 0.17 | 0.00 |
| 181 | 13 | 1.00 | 0.17 | 0.00 |
| 182 | 14 | 0.46 | 1.00 | 0.38 |
| 183 | 13 | 0.50 | 0.67 | 0.33 |
| 184 | 13 | 0.33 | 0.33 | 0.67 |
| 185 | 13 | 0.33 | 0.67 | 0.50 |
| 186 | 13 | 1.00 | 0.17 | 0.00 |
| 187 | 13 | 0.17 | 0.67 | 0.75 |
| 188 | 13 | 0.83 | 0.67 | 0.08 |
| 189 | 13 | 1.00 | 0.17 | 0.00 |
| 190 | 13 | 1.00 | 0.17 | 0.00 |
| 191 | 13 | 0.67 | 1.00 | 0.17 |
| 192 | 14 | 1.00 | 0.17 | 0.00 |
| 193 | 13 | 1.00 | 0.17 | 0.00 |
| 194 | 13 | 0.67 | 0.67 | 0.25 |
| 195 | 13 | 0.92 | 0.50 | 0.00 |
| 196 | 13 | 0.92 | 0.50 | 0.00 |
| 197 | 13 | 0.83 | 0.50 | 0.00 |
| 198 | 13 | 0.92 | 0.50 | 0.00 |
| 199 | 13 | 1.00 | 0.17 | 0.00 |
| 200 | 13 | 0.92 | 0.33 | 0.08 |
| 201 | 14 | 0.23 | 0.33 | 0.77 |
| 202 | 13 | 0.92 | 0.50 | 0.00 |
| 203 | 13 | 0.92 | 0.50 | 0.00 |
| 204 | 13 | 1.00 | 0.17 | 0.00 |
| 205 | 13 | 0.50 | 0.33 | 0.50 |
| 206 | 14 | 1.00 | 0.17 | 0.00 |
| 207 | 13 | 1.00 | 0.17 | 0.00 |

**Induction Phenotype**

| Position | n | Neutral | Weighted Rheostat | Toggle |
|---|---|---|---|---|
| 165 | 14 | 1.00 | 0.17 | 0.00 |
| 166 | 7 | 0.67 | 0.50 | 0.00 |
| 167 | 14 | 1.00 | 0.17 | 0.00 |
| 168 | 13 | 1.00 | 0.17 | 0.00 |
| 169 | 12 | 0.91 | 0.50 | 0.00 |
| 170 | 9 | 0.75 | 0.50 | 0.00 |
| 171 | 8 | 1.00 | 0.17 | 0.00 |
| 172 | 12 | 1.00 | 0.17 | 0.00 |
| 173 | 13 | 0.92 | 0.50 | 0.00 |
| 174 | 8 | 0.86 | 0.33 | 0.14 |
| 175 | 14 | 1.00 | 0.17 | 0.00 |
| 176 | 13 | 1.00 | 0.17 | 0.00 |
| 177 | 13 | 1.00 | 0.17 | 0.00 |
| 178 | 13 | 1.00 | 0.17 | 0.00 |
| 179 | FALSE | | | |
| 180 | 13 | 1.00 | 0.17 | 0.00 |
| 181 | 13 | 1.00 | 0.17 | 0.00 |
| 182 | 9 | 1.00 | 0.17 | 0.00 |
| 183 | 9 | 0.88 | 0.33 | 0.13 |
| 184 | 5 | 1.00 | 0.17 | 0.00 |
| 185 | 7 | 1.00 | 0.17 | 0.00 |
| 186 | 13 | 1.00 | 0.17 | 0.00 |
| 187 | 5 | 0.00 | 0.67 | 0.75 |
| 188 | 12 | 0.73 | 0.83 | 0.00 |
| 189 | 13 | 1.00 | 0.17 | 0.00 |
| 190 | 13 | 1.00 | 0.17 | 0.00 |
| 191 | 13 | 0.00 | 0.67 | 0.92 |
| 192 | 14 | 0.08 | 1.00 | 0.38 |
| 193 | 13 | 0.00 | 0.33 | 1.00 |
| 194 | 10 | 0.44 | 0.67 | 0.44 |
| 195 | 13 | 0.33 | 0.67 | 0.08 |
| 196 | 13 | 0.83 | 0.67 | 0.08 |
| 197 | 13 | 0.00 | 0.67 | 0.83 |
| 198 | 13 | 0.92 | 0.50 | 0.00 |
| 199 | 13 | 1.00 | 0.17 | 0.00 |
| 200 | 12 | 0.27 | 1.00 | 0.27 |
| 201 | FALSE | | | |
| 202 | 13 | 1.00 | 0.17 | 0.00 |
| 203 | 13 | 1.00 | 0.17 | 0.00 |
| 204 | 13 | 1.00 | 0.17 | 0.00 |
| 205 | 7 | 1.00 | 0.17 | 0.00 |
| 206 | 14 | 1.00 | 0.17 | 0.00 |
| 207 | 13 | 1.00 | 0.17 | 0.00 |

**Repression Phenotype**

| Position | n | Neutral | Weighted Rheostat | Toggle |
|---|---|---|---|---|
| 208 | 14 | 1.00 | 0.17 | 0.00 |
| 209 | 13 | 1.00 | 0.17 | 0.00 |
| 210 | 14 | 0.92 | 0.50 | 0.00 |
| 211 | 13 | 1.00 | 0.17 | 0.00 |
| 212 | 13 | 1.00 | 0.17 | 0.00 |
| 213 | 14 | 0.77 | 0.33 | 0.23 |
| 214 | 13 | 1.00 | 0.17 | 0.00 |
| 215 | 13 | 1.00 | 0.17 | 0.00 |
| 216 | 13 | 1.00 | 0.17 | 0.00 |
| 217 | 13 | 1.00 | 0.17 | 0.00 |
| 218 | 13 | 0.25 | 0.67 | 0.67 |
| 219 | 14 | 1.00 | 0.17 | 0.00 |
| 220 | 14 | 0.77 | 0.67 | 0.08 |
| 221 | 13 | 0.75 | 0.83 | 0.00 |
| 222 | 13 | 0.33 | 0.67 | 0.58 |
| 223 | 14 | 1.00 | 0.17 | 0.00 |
| 224 | 13 | 0.92 | 0.50 | 0.00 |
| 225 | 13 | 0.08 | 0.67 | 0.83 |
| 226 | 13 | 0.58 | 1.00 | 0.17 |
| 227 | 13 | 0.92 | 0.50 | 0.00 |
| 228 | 13 | 0.92 | 0.50 | 0.00 |
| 229 | 14 | 0.54 | 0.67 | 0.38 |
| 230 | 14 | 0.92 | 0.33 | 0.08 |
| 231 | 13 | 0.92 | 0.50 | 0.00 |
| 232 | 14 | 0.77 | 0.33 | 0.23 |
| 233 | 13 | 0.42 | 0.67 | 0.42 |
| 234 | 14 | 1.00 | 0.17 | 0.00 |
| 235 | 13 | 0.92 | 0.50 | 0.00 |
| 236 | 13 | 1.00 | 0.17 | 0.00 |
| 237 | 14 | 1.00 | 0.17 | 0.00 |
| 238 | 14 | 1.00 | 0.17 | 0.00 |
| 239 | 13 | 0.75 | 0.67 | 0.17 |
| 240 | 14 | 1.00 | 0.17 | 0.00 |
| 241 | 13 | 0.25 | 0.33 | 0.75 |
| 242 | 14 | 0.46 | 0.33 | 0.54 |
| 243 | 13 | 0.08 | 0.67 | 0.75 |
| 244 | 14 | 0.46 | 0.67 | 0.46 |
| 245 | 13 | 0.75 | 0.33 | 0.25 |
| 246 | 14 | 0.62 | 0.67 | 0.15 |
| 247 | 14 | 0.00 | 0.67 | 0.92 |
| 248 | 13 | 1.00 | 0.17 | 0.00 |
| 249 | 14 | 0.38 | 0.67 | 0.54 |
| 250 | 13 | 0.25 | 0.33 | 0.75 |

**Induction Phenotype**

| Position | n | Neutral | Weighted Rheostat | Toggle |
|---|---|---|---|---|
| 208 | 14 | 1.00 | 0.17 | 0.00 |
| 209 | 13 | 1.00 | 0.17 | 0.00 |
| 210 | 14 | 1.00 | 0.17 | 0.00 |
| 211 | 13 | 1.00 | 0.17 | 0.00 |
| 212 | 13 | 1.00 | 0.17 | 0.00 |
| 213 | 11 | 1.00 | 0.17 | 0.00 |
| 214 | 13 | 1.00 | 0.17 | 0.00 |
| 215 | 13 | 1.00 | 0.17 | 0.00 |
| 216 | 13 | 1.00 | 0.17 | 0.00 |
| 217 | 13 | 1.00 | 0.17 | 0.00 |
| 218 | 5 | 1.00 | 0.17 | 0.00 |
| 219 | 14 | 0.92 | 0.50 | 0.00 |
| 220 | 14 | 0.00 | 0.67 | 0.77 |
| 221 | 13 | 1.00 | 0.17 | 0.00 |
| 222 | 6 | 0.80 | 0.50 | 0.00 |
| 223 | 14 | 0.92 | 0.50 | 0.00 |
| 224 | 13 | 1.00 | 0.17 | 0.00 |
| 225 | FALSE | | | |
| 226 | 11 | 1.00 | 0.17 | 0.00 |
| 227 | 13 | 1.00 | 0.17 | 0.00 |
| 228 | 13 | 1.00 | 0.17 | 0.00 |
| 229 | 10 | 0.89 | 0.33 | 0.11 |
| 230 | 13 | 1.00 | 0.17 | 0.00 |
| 231 | 13 | 1.00 | 0.17 | 0.00 |
| 232 | 11 | 0.90 | 0.33 | 0.10 |
| 233 | 8 | 1.00 | 0.17 | 0.00 |
| 234 | 14 | 1.00 | 0.17 | 0.00 |
| 235 | 13 | 1.00 | 0.17 | 0.00 |
| 236 | 13 | 1.00 | 0.17 | 0.00 |
| 237 | 14 | 1.00 | 0.17 | 0.00 |
| 238 | 14 | 1.00 | 0.17 | 0.00 |
| 239 | 11 | 1.00 | 0.17 | 0.00 |
| 240 | 14 | 1.00 | 0.17 | 0.00 |
| 241 | FALSE | | | |
| 242 | 7 | 1.00 | 0.17 | 0.00 |
| 243 | FALSE | | | |
| 244 | 8 | 1.00 | 0.17 | 0.00 |
| 245 | 10 | 0.33 | 0.33 | 0.67 |
| 246 | 12 | 0.09 | 0.67 | 0.45 |
| 247 | FALSE | | | |
| 248 | 13 | 0.08 | 0.33 | 0.92 |
| 249 | 7 | 0.83 | 0.50 | 0.00 |
| 250 | FALSE | | | |

### Repression Phenotype

| Position | n | Neutral | Weighted Rheostat | Toggle |
|---|---|---|---|---|
| 251 | 13 | 0.17 | 0.67 | 0.58 |
| 252 | 13 | 0.08 | 0.33 | 0.92 |
| 253 | 13 | 0.33 | 0.33 | 0.67 |
| 254 | 14 | 0.62 | 0.67 | 0.31 |
| 255 | 13 | 0.42 | 0.67 | 0.08 |
| 256 | 12 | 0.55 | 0.67 | 0.36 |
| 257 | 14 | 0.38 | 1.00 | 0.46 |
| 258 | 14 | 1.00 | 0.17 | 0.00 |
| 259 | 13 | 0.33 | 1.00 | 0.33 |
| 260 | 13 | 1.00 | 0.17 | 0.00 |
| 261 | 13 | 1.00 | 0.17 | 0.00 |
| 262 | 13 | 0.92 | 0.50 | 0.00 |
| 263 | 13 | 0.50 | 0.67 | 0.33 |
| 264 | 14 | 0.46 | 0.67 | 0.46 |
| 265 | 13 | 0.58 | 1.00 | 0.17 |
| 266 | 13 | 1.00 | 0.17 | 0.00 |
| 267 | 14 | 0.62 | 0.67 | 0.15 |
| 268 | 14 | 0.00 | 0.33 | 1.00 |
| 269 | 13 | 0.33 | 0.67 | 0.50 |
| 270 | 14 | 0.31 | 0.33 | 0.69 |
| 271 | 14 | 0.46 | 0.67 | 0.46 |
| 272 | 13 | 0.00 | 0.67 | 0.92 |
| 273 | 13 | 0.58 | 0.67 | 0.33 |
| 274 | 14 | 0.77 | 0.33 | 0.23 |
| 275 | 14 | 0.85 | 0.67 | 0.08 |
| 276 | 14 | 1.00 | 0.17 | 0.00 |
| 277 | 13 | 1.00 | 0.17 | 0.00 |
| 278 | 14 | 0.77 | 0.67 | 0.15 |
| 279 | 13 | 0.25 | 0.33 | 0.75 |
| 280 | 12 | 1.00 | 0.17 | 0.00 |
| 281 | 13 | 0.75 | 0.67 | 0.17 |
| 282 | 13 | 0.17 | 0.33 | 0.83 |
| 283 | 14 | 0.92 | 0.33 | 0.08 |
| 284 | 13 | 0.42 | 1.00 | 0.33 |
| 285 | 13 | 0.42 | 0.67 | 0.42 |
| 286 | 13 | 0.00 | 0.67 | 0.83 |
| 287 | 14 | 0.23 | 0.33 | 0.77 |
| 288 | 14 | 0.31 | 0.33 | 0.69 |
| 289 | 14 | 0.15 | 0.67 | 0.77 |
| 290 | 13 | 0.92 | 0.50 | 0.00 |
| 291 | 13 | 0.67 | 0.50 | 0.00 |
| 292 | 14 | 0.54 | 1.00 | 0.23 |
| 293 | 13 | 1.00 | 0.17 | 0.00 |

### Induction Phenotype

| Position | n | Neutral | Weighted Rheostat | Toggle |
|---|---|---|---|---|
| 251 | 6 | 1.00 | 0.17 | 0.00 |
| 252 | FALSE | | | |
| 253 | 5 | 1.00 | 0.17 | 0.00 |
| 254 | 10 | 1.00 | 0.17 | 0.00 |
| 255 | 12 | 1.00 | 0.17 | 0.00 |
| 256 | 9 | 1.00 | 0.17 | 0.00 |
| 257 | 8 | 1.00 | 0.17 | 0.00 |
| 258 | 14 | 1.00 | 0.17 | 0.00 |
| 259 | 9 | 1.00 | 0.17 | 0.00 |
| 260 | 13 | 1.00 | 0.17 | 0.00 |
| 261 | 13 | 1.00 | 0.17 | 0.00 |
| 262 | 13 | 1.00 | 0.17 | 0.00 |
| 263 | 12 | 0.55 | 0.33 | 0.45 |
| 264 | 8 | 1.00 | 0.17 | 0.00 |
| 265 | 11 | 1.00 | 0.17 | 0.00 |
| 266 | 13 | 1.00 | 0.17 | 0.00 |
| 267 | 12 | 1.00 | 0.17 | 0.00 |
| 268 | FALSE | | | |
| 269 | 7 | 1.00 | 0.17 | 0.00 |
| 270 | 5 | 1.00 | 0.17 | 0.00 |
| 271 | 8 | 0.86 | 0.33 | 0.14 |
| 272 | FALSE | | | |
| 273 | 9 | 0.13 | 0.67 | 0.75 |
| 274 | 11 | 0.10 | 0.33 | 0.90 |
| 275 | 14 | 0.00 | 0.67 | 0.85 |
| 276 | 14 | 0.00 | 0.67 | 0.92 |
| 277 | 13 | 1.00 | 0.17 | 0.00 |
| 278 | 12 | 0.64 | 0.67 | 0.27 |
| 279 | FALSE | | | |
| 280 | 13 | 0.92 | 0.50 | 0.00 |
| 281 | 11 | 1.00 | 0.17 | 0.00 |
| 282 | FALSE | | | |
| 283 | 13 | 0.92 | 0.50 | 0.00 |
| 284 | 9 | 1.00 | 0.17 | 0.00 |
| 285 | 8 | 1.00 | 0.17 | 0.00 |
| 286 | FALSE | | | |
| 287 | FALSE | | | |
| 288 | 5 | 1.00 | 0.17 | 0.00 |
| 289 | 5 | 0.50 | 0.83 | 0.00 |
| 290 | 13 | 0.92 | 0.50 | 0.00 |
| 291 | 13 | 0.83 | 0.50 | 0.00 |
| 292 | 12 | 0.64 | 0.83 | 0.00 |
| 293 | 13 | 0.08 | 0.67 | 0.75 |

**Repression Phenotype**

| Position | n | Neutral | Weighted Rheostat | Toggle |
|---|---|---|---|---|
| 294 | 13 | 1.00 | 0.17 | 0.00 |
| 295 | 13 | 1.00 | 0.17 | 0.00 |
| 296 | 13 | 0.83 | 0.67 | 0.08 |
| 297 | 13 | 0.00 | 0.67 | 0.83 |
| 298 | 13 | 1.00 | 0.17 | 0.00 |
| 299 | 14 | 0.92 | 0.50 | 0.00 |
| 300 | 13 | 0.33 | 0.67 | 0.50 |
| 301 | 14 | 0.46 | 1.00 | 0.38 |
| 302 | 14 | 1.00 | 0.17 | 0.00 |
| 303 | 13 | 0.92 | 0.50 | 0.00 |
| 304 | 13 | 0.67 | 0.67 | 0.17 |
| 305 | 13 | 1.00 | 0.17 | 0.00 |
| 306 | 13 | 1.00 | 0.17 | 0.00 |
| 307 | 13 | 1.00 | 0.17 | 0.00 |
| 308 | 13 | 1.00 | 0.17 | 0.00 |
| 309 | 13 | 1.00 | 0.17 | 0.00 |
| 310 | 13 | 1.00 | 0.17 | 0.00 |
| 311 | 13 | 1.00 | 0.17 | 0.00 |
| 312 | 13 | 1.00 | 0.17 | 0.00 |
| 313 | 14 | 1.00 | 0.17 | 0.00 |
| 314 | 13 | 1.00 | 0.17 | 0.00 |
| 315 | 13 | 1.00 | 0.17 | 0.00 |
| 316 | 14 | 1.00 | 0.17 | 0.00 |
| 317 | 13 | 1.00 | 0.17 | 0.00 |
| 318 | 13 | 1.00 | 0.17 | 0.00 |
| 319 | 13 | 0.58 | 1.00 | 0.08 |
| 320 | 13 | 1.00 | 0.17 | 0.00 |
| 321 | 14 | 0.54 | 1.00 | 0.23 |
| 322 | 13 | 1.00 | 0.17 | 0.00 |
| 323 | 13 | 0.33 | 1.00 | 0.33 |
| 324 | 14 | 0.92 | 0.50 | 0.00 |
| 325 | 13 | 1.00 | 0.17 | 0.00 |
| 326 | 13 | 0.00 | 0.67 | 0.92 |
| 327 | 13 | 1.00 | 0.17 | 0.00 |
| 328 | 14 | 0.15 | 0.67 | 0.77 |
| 329 | 14 | 0.85 | 0.50 | 0.00 |

**Induction Phenotype**

| Position | n | Neutral | Weighted Rheostat | Toggle |
|---|---|---|---|---|
| 294 | 13 | 1.00 | 0.17 | 0.00 |
| 295 | 13 | 0.83 | 0.83 | 0.00 |
| 296 | 12 | 0.55 | 0.67 | 0.27 |
| 297 | 5 | 0.50 | 0.50 | 0.00 |
| 298 | 13 | 0.83 | 0.50 | 0.00 |
| 299 | 14 | 0.92 | 0.50 | 0.00 |
| 300 | 7 | 0.83 | 0.33 | 0.17 |
| 301 | 10 | 0.56 | 0.67 | 0.11 |
| 302 | 14 | 1.00 | 0.17 | 0.00 |
| 303 | 13 | 1.00 | 0.17 | 0.00 |
| 304 | 11 | 0.50 | 0.67 | 0.10 |
| 305 | 13 | 0.83 | 0.50 | 0.00 |
| 306 | 13 | 1.00 | 0.17 | 0.00 |
| 307 | 13 | 1.00 | 0.17 | 0.00 |
| 308 | 13 | 1.00 | 0.17 | 0.00 |
| 309 | 13 | 1.00 | 0.17 | 0.00 |
| 310 | 13 | 1.00 | 0.17 | 0.00 |
| 311 | 13 | 1.00 | 0.17 | 0.00 |
| 312 | 13 | 1.00 | 0.17 | 0.00 |
| 313 | 14 | 1.00 | 0.17 | 0.00 |
| 314 | 13 | 1.00 | 0.17 | 0.00 |
| 315 | 13 | 1.00 | 0.17 | 0.00 |
| 316 | 14 | 1.00 | 0.17 | 0.00 |
| 317 | 13 | 1.00 | 0.17 | 0.00 |
| 318 | 13 | 1.00 | 0.17 | 0.00 |
| 319 | 12 | 0.27 | 1.00 | 0.09 |
| 320 | 13 | 1.00 | 0.17 | 0.00 |
| 321 | 11 | 0.80 | 0.50 | 0.00 |
| 322 | 13 | 1.00 | 0.17 | 0.00 |
| 323 | 9 | 1.00 | 0.17 | 0.00 |
| 324 | 14 | 1.00 | 0.17 | 0.00 |
| 325 | 13 | 1.00 | 0.17 | 0.00 |
| 326 | FALSE | | | |
| 327 | 13 | 1.00 | 0.17 | 0.00 |
| 328 | FALSE | | | |
| 329 | 14 | 0.69 | 0.67 | 0.15 |
